# Nutrient restriction, inducer of yeast meiosis, induces meiotic initiation in mammals

**DOI:** 10.1101/2020.05.15.074542

**Authors:** Xiaoyu Zhang, Sumedha Gunewardena, Ning Wang

## Abstract

From yeasts to mammals, the molecular machinery and chromosome structures carrying out meiosis are frequently conserved. However, the signal to initiate meiosis appears divergent: while nutrient restriction induces meiosis in the yeast system, retinoic acid (RA), a chordate morphogen, is necessary but not sufficient to induce meiotic initiation in mammalian germ cells via its target, *Stra8*. Here, using cultured mouse male germline stem cells without the support of gonadal somatic cells, we show that nutrient restriction in combination with RA robustly induces *Spo11*-dependent meiotic DNA double strand breaks (DSBs) and *Stra8*-dependent meiotic gene programs recapitulating those of early meiosis *in vivo*. Moreover, a distinct network of 11 nutrient restriction-upregulated transcription factor genes was identified, whose expression does not require RA and is associated with early meiosis *in vivo*. Thus, our study proposes a conserved model, in which nutrient restriction induces meiotic initiation by upregulating transcriptional factors for meiotic gene programs, and provides an *in vitro* platform to derive haploid gametes in culture.

**One Sentence Summary:** nutrient restriction synergizes with retinoic acid to induce mammalian meiotic initiation

---

Sexual reproduction depends on the formation of haploid gametes through meiosis, which exchanges replicated parental chromosomes through homologous recombination during meiotic prophase, followed by two rounds of successive divisions to generate haploid gametes carrying novel genetic constitution (*1*). Meiotic initiation requires the activation of an array of meiosis-specific genes that implement the chromosomal events during meiosis prophase, including homologous pairing, synapsis, and recombination (*2*). From yeasts to mammals, meiosis-specific chromosome structures exhibit remarkable evolution conservation, including synapsis formation mediated by the assembly of the synaptonemal complex (SC), homologous recombination through the formation and the subsequent repair of meiotic DNA double-strand breaks (DSBs). Many genes underlying these events are often evolutionarily-related or -conserved. For instances, *Spo11* encodes a DNA topoisomerase-like enzyme that catalyzes meiotic DSB formation. *Dmc1* encodes a meiotic recombinase that repairs DSBs by searching for allelic DNA sequences on the homologous chromatids. *Hormad* genes encode meiosis-specific chromosome factors (Hop1 in yeasts and Hormad1/2 in mammals) that are critical for synapsis and DSB formation and repair.

Despite these evolutionary conservations in meiotic genes and structures, the overarching signal to trigger meiosis appears to be divergent. In yeasts, transition from mitotic to meiotic cell cycles is induced by nutrient restriction. Subsequently, nutrient-sensing pathway induces the expression of *inducer of meiosis 1* (*IME1*), which encodes a single master transcriptional activator for meiotic genes, including *Spo11, Dmc1*, and *Hop1* (*3, 4*). In mammalian germ cells, meiotic initiation requires retinoic acid (RA), an activate metabolite of vitamin A. RA is a morphogen essential for growth and development in chordate animals (*5*). In female oogenesis, RA is synthesized primarily in the mesonephric ducts in embryonic ovaries (*6*). In male spermatogenesis, RA is produced by meiotic and somatic cells in testes (*7*). RA induces meiosis primarily by activating *stimulated by retinoic acid gene 8* (*Stra8*) (*8, 9*). Although STRA8-mediated transcriptional activation of meiotic genes by itself or with other protein has been reported (*10, 11*), neither RA nor STRA8 is sufficient to induce meiosis, suggesting that other signal(s) works with RA signaling to induce mammalian meiotic initiation.

Our recent work in *Stra8*-deficient mice reveals that an autophagy-inducing signal is engaged on meiosis-initiating germ cells (*12*). Interestingly, nutrient restriction, the aforementioned inducer of yeast meiosis, is perhaps the most potent inducer of autophagy (*13*). Thus, we asked whether nutrient restriction might have a conserved role in inducing meiotic initiation in mammalian germ cells. To test this, we established primary culture for male mammalian germline stem (GS) cells by using neonatal mouse testicular cells in C57BL/6 X DBA/2 F1 hybrid background (*14*) (**fig. S1A**; see below for the whole transcriptome profile of the cultured GS cells). Nutrient restriction was applied to cultures by adding 90% Earle’s Balanced Salt Solution (EBSS) to the complete GS culture medium.

Co-treatment of nutrient restriction and RA (referred to as “NRRA”) for 2 days was sufficient to trigger a distinct morphology of cellular enlargement in GS cultures, suggesting cell differentiation (**Fig. 1A; fig. S1B**). NRRA induced a significant activation of essential meiotic genes, including *Spo11, Dmc1*, and *Sycp3* (**Fig. 1B**). *Spo11* and *Dmc1* encodes enzymes that specifically catalyze meiotic DNA double-strand break (DSB) formation and repair (*15, 16*). *Sycp3* encodes a lateral element of synaptonemal complex that forms between two meiotic homologous chromosomes (*17*). Consistently, phosphorylated histone AX (γH2AX) shows that DNA DSBs were most profoundly formed in NRRA-treated culture (**Fig. 1C; fig. S1C**). These effects were not observed in cultures treated with RA or NR alone (**Fig. 1A-C**). Similar effects of NRRA in meiotic gene activation were observed in F9 premeiotic cells (**fig. S2**). Importantly, the meiotic origin of these DSBs was confirmed by DMC1 staining (**Fig. 1D**). In addition, cultured GS cells generated from *Spo11*-deficient mice lack DMC1 foci upon NRRA treatment, constituting genetic evidence that meiotic DSB formation *in vitro* requires *Spo11* (**Fig. 1D;** see below for the transcriptome profiles of *Spo11*-deficient culture). In addition, cultured GS cells exhibited rapid loss of undifferentiation spermatogonia markers, Gfra1 and E-cadherin (**fig. S1D**).

**Fig. 1.**
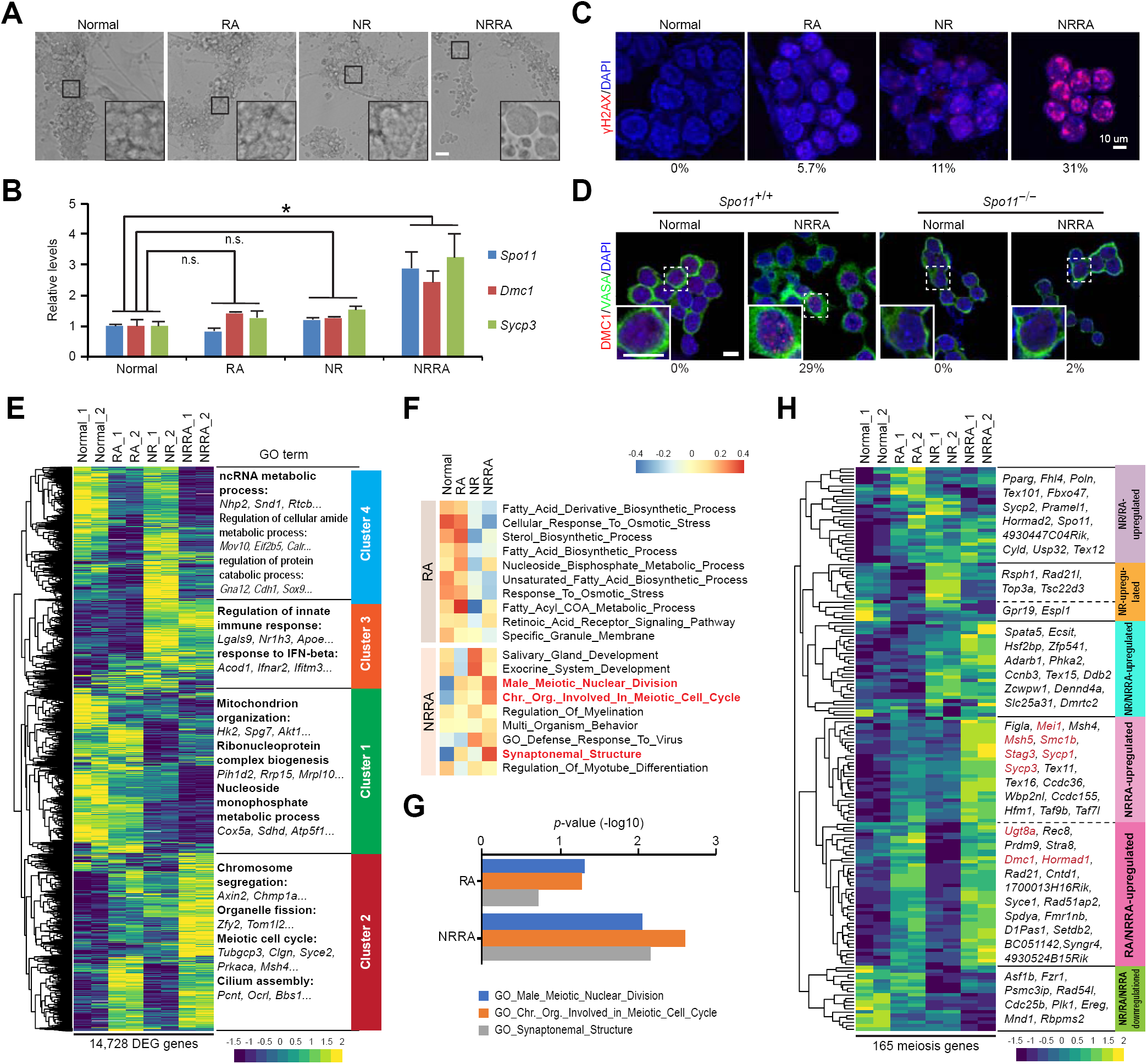
NR is required to induce meiotic initiation with RA in primary GS cell culture. (**A**) Bright-field images of cultured spermatogonia with indicated treatments for 2 days. Scale bars, 50 µm. (**B**) Relative expression of *Spo11, Dmc1*, and *Sycp3* against *Gapdh* analyzed by qRT-PCR in cultured spermatogonia with indicated treatments for 2 days. \**P* < 0.05. n = 3 independent cultures. n.s., no significant. (**C**) Immunostaining for γH2AX (red) and DAPI (blue) with indicated treatment for 2 days. Scale bars, 10 µm. (**D**) Immunostaining for DMC1 (red), DDX4 (green), and DAPI (blue) in *Spo11*^+/+^ and *Spo11*^™/™^ primary spermatogonia culture. Scale bars, 10 µm. (**E**) (Left) UHC and heatmap of gene expression in cultured primary spermatogonia with indicated treatments for 2 days. (Right) Top GO enrichments with representative genes in each cluster. (**F**) GSVA analysis for indicated treatments. In the heatmap, rows are defined by the selected gene sets, and columns by consensus scores for each treatment. Group enriched gene sets are highlighted by different color. (**G**) Bar plot shows *p*-value of GO functions between RA and NRRA treatment. (**H**) UHC and heatmap for the expression of 165 early meiosis-associated genes. STRA8-dependent genes characterized in a previous study in ref. (*2*) are shown in red.

Transcriptomic analysis shows that NRRA induced a gene expression pattern distinct from the treatment of RA or NR alone (**Fig. 1E; fig. S3; table S1**). Four clusters of gene sets were identified by unsupervised hierarchical clustering (UHC) (**table S2**). Notably, genes in Cluster 2, which appear to be genes upregulated by RA and NRRA, are enriched with genes bearing gene ontology (GO) functional terms related to meiosis (**Fig. 1E and fig. S4**). In contrast, Cluster 1, 3, or 4 is enriched with genes for “mitochondrion organization” (Cluster 1, nutrient restriction-downregulated genes), “regulation of innate immune response” (Cluster 3, nutrient restriction-upregulated genes), “ncRNA metabolic process” (Cluster 4, RA-downregulated genes) (**fig. S4**). Consistently, nonparametric and unsupervised gene set variation analysis (GSVA) and hallmark gene set analysis revealed stronger positive correlations of meiosis-related pathways (male meiotic nuclear division, chromosome organization in meiotic cell cycle, synaptonemal structure) in genes upregulated by NRRA than those upregulated by RA (**Fig. 1F and G, fig. S5**) (**table S3**). These data together indicate that nutrient restriction is required to work with RA to induce meiotic initiation *in vitro*.

To examine whether the effect of NRRA in inducing meiotic initiation depends on the genetic background of the cultured GS cells, we established cultured GS cells using testes from neonatal CD1 inbred mice. CD1 cultured GS cells exhibited comparable transcriptomic changes including activation of meiotic genes in response to NRRA treatment (**fig. S6**), suggesting that the sufficient role of NRRA in meiotic initiation is independent of genetic backgrounds.

To dissect the role of nutrient restriction in meiotic gene programs, we assembled a set of 193 meiosis genes by combining two collections of genes that were previously found to be associated with mammalian early meiosis during mouse (*2*) and human meiosis (*18*) spermatogenesis (**fig. S7**). Unsupervised UHC divided 165 differentially expressed genes (DEGs) into five major clusters (**Fig. 1H; table S4**). Nutrient restriction appears to play four roles on the expression of key meiotic genes by: 1) inducing a subset of meiotic gene expression that was not regulated by RA, such as *Rad21l*; 2) initiating the activation of a subset of meiotic genes, which was further enhanced by RA, such as *Zfp541*; 3) synergizing with RA to induce the activation of a subset of meiosis gene expressions, such as *Sycp1, Sycp3, Mei1, Msh5*, and *Stag3*; and 4) augmenting the expression of a subset of meiotic genes induced by RA, including *Dmc1, Smc1b, Ugt8a, Stag3, Hormad1*, as well as *Gm4969*, which encodes MEIOSIN, a transcriptional cofactor for STRA8 required for meiotic prophase program (*11*). Notably, many meiotic genes whose expression depends on STRA8 require nutrient restriction (**Fig. 1H**). Together, these data suggest crucial roles of nutrient restriction by itself or by synergizing with RA in inducing meiotic gene expression; in the absence of nutrient restriction, RA alone is not sufficient (**Fig. 1H**; see the effect of *in vivo* RA treatment on meiotic gene expression despite *Stra8* activation in **fig. S**) (*19*).

To examine NRRA-induced meiotic initiation and its progression potential at the single cell resolution, we isolated single cells from cultured GS cells treated with NRRA for 2 days to induce meiotic initiation, followed by media that support progression for an additional 1 and 2 days. Cultured GS cells treated with normal medium for 4 days were used as control. Single cell RNA-sequencing (scRNA-seq) was then performed using the 10x Genomics platform (**Fig. 2A**). A total of 25,000 cells were sequenced from these samples. Using stringent quality control, 18,088 cells were selected for further analysis (**fig. S9**). All of the cells were pooled to perform clustering analysis, which revealed four major cell clusters based on their distinct gene expression patterns (**Fig. 2B and C**). These four cluster were subsequently annotated by using known marker genes (**fig. S10**). Cluster 0 appears to be undifferentiated spermatogonia (e.g., *Gfra1, Evt5*). Clusters 1, 2, and 3 appear to be differentiating spermatogonia/pre- and meiotic spermatocytes at progressively advanced meiotic stages with upregulated expression of spermatogonial differentiation (e.g., *Sohlh1*) and meiotic genes (e.g., *Dmc1*). Cluster 4 is mostly feeder cells (mouse embryonic fibroblast or MEF) due to the expression of fibroblast genes (*s100a4*). Progressive upregulation of meiotic genes and downregulation of undifferentiated spermatogonia genes on each time points were confirmed by qRT-PCR analysis (**fig. S11**).

**Fig. 2.**
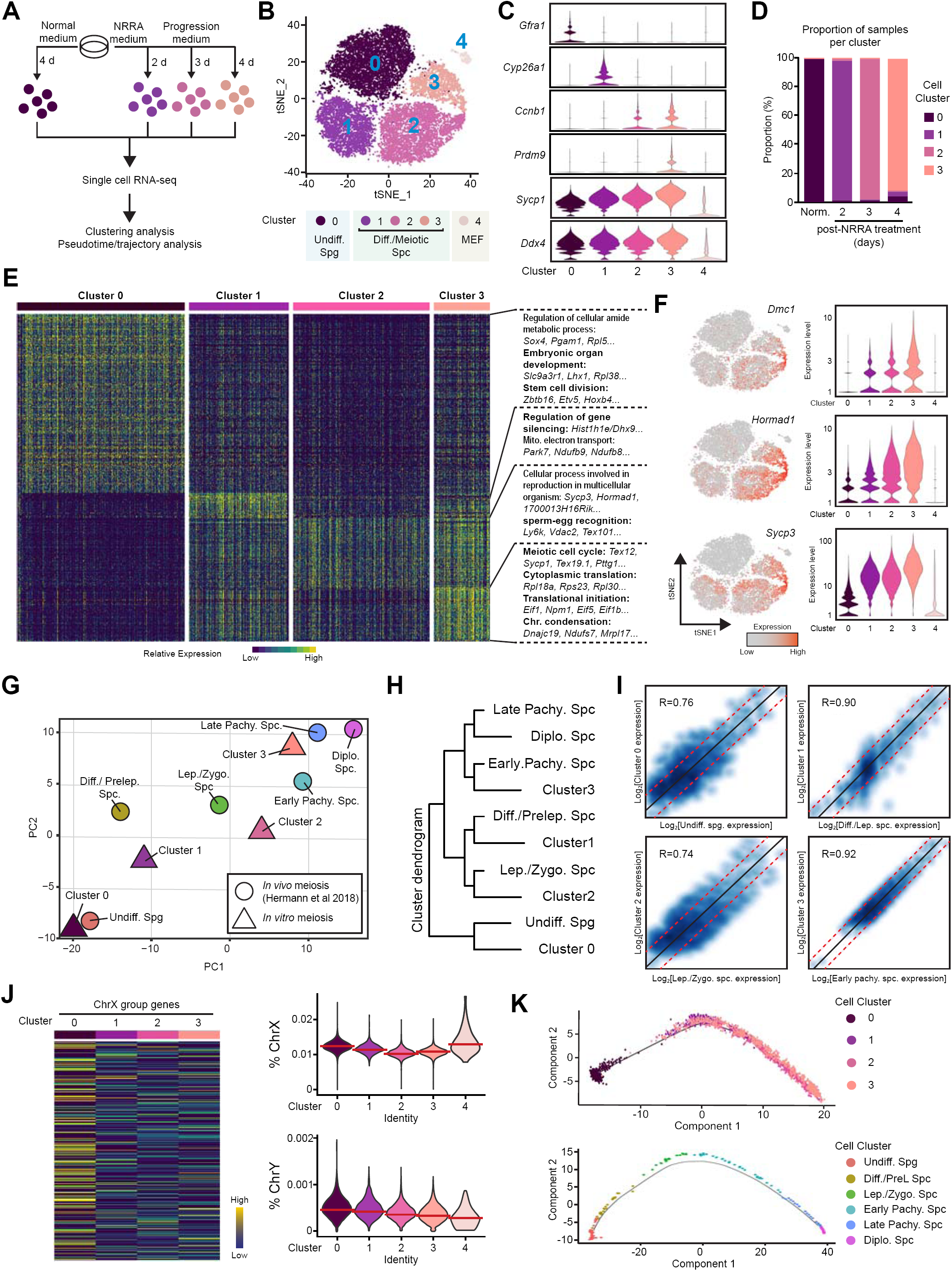
scRNA-seq analysis of primary GS cell culture during NRRA-induced meiotic initiation and progression. (**A**) Workflow of scRNA-seq experiment. Number of cells collected: 6,884 (0 day), 6,936 (2 day), 6,200 (3 day), and 5,587 (4 day). (**B**) A t-distributed Stochastic Neighbor Embedding (tSNE) plot for analyzed cells. Cluster 0 – 3 were germ cells and Cluster 4 was somatic cells. Number of cells selected for analysis: Cluster 0 (6,487 cells), Cluster 1 (5,286 cells), Cluster 2 (3,860 cells), Cluster 3 (2,170 cells), Cluster 4 (285 cells). (**C**) Violin plots showing the expression level of representative genes in each cluster. (**D**) A bar plot showing the proportion of the different cell clusters at different time points. (**E**) A heatmap showing the expression of marker genes and GO functions in each cluster. (**F**) Gene expression patterns of indicated genes on tSNE plots and violin plots. (**G**) PCA analysis showing the relationship between single cell clusters from NRRA-treated culture *in vitro* (n = 4 clusters) and early mouse spermatogenesis *in vivo* (n = 6 clusters) based on the transcriptional profiles of commonly expressed genes (n = 7,803 genes). (**H**) Hierarchical clustering showing the relationship between single cell clusters from NRRA-treated culture *in vitro* and early mouse spermatogenesis *in vivo*. (**I**) Scatter plots comparing marker genes expression profile between *in vitro* clusters and *in vivo* clusters. (**J**) (Left) Heatmap of sex chromosome genes in Clusters 0 – 3. (Right). Violin plots showing percentage of X and Y chromosome genes profiles in Clusters 0 – 3. The median is shown by a red line. (**K**) Scatter plots showing cells along the projected pseudo-time in Cluster 0 – 3 (upper) and in indicated clusters from early mouse spermatogenesis *in vivo* (lower).

Analysis of the distribution of the different cell clusters at each time point demonstrate a dramatic transition of cell populations (**Fig. 2D**). Analysis of GO functional terms for DEGs revealed rapid changes in their cellular processes (**Fig. 2E**). Cluster 0 is enriched in genes involved in regulation of cellular amide metabolic process (e.g., *Sox4*), stem cell division (e.g., *Zbtb16, Etv5*), etc (**fig. S12A, table S5**). Cluster 1 is enriched in genes involved in the regulation of gene silencing (e.g., *Hist1h1e*), mitochondrial electron transport, NADH-ubiquinone (e.g., *Park7*), retinoic acid and metabolic process/response (e.g., *Stra8*), etc (**fig. S12B**). Cluster 2 is enriched in genes involved in reproduction in multicellular organisms (e.g., *Sycp3*), sperm-egg recognition (e.g., *Ly6k*), etc (**fig. S12C**). Cluster 3 is enriched in genes for meiotic cell cycle (e.g., *Prdm9*), cytoplasmic translation (e.g., *Rpl18a*), translation initiation (e.g., *Eif1*), chromosome condensation (e.g., *Dnajc19*), etc (**fig. S12D**).

Importantly, cells from Cluster 1 to Cluster 3 exhibit a progressive and coordinated upregulation of meiotic genes bearing GO terms for mouse DNA double-strand break formation, mouse histone variants, meiotic nuclear division, mouse synapsis (**fig. S13**). Notably, meiotic genes that are fully (*Dmc1, Hormad1, Mei1, M1ap*) or partially (*Smc1b, Stag3, Sycp1, Sycp2, Sycp3, Ugt8a, Meioc*) (**Fig. 2F**; **fig. S14**) dependent on STRA8 expression displayed progressive upregulation from Cluster 1 to Cluster 3. Since activation of RA signaling with *Stra8* expression is not sufficient to induce their expression *in vitro* and *in vivo* (**fig. S8**) (*19*), this data suggests that nutrient restriction is required to work with RA to meiotic gene programs.

To assess the relationship between NRRA-induced meiotic initiation and progression *in vitro* with those during *in vivo* meiosis during spermatogenesis, we performed principal component analysis (PCA) analysis with a published scRNA-seq database (*20*) (**fig. S15**), which shows that the transcriptional profiles of Clusters 0 to 3 correlate with meiotic initiation (leptotene, the first stage of meiotic prophase) and progression (zygotene and early pachytene) during *in vivo* spermatogenesis (*20*) (**Fig. 2G**). Subsequently, hclust and differential gene correlation analysis (DGCA) demonstrate independently that cell population in Cluster 0 from *in vitro* meiosis correlates with undifferentiated spermatogonia during *in vivo* spermatogenesis, cell population in Cluster 1 with differentiating spermatogonia/preleptotene spermatocytes, cell population in Cluster 2 with leptotene/zygotene spermatocytes, and cell population in Cluster 3 with early pachytene spermatocytes (**Fig. 2H and I**). A hallmark event that occurs in male germ cells during pachytene stage is meiotic sex chromosome inactivation (MSCI) (*21*). Similarly, we observed a progressive decline in sex chromosome gene transcription from Cluster 0 to Cluster 3, supporting that cells in Cluster 3 have reached to a pachytene-like stage (**Fig. 2J**).

Moreover, past study indicates that early meiosis during *in vivo* spermatogenesis follows a step-wise pattern without lineage branching (*20*) (**Fig. 2K**). To examine the transcriptional dynamics of NRRA-induced *in vitro* meiosis, we performed pseudo-time analysis, which shows that the projected timeline recapitulated early meiosis during *in vivo* spermatogenesis (**Fig. 2K, S16-18**). The pseudo-time indicates that Cluster 0 is mainly at the start of the projected timeline trajectory, that Clusters 1, followed by Cluster 2, is positioned in the middle, and that Cluster 3 is at the end (**Fig. 2K, S16-18**). Consistently, expression of genes specific to undifferentiated spermatogonia was located preferentially at the beginning of the trajectory, while expression of genes functionally involved in many essential meiotic programs (e.g., cohesion, DNA DSB formation, chromosome segregation) was located preferentially towards to the end of the trajectory, which further supports the validity of the analysis (**fig. S17 and S18**).

*Stra8* is a best characterized gatekeeper of meiotic initiation in vertebrates (*8*). To examine whether NRRA-induced meiotic initiation meets this genetic requirement, we generated primary GS cell culture using *Stra8*-deficient mice. Consistent with the *in vivo* role of STRA8 in meiotic DSB formation (*8, 22*) (**fig. S19**), *Stra8*-deficient culture did not exhibit DMC1 foci upon NRRA treatment (**Fig. 3A**), suggesting that NRRA-induced *in vitro* meiotic DSB formation requires *Stra8* (**Fig. 3A**). Moreover, RNA-seq analysis shows that WT, *Stra8*-deficient, and *Spo11*-deficient cultures exhibited similar transcriptome profiles in normal medium (**Fig. 3B**). However, upon parallel NRRA treatment, *Stra8*-deficient culture underwent a unique transcriptomic change from WT and *Spo11*-deficient cultures, in that *Stra8*-deficient culture fails to upregulate a cluster of genes, which is enriched for meiosis-related GO terms (meiotic cell cycle, meiosis I, synapsis) (**Fig. 3B; Table S6**) (*2*). This is consistent with the essential role of STRA8 in activating the gene programs of meiotic prophase, and indicates that the lack of meiotic DSBs in *Stra8*-deficient and *Spo11*-deficient cultures resulted from discrete mechanisms (SPO11 is an enzyme that catalyze meiotic DSBs). GSVA analysis shows that indeed STRA8 sits at the foundation of NRRA-induced meiotic initiation: in contrast to WT and *Spo11*-deficient cultures, despite activation of retinoid acid receptor signaling pathway, gametogenesis- and meiosis-related pathways were not activated in *Stra8*-deficient culture upon NRRA treatment (**Fig. 3C**). Moreover, STRA8-dependent meiotic genes (*Sycp3, Mei1, Dmc1, Stag3, Sycp2*) were only upregulated in WT, but not *Stra8*-deficient, culture (**Fig. 3D; fig. S20; Table S7**). And the genes only upregulated in *Stra8*-deficient culture bears no meiosis-related GO terms, which is consistent with the robust meiotic initiation arrest phenotype of *Stra8*-deficient germ cells *in vivo*. Interestingly, our system reveals that many genes associated with undifferentiated spermatogonia were not downregulated in *Stra8*-deficient culture (e.g., *Pou5f1, Etv5*), which is in line with the role for STRA8 in promoting spermatogonial differentiation (*7*) (**Fig. 3E; fig. S21; Table S7**).

**Fig. 3.**
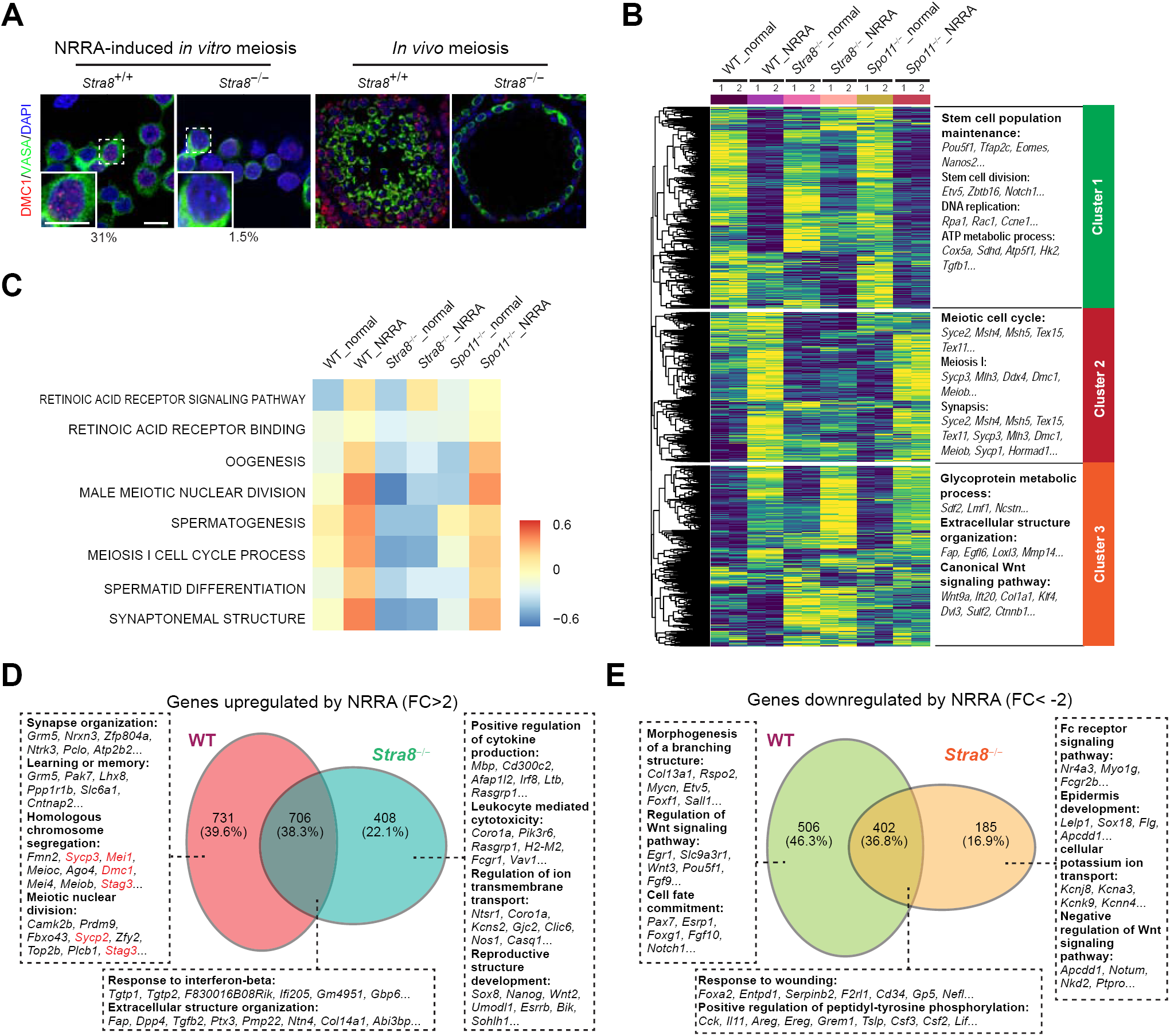
*Stra8* is required for NRRA-induced meiotic initiation. (**A**) Immunostaining for DMC1 (red), DDX4 (green), and DAPI (blue) in *Stra8*^+/+^ and *Stra8*^™/™^ primary spermatogonia culture. Scale bars, 10 µm. (**B**) (Left) UHC and heatmap of gene expression in cultured primary spermatogonia with indicated genotypes for 2 days. (Right) Top GO enrichments with representative genes in each cluster. (**C**) GSVA analysis for indicated genotypes. In the heatmap, rows are defined by the selected gene sets, and columns by consensus scores for each genotype. Group enriched gene sets are highlighted by different color. (**D and E**) Venn plots with top GO enrichments and representative genes for NRRA-upregulated (**D**) and -downregulated (**E**) genes in *Stra8*^+/+^ and *Stra8*^™/™^ cultures.

Our results show for the first time that *Stra8*-dependent and *Spo11*-dependent meiotic DSBs are induced *in vitro* (**Figs. 1D and 3A**). To further examine the processing of meiotic DSB formed *in vitro*, our spread analysis shows that, similar to *in vivo* meiosis, sing-strand DNA (ssDNA)-binding proteins, SPATA22 and MEIOB (*23, 24*), were progressively recruited onto the meiotic chromosomes from a leptotene-like to an early pachytene-like stage, suggesting that meiotic DSBs formed *in vitro* were resected into ssDNA before recruiting meiotic recombinases RAD51 and DMC1 for repair (**fig. S22A**). Quantification show that similar numbers of SPATA22, MEIOB, RAD51, and DMC1 foci were detected on meiotic chromosomes per germ cell between *in vivo* meiosis and *in vitro* meiosis (**fig. S22B and C**). In addition, histone methyltransferase PRDM9 directs meiotic DSBs to be distributed to recombination hotspots by (*25, 26*). PRDM9 expression is induced by NRRA in cultures at both mRNA and protein levels (**Fig. 2**; **fig. S23A**). Consistently, chromatin immunoprecipitation (ChIP) assay for DMC1-associated ssDNA fragments, a direct method to detect recombination hotspots (*27*), revealed that the DSBs formed *in vitro* upon NRRA treatment were mapped to the strong hotspots of meiotic recombination *in vivo* (*28*) (**fig. S23B**).

To examine the mechanism of nutrient restriction-induced meiotic initiation, we identified that nutrient restriction upregulated 120 transcription factor (TF) genes, whose expression does not require RA (**Fig. 4A and B**). To identify those potentially involved in regulating the meiotic gene programs, we examined the expression of these genes in the transcriptomic database for mouse spermatogenesis and found that 30 of them are expressed before meiotic prophase (**Fig. 4B and C**) (*20*). Then, we further investigated their correlation with meiotic gene expression in the Genotype-Tissue Expression database (GTEx) (*29*). Using *Dmc1, Sycp3, Hormad1* as three representatives, we found that 11 of them shows strong (Pearson correlation > 0.6) or moderate correlation (Pearson correlation > 0.4) with meiotic gene expression (**Fig. 4B and D, fig. S24**). We further confirmed that the expression of these 11 TFs *in vivo* does not require RA (**fig. S25**). Notably, these 11 TFs include Sohlh1 and Sox3, two characterized TFs implicated in early meiosis and spermatogenesis. Sohlh1 is basic helix-loop-helix (bHLH) transcription factor, whose deletion results in many tubules lacking meiotic spermatocytes (*30*). Recently, Sohlh1 is shown to meiotic gene expression (*Sycp1, Sycp3*) by directly binding to their proximal promoters (*31*). Sox3 is expressed exclusively in spermatogonia committed to differentiation (*32*), and loss of Sox3 impairs early meiosis (*33*). Thus, these data suggest that nutrient restriction induces a distinct network of RA signaling-independent TFs to activate meiotic gene program.

**Fig. 4.**
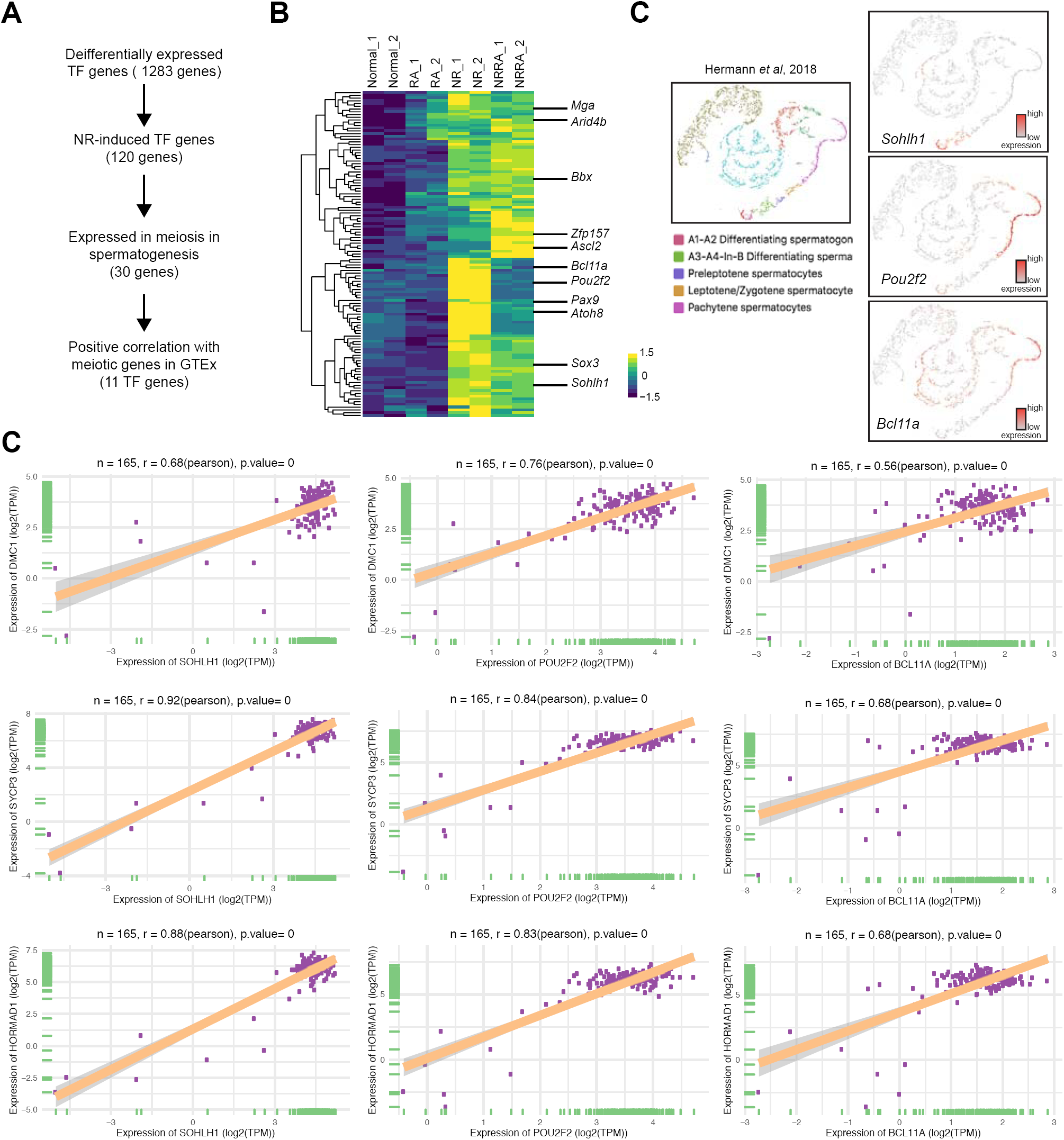
Nutrient restriction induces a network of TF genes involved in early meiosis. (**A**) A workflow showing filters to identify genes of interest. (**B**) Heatmap of 120 TF genes upregulated by nutrient restriction. 11 identified TF genes involved in early meiosis are shown on the right. (**C**) Gene expression patterns of 3 representative TF genes during early spermatogenesis from a published scRNA-seq database (*20*). (**D**) GTEx database showing correlation of representative TF genes (*Sohlh1, Pou2f2, Bcl11a*) with selected meiotic genes in 165 testis samples.

Nutrient restriction is a metabolic cue. Our scRNA-seq revealed that, while genes involved in the glycolytic pathways were downregulated, genes involved in mitochondrial function pathways were upregulated during NRRA-induced *in vitro* meiotic initiation and progression (Cluster 0 to Cluster 3), which mirrors the transition from glycolysis to mitochondrial biogenesis and oxidative phosphorylation during *in vivo* spermatogonial differentiation (*34*) (**fig. S26**). Together, our data at the transcriptomic, cytologic, mechanistic, and metabolic levels suggest that nutrient restriction in combination with RA induces meiotic initiation that faithfully recapitulates that during *in vivo* spermatogenesis. Interestingly, unlike the nutrient-rich basal compartment where mitotic spermatogonia reside, the apical compartment of the mammalian seminiferous tubule, where meiosis takes place, is a nutrient-restricted microenvironment (i.e., low in glucose and most amino acids) created by the blood-testis barrier (BTB) (*35*). Thus, nutrient restriction as a physiological stimulus for meiotic initiation that requires BTB function warrants further characterization. Our study provides an *in vitro* platform to study meiotic initiation and to facilitate production of haploid gametes in culture.

## ACKNOWLEDGEMENTS

We thank L. L. Heckert for helpful input on this study, P. E. Cohen for chromosome spread technique, K. E. Orwig for suggestion on primary mouse GS cell culture, and J. P. Wang for MEIOB antibody and insightful discussion.

## Author contributions

X.Z. performed all the experiments. S.G. assisted the analyses of RNA-seq and scRNA-seq data. X.Z. and N.W. designed the experiments, analyzed and curated the data. N.W. and X.Z. wrote the manuscript.

## Data and materials availability

All data are available in the manuscript or the supplementary material.

## Supplementary Materials

This manuscript contains 26 Supplementary Figures and 7 Supplementary Tables.

