## Supplementary Materials for "Nutrient restriction, inducer of yeast meiosis, induces meiotic initiation in mammals"

**FIGURE LEGENDS**

**fig. S1. Primary GS cell culture.**

(**A**) Confocal immunofluorescence detection of spermatogonia marker by staining of the cultured spermatogonia *in vitro*. (**B**) Morphology of cultured GS cells when exposed to NRRA. Appearance of $A_{pr}$, $A_4$, $A_8$ and $A_{al}$ spermatogonia, reminiscent of in vivo spermatogonial differentiation, were detected after 2 days. Scale bar, 20 μm. (**C**) Cell lysates from GS cell culture with indicated treatments for 2 days were subjected to Western-blot analysis using γH2AX antibody. (**D**) Immunostaining of cultured GS cells with indicated antibodies as markers for undifferentiated spermatogonia combined with DAPI (blue). Scale bar, 10 μm.

**fig. S2. NRRA induces meiotic gene expression in F9 premeiotic cells.**

qRT-PCR analysis of meiotic genes in F9 cells with indicated treatment normalized to *β-actin*. Data represent mean ± SD; n = 3. $P < 0.05$ (Student's *t*-test). F9 cells were cultured in DMEM medium with 10% FBS (complete medium). Nutrient restriction was applied to F9 cells by adding 90% of EBSS to the complete medium.

**fig. S3. Differentially expressed genes (DEGs) in primary GS cell culture upon normal medium, RA, NR, and NRRA treatment.**

Scatter plot representations of DEGs between the indicated groups (left panel). The up- or down-regulated genes (fold change > 2 or < -2) in each culture conditions are plotted in red and blue, respectively. The GO function analyses of the upregulated (middle panel) and downregulated (right panel) DEGs between two treatment groups are presented. Representative genes in each GO category are indicated.

**fig. S4. GO function analysis for differentially expressed genes upon normal medium, RA, NR, NRRA treatment.**

(**A-D**) The bubble plots show GO enrichment analysis for four gene clusters in Fig. 1E.

**fig. S5. GSVA analysis for RA and NRRA treatments in primary GS cell culture.**

(A) GSEA analysis reveals distinct enriched gene sets between different groups. In the heatmap, rows are defined by the selected gene sets, and columns by consensus scores for each group. Groups enriched gene sets are highlighted by different color.

**fig. S6. RNA-seq analysis of primary GS cell culture in CD1 background.**

(**A**) UHC and heatmap of globally changed genes in primary GS cell cultures with indicated treatments and genetic backgrounds. (**B**) Scatter plot representations of DEGs between normal medium and NRRA treatment groups in CD1 background. The up or down-regulated genes (fold change > 2 or < -2) in each cell culture conditions are plotted in red and blue, respectively. (**C**) The bar plots show GO enrichment analysis data for up and down genes (fold change >2 or < -2) when NRRA treatment in CD1 background.

**fig. S7. Changes in the expression of early meiosis-related genes upon NRRA treatment in primary GS cell culture.**

(**A**) Heatmap showing the expression levels of 104 mouse early meiosis-related genes (Soh *et al*, 2015) with indicated treatments. (**B**) Heatmap showing the expression levels of 107 human early meiosis-related genes (Guo *et al.*, 2017) with indicated treatments.

**fig. S8. Changes of meiosis-related gene expression upon RA treatment in a previously reported scRNA-seq database.**

(**A**) Heatmap showing the expression levels of 165 meiosis-related genes in DMSO and RA treatment groups in testes of postnatal day 5. (**B**) Violin plots showing representative meiotic genes profiles with DMSO and RA treatments. Please note that *Stra8* is a direct target gene of RA, and Stra8 expression serves as “a positive control” for the activation of RA signaling upon RA treatment.

**fig. S9. 10X genomics quality control in scRNA-seq.**

(**A**) QC metrics. (**B**) A subset of features that exhibit high cell-to-cell variation in the dataset. (**C**) An alternative heuristic method generates an ‘Elbow plot’: a ranking of principle components based on the percentage of variance explained by each one (ElbowPlot function).

**fig. S10. Expression pattern of marker genes for Cluster 0 – 4 in scRNA-seq.**

Violin plot and scatter plot showing the expression of marker genes for undifferentiated spermatogonia (**A**), differentiating spermatogonia (**B**), meiocytes (**C**), and fibroblast (**D**) in each cluster.

**fig. S11. Expression of undifferentiated spermatogonia and meiosis genes in primary GS cell culture.**

(**A**) qRT-PCR analysis of undifferentiated spermatogonia marker genes in primary GS cell culture with indicated treatments normalized to *β-actin*. Data represent mean ± SD; n = 3.

(**B**) qRT-PCR analysis of meiotic genes in primary GS cell culture with indicated treatments normalized to β-actin. Data represent mean ± SD; n = 3.

**fig. S12. GO function and KEGG analysis for Cluster 0 to Cluster 3 in scRNA-seq.**

(**A**) The bubble plots show GO enrichment and KEGG pathway analysis for cluster 0 DEGs.

(**B**) The bubble plots show GO enrichment and KEGG pathway analysis for cluster 1 DEGs.

(**C**) The bubble plots show GO enrichment and KEGG pathway analysis for cluster 2 DEGs.

(**D**) The bubble plots show GO enrichment and KEGG pathway analysis for cluster 3 DEGs.

**fig. S13. Analysis of Clusters 0 – 3 and meiotic pathways enriched in GO function terms.**

(**A-B**) Heatmap of "DNA double strand break formation" and "meiotic prophase-specific histone variants" pathway-enriched genes.

(**C-D**) Heatmap (left) and violin plot (right) of the enriched "meiotic nuclear division" and "synapsis" pathway-enriched genes. Violin plots showing the relative percentage expression levels of "meiotic nuclear division" and "synapsis" pathway-enriched genes versus all the genes in each cluster.

**fig. S14. Expression of meiotic genes in scRNA-seq.**

(**A**) Violin plot and scatter plot showing the expression of meiotic genes whose expression is fully dependent on STRA8 in each cluster.

(**B**) Violin plot and scatter plot showing the expression of meiotic genes whose expression is partially dependent on STRA8 in each cluster.

(**C**) Heatmap describing the expression levels of 165 early meiosis-related genes in Cluster 0 – 4 from single cell RNA-seq. Please compare to fig. S8, which shows that RA alone is not sufficient to induce meiosis-related gene expression (except *Stra8* and *Rec8*, which are direct targets of RA).

**fig. S15. Analysis of early spermatogenesis to meiotic prophase in a published scRNA-seq database on mouse spermatogenesis.**

(**A-C**) Identification of different stages of early spermatogenesis to meiotic prophase.

(**D**) Pseudo-time analysis of different stages early spermatogenesis to meiotic prophase.

**fig. S16. Single-cell trajectories reveal dynamic biological transitions in Cluster 0 – 3 in pseudo-time.**

(**A**) Clusters of genes that were differentially expressed across pseudotime and top GOs.

(**B**) KEGG analysis for Clusters 0 – 3 in pseudotime DEGs.

**fig. S17 – S18. Single-cell trajectories reveal dynamic meiotic function changes in Cluster 0 – 3 (C1-C4) in pseudotime.**

Expression levels (vertical axis) of key genes for “meiotic cell cycle”, “cohesion”, “chromosome segregation” and “DNA double-strand break formation” are ordered in pseudotime.

**fig. S18. Single-cell trajectories reveal dynamic meiotic function changes in Cluster 0 – 3 (C1-C4) in pseudotime.**

Expression levels (vertical axis) of key genes for “DNA repair”, “homologous recombination” and “synapsis” are ordered in pseudotime.

**fig. S19. *Stra8*-deficient germ cells (C57BL/6XDBA/2 F1 hybrid background) lack meiotic DSB formation.**

Immunofluorescent staining for DMC1, MEIOB, and SPATA22 is showed in testicular cross-sections from WT, *Stra8*$^{-/-}$, *Spo11*$^{-/-}$, *Stra8*$^{-/-}$;*Spo11*$^{-/-}$ mice (28 days of age). Scale bars, 10 µm.

**fig. S20. GO function analysis for upregulated genes in WT and *Stra8*$^{-/-}$ primary GS cell culture.**

(**A**) The bubble plots show GO enrichment analysis for genes upregulated in WT only.

(**B**) The bubble plots show GO enrichment analysis for genes upregulated in both WT and *Stra8*KO.

(**C**) The bubble plots show GO enrichment analysis for genes upregulated in *Stra8*KO only.

**fig. S21. GO function analysis for downregulated genes in WT and *Stra8*$^{-/-}$ primary GS cell culture.**

(**A**) The bubble plots show GO enrichment analysis for genes downregulated in WT only.

(**B**) The bubble plots show GO enrichment analysis for genes downregulated in both WT and *Stra8*KO.

(**C**) The bubble plots show GO enrichment analysis for genes downregulated in *Stra8*KO only.

**fig. S22. Cytological analysis of meiotic DSB formation *in vitro*.**

(**A**) RAD51, DMC1, MEIOB, and SPATA22 foci on meiotic chromosomes (SYCP3) were used to identify different stages of meiosis (leptotene-like to zygo-/early-pachytene-like stage). Percentages of leptotene-like to zygo-/early-pachytene-like stage in culture on each day following treatment is shown on the right.

(**B**) Representative chromosome spreads stained by RAD51, DMC1, MEIOB, SPATA22, and SYCP3 from juvenile mice (day 14 of age) or primary GS cell culture treated with NRRA to induce meiosis are shown.

(**C**) Quantification of foci for zygonema-early pachynema stage. Error bars, mean ± SD. Each solid circle indicates the total number of foci from a single nucleus. Total number of cells quantified from three independent cultures are shown on each graph. n.s., not significant.

**fig. S23. Distribution of meiotic DSB formation *in vitro* to genomic hotspots of recombination.**

**(A)** Immunofluorescence staining for SYCP3 and PRDM9 on chromosome spreads in testicular germ cells from juvenile mice (day 14 of age) or in cells from primary GS cell culture following meiotic initiation induced by NRRA and progression. Scale bars, 10 μm.

**(B)** Enrichment of the DNA corresponding to several published hotspots by anti-DMC1 ChIP estimated by qPCR. ChIP was performed from juvenile testis extracts (*in vivo*; upper panel) and primary GS cell cultured following NRRA treatment (*in vitro*; lower panel). Enrichment of the DNA corresponding to genomic regions on Chr17, Chr5, Chr8, Chr11, Chr13, Chr14, and the *β-actin* gene was estimated by qPCR. All data were averages of three independent experiments.

**fig. S24. Correlation of NR-induced TF genes with representative meiotic genes.**

Scatter plots showing correlation of representative TF genes with selected meiotic genes in 165 testis tissues based on the data from Genotype Tissue Expression (GTEx). Note that every dot represents one tissue type. Correlation coefficient (r) and *P* values were calculated by Pearson's correlation analysis. The expression pattern of each TF genes in scRNA-seq dataset during *in vivo* spermatogenesis is shown on the right.

**fig. S25. NR-induced TF genes do not respond to RA treatment *in vivo.***

(**A**) Heatmap describing the expression levels of RA-responsive genes and NR-induced TF genes in DMSO and RA treatment in testes at day 5 of age (Velte *et al*, 2019).

(**B**) Violin plots showing NR-induced TF genes profiles in DMSO and RA treatment in *in vivo* (Velte *et al*, 2019).

**fig. S26. Analysis of metabolic pathways enriched in GO function terms in Clusters 0 – 3.**

(**A-F**) Heatmap (left) and violin plot (right) of the enriched glycolytic process, glycolysis and oxidative phosphorylation related genes. Violin plot showing the relative percentage expression levels of glycolytic process, glucose metabolic process, and oxidative phosphorylation related genes versus all the genes in each cluster.

(**G-J**) Heatmap (upper) and violin plot (lower) of the enriched mitochondrial respiratory chain complex I & IV assembly and mitochondrial organization related genes. Violin plot showing the relative percentage expression levels of mitochondrial respiratory chain complex I & IV assembly and mitochondrial electron transport NADH to ubiquinone related genes versus all the genes in each cluster.

### fig. S1

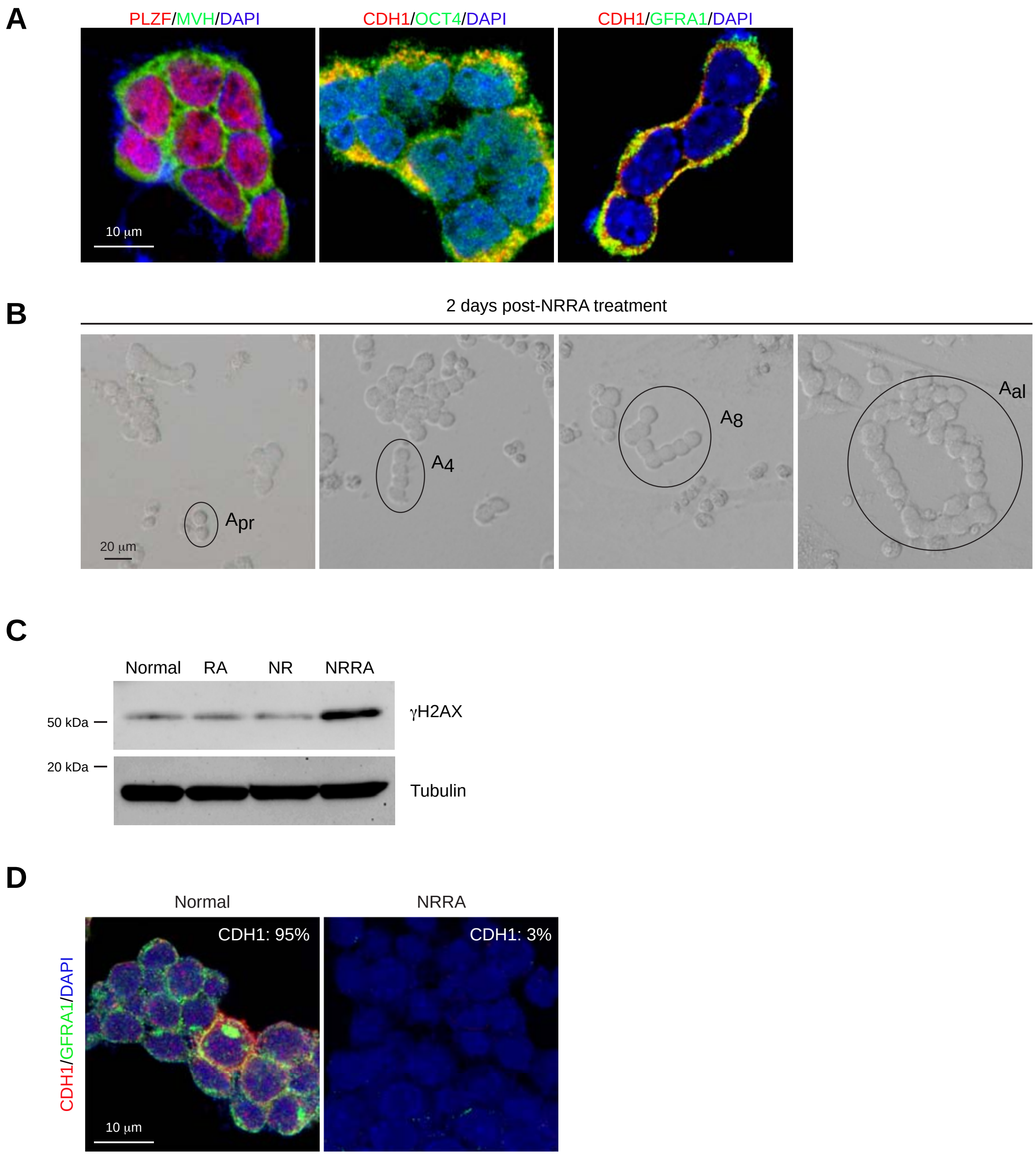
A
PLZF/MVH/DAPI
CDH1/OCT4/DAPI
CDH1/GFRA1/DAPI
10 μm
B
2 days post-NRRA treatment
Apr
A4
A8
Aal
20 μm
C
Normal
RA
NR
NRRA
50 kDa
20 kDa
γH2AX
Tubulin
D
Normal
NRRA
CDH1/GFRA1/DAPI
CDH1: 95%
CDH1: 3%
10 μm

### fig. S2

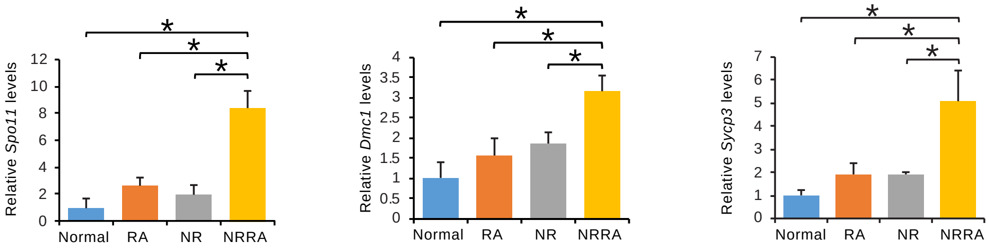

Relative *Spo11* levels
12
10
8
6
4
2
0
Normal
RA
NR
NRRA
Relative *Dmc1* levels
4
3.5
3
2.5
2
1.5
1
0.5
0
Relative *Sycp3* levels
7
6
5
4
3
2
1
0

### fig. S3

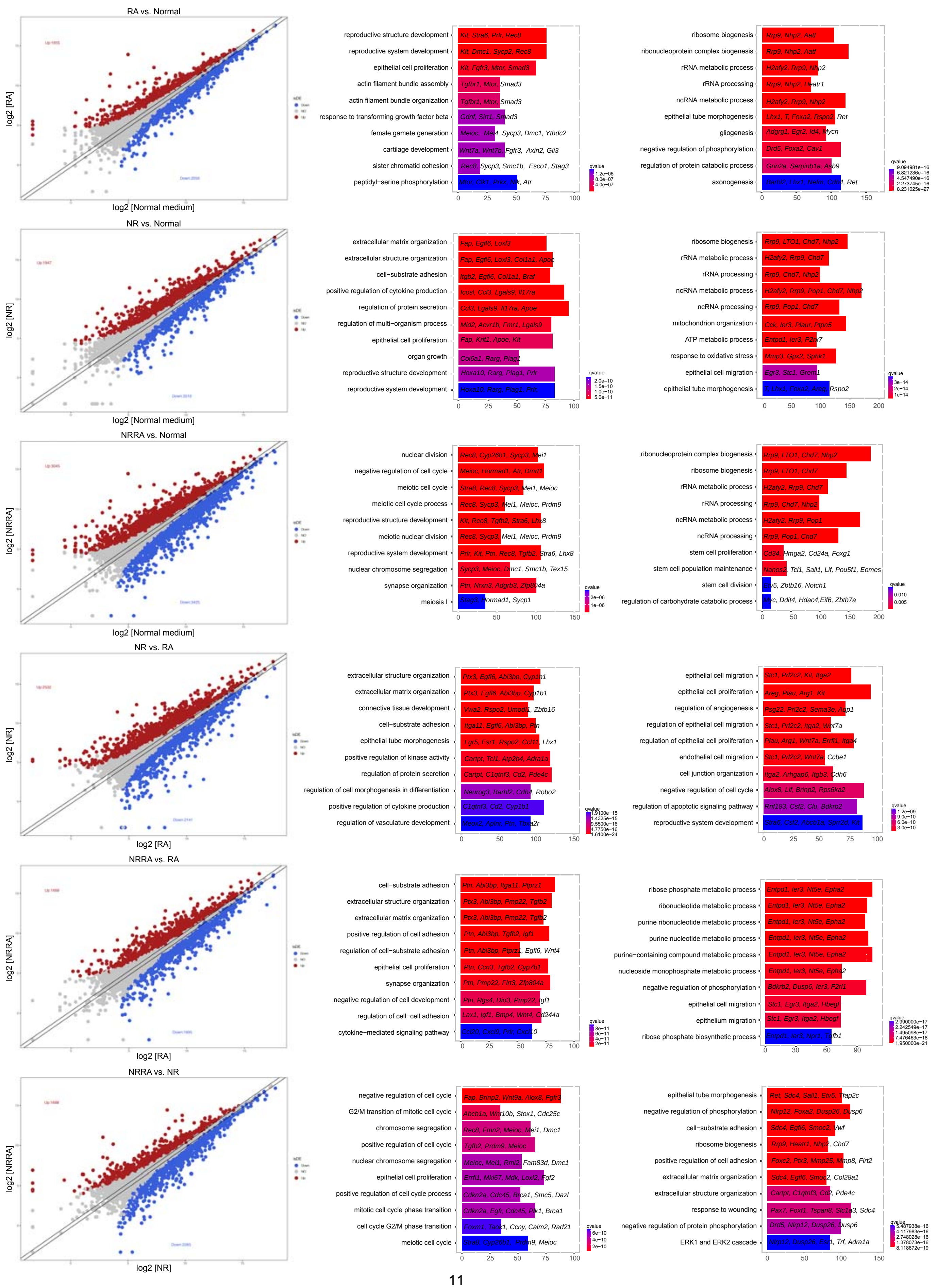
RA vs. Normal
log2 [RA]
log2 [Normal medium]
reproductive structure development
reproductive system development
epithelial cell proliferation
actin filament bundle assembly
actin filament bundle organization
response to transforming growth factor beta
female gamete generation
cartilage development
sister chromatid cohesion
peptidyl−serine phosphorylation
Kit, Stra8, Prlr, Rec8
Kit, Dmc1, Sycp2, Rec8
Kit, Fgfr3, Mtor, Smad3
Tgfbr1, Mtor, Smad3
Tgfbr1, Mtor, Smad3
Gdnf, Sirt1, Smad3
Meioc, Mei4, Sycp3, Dmc1, Ythdc2
Wnt7a, Wnt7b, Fgfr3, Axin2, Gli3
Rec8, Sycp3, Smc1b, Esco1, Stag3
Nbn, Clk2, Prkx, Nlk, Atr
ribosome biogenesis
ribonucleoprotein complex biogenesis
rRNA metabolic process
rRNA processing
ncRNA metabolic process
epithelial tube morphogenesis
gliogenesis
negative regulation of phosphorylation
regulation of protein catabolic process
axonogenesis
Rrp9, Nhp2, Aatf
Rrp9, Nhp2, Aatf
H2afy2, Rrp9, Nhp2
Rrp9, Nhp2, Heatr1
H2afy2, Rrp9, Nhp2
Lhx1, T, Foxa2, Rspo2, Ret
Adgrg1, Egr2, Id4, Mycn
Drd5, Foxa2, Cav1
Grin2a, Serpinb1a, Hdac9
Barhl2, Lhx1, Nefm, Cdh4, Ret
NR vs. Normal
log2 [NR]
log2 [Normal medium]
extracellular matrix organization
extracellular structure organization
cell−substrate adhesion
positive regulation of cytokine production
regulation of protein secretion
regulation of multi−organism process
epithelial cell proliferation
organ growth
reproductive structure development
reproductive system development
Fap, Egfl6, Loxl3
Fap, Egfl6, Loxl3, Col1a1, Apoe
Itgb2, Egfl6, Col1a1, Braf
Icosl, Ccl3, Lgals9, Il17ra
Ccl3, Lgals9, Il17ra, Apoe
Mid2, Acvr1b, Fmr1, Lgals9
Fap, Krit1, Apoe, Kit
Col6a1, Rarg, Plag1
Hoxa10, Rarg, Plag1, Prlr
Hoxa10, Rarg, Plag1, Prlr
ribosome biogenesis
rRNA metabolic process
rRNA processing
ncRNA metabolic process
ncRNA processing
mitochondrion organization
ATP metabolic process
response to oxidative stress
epithelial cell migration
epithelial tube morphogenesis
Rrp9, LTO1, Chd7, Nhp2
H2afy2, Rrp9, Chd7
Rrp9, Chd7, Nhp2
H2afy2, Rrp9, Pop1, Chd7, Nhp2
Rrp9, Pop1, Chd7
Cck, Ier3, Plaur, Ptpn5
Entpd1, Ier3, P2rx7
Mmp3, Gpx2, Sphk1
Egr3, Stc1, Grem1
T, Lhx1, Foxa2, Areg, Rspo2
NRRA vs. Normal
log2 [NRRA]
log2 [Normal medium]
nuclear division
negative regulation of cell cycle
meiotic cell cycle
meiotic cell cycle process
reproductive structure development
meiotic nuclear division
reproductive system development
nuclear chromosome segregation
synapse organization
meiosis I
Rec8, Cyp26b1, Sycp3, Mei1
Meioc, Hormad1, Atr, Dmrt1
Stra8, Rec8, Sycp3, Mei1, Meioc
Rec8, Sycp3, Mei1, Meioc, Prdm9
Kit, Rec8, Tgfb2, Stra6, Lhx8
Rec8, Sycp3, Mei1, Meioc, Prdm9
Prlr, Kit, Ptn, Rec8, Tgfb2, Stra6, Lhx8
Sycp3, Meioc, Dmc1, Smc1b, Tex15
Ptn, Nrxn3, Adgrb3, Zfp804a
Stag3, Hormad1, Sycp1
ribonucleoprotein complex biogenesis
ribosome biogenesis
rRNA metabolic process
rRNA processing
ncRNA metabolic process
ncRNA processing
stem cell proliferation
stem cell population maintenance
stem cell division
regulation of carbohydrate catabolic process
Rrp9, LTO1, Chd7, Nhp2
Rrp9, LTO1, Chd7
H2afy2, Rrp9, Chd7
Rrp9, Chd7, Nhp2
H2afy2, Rrp9, Pop1
Rrp9, Pop1, Chd7
Cd34, Hmga2, Cd24a, Foxg1
Nanos2, Tcl1, Sall1, Lif, Pou5f1, Eomes
Etv5, Zbtb16, Notch1
Myc, Ddit4, Hdac4,Eif6, Zbtb7a
NR vs. RA
log2 [NR]
log2 [RA]
extracellular structure organization
extracellular matrix organization
connective tissue development
cell−substrate adhesion
epithelial tube morphogenesis
positive regulation of kinase activity
regulation of protein secretion
regulation of cell morphogenesis in differentiation
positive regulation of cytokine production
regulation of vasculature development
Ptx3, Egfl6, Abi3bp, Cyp1b1
Ptx3, Egfl6, Abi3bp, Cyp1b1
Vwa2, Rspo2, Umodl1, Zbtb16
Itga11, Egfl6, Abi3bp, Ptn
Lgr5, Esr1, Rspo2, Ccl11, Lhx1
Cartpt, Tcl1, Atp2b4, Adra1a
Cartpt, C1qtnf3, Cd2, Pde4c
Neurog3, Barhl2, Cdh4, Robo2
C1qtnf3, Cd2, Cyp1b1
Mecx2, Apln, Ptn, Tbxa2r
epithelial cell migration
epithelial cell proliferation
regulation of angiogenesis
regulation of epithelial cell migration
regulation of epithelial cell proliferation
endothelial cell migration
cell junction organization
negative regulation of cell cycle
regulation of apoptotic signaling pathway
reproductive system development
Stc1, Prl2c2, Kit, Itga2
Areg, Plau, Arg1, Kit
Psg22, Prl2c2, Sema3e, Aqp1
Stc1, Prl2c2, Itga2, Wnt7a
Plau, Arg1, Wnt7a, Errfi1, Itga4
Stc1, Prl2c2, Wnt7a, Ccbe1
Itga2, Arhgap6, Itgb3, Cdh6
Alox8, Lif, Brinp2, Rps6ka2
Rnf183, Csf2, Clu, Bdkrb2
NRRA vs. RA
log2 [NRRA]
log2 [RA]
cell−substrate adhesion
extracellular structure organization
extracellular matrix organization
positive regulation of cell adhesion
regulation of cell−substrate adhesion
epithelial cell proliferation
synapse organization
negative regulation of cell development
regulation of cell−cell adhesion
cytokine−mediated signaling pathway
Ptn, Abi3bp, Itga11, Ptprz1
Ptx3, Abi3bp, Pmp22, Tgfb2
Ptx3, Abi3bp, Pmp22, Tgfb2
Ptn, Abi3bp, Tgfb2, Igf1
Ptn, Abi3bp, Ptprz1, Egfl6, Wnt4
Ptn, Ccn3, Tgfb2, Cyp7b1
Ptn, Pmp22, Flrt3, Zfp804a
Ptn, Rgs4, Dio3, Pmp22, Igf1
Lax1, Igf1, Bmp4, Wnt4, Cd244a
Ccl20, Cxcl9, Prlr, Cxcl10
ribose phosphate metabolic process
ribonucleotide metabolic process
purine ribonucleotide metabolic process
purine nucleotide metabolic process
purine−containing compound metabolic process
nucleoside monophosphate metabolic process
negative regulation of phosphorylation
epithelial cell migration
epithelium migration
ribose phosphate biosynthetic process
Entpd1, Ier3, Nt5e, Epha2
Entpd1, Ier3, Nt5e, Epha2
Entpd1, Ier3, Nt5e, Epha2
Entpd1, Ier3, Nt5e, Epha2
Entpd1, Ier3, Nt5e, Epha2
Entpd1, Ier3, Nt5e, Epha2
Bdkrb2, Dusp6, Ier3, F2rl1
Stc1, Egr3, Itga2, Hbegf
Stc1, Egr3, Itga2, Hbegf
NRRA vs. NR
log2 [NRRA]
log2 [NR]
negative regulation of cell cycle
G2/M transition of mitotic cell cycle
chromosome segregation
positive regulation of cell cycle
nuclear chromosome segregation
epithelial cell proliferation
positive regulation of cell cycle process
mitotic cell cycle phase transition
cell cycle G2/M phase transition
meiotic cell cycle
Fap, Brinp2, Wnt9a, Alox8, Fgfr3
Abcb1a, Wnt10b, Stox1, Cdc25c
Rec8, Fmn2, Meioc, Mei1, Dmc1
Tgfb2, Prdm9, Meioc
Meioc, Mei1, Rmi2, Fam83d, Dmc1
Errfi1, Mki67, Mdk, Loxl2, Fgf2
Cdkn2a, Cdc45, Brca1, Smc5, Dazl
Cdkn2a, Egfr, Cdc45, Plk1, Brca1
Foxm1, Taok1, Ccny, Calm2, Rad21
Prdm9, Meioc
epithelial tube morphogenesis
negative regulation of phosphorylation
cell−substrate adhesion
ribosome biogenesis
positive regulation of cell adhesion
extracellular matrix organization
extracellular structure organization
response to wounding
negative regulation of protein phosphorylation
ERK1 and ERK2 cascade
Ret, Sdc4, Sall1, Etv5, Tfap2c
Nlrp12, Foxa2, Dusp26, Dusp6
Sdc4, Egfl6, Smoc2, Vwf
Rrp9, Heatr1, Nhp2, Chd7
Foxc2, Ptx3, Mmp25, Mmp8, Flrt2
Sdc4, Egfl6, Smoc2, Col28a1
Cartpt, C1qtnf3, Cd2, Pde4c
Pax7, Foxf1, Tspan8, Slc1a3, Sdc4
Drd5, Nlrp12, Dusp26, Dusp6
Trf, Adra1a
qvalue

### fig. S4

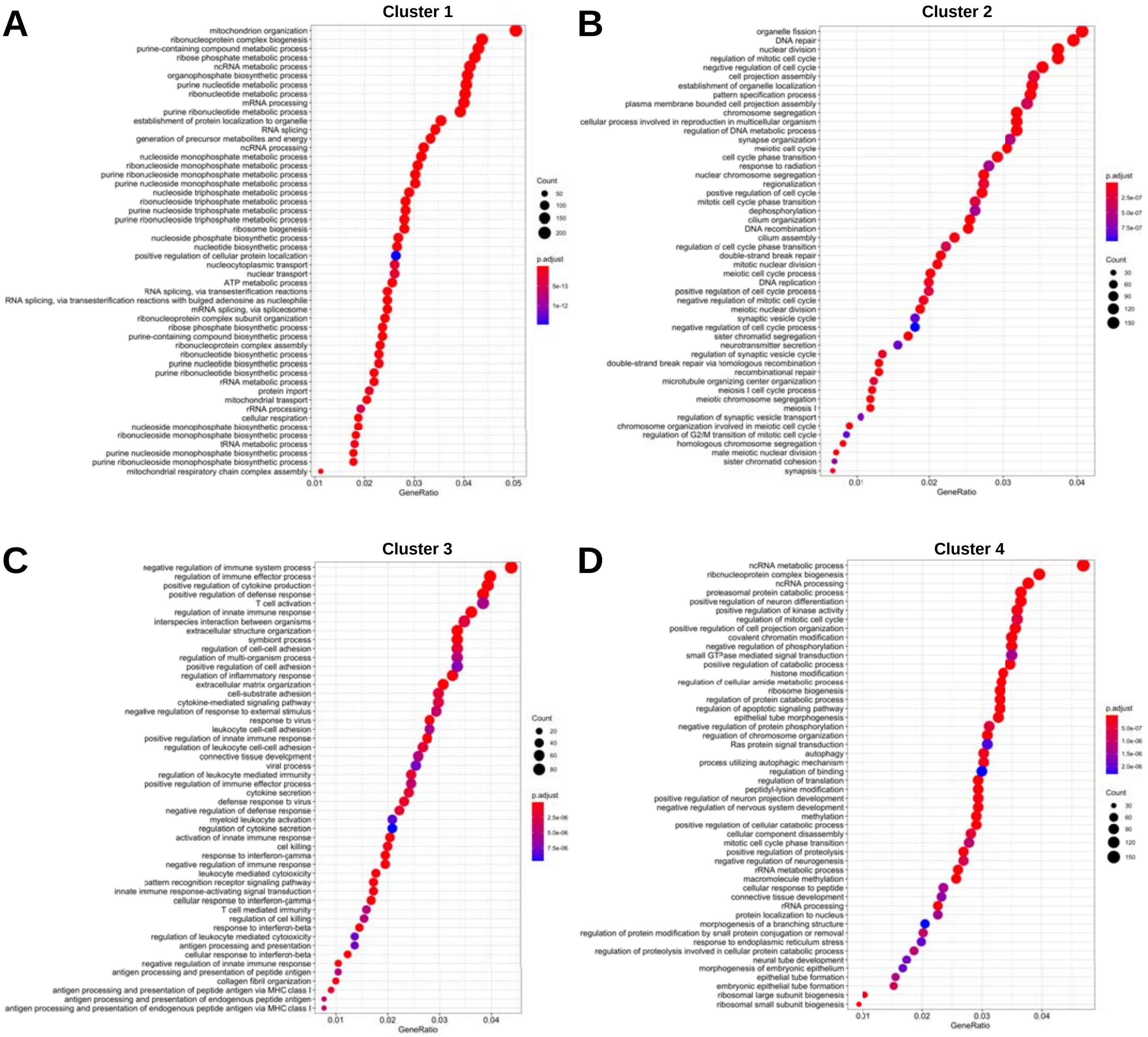
A
Cluster 1
mitochondrion organization
ribonucleoprotein complex biogenesis
purine-containing compound metabolic process
ribose phosphate metabolic process
ncRNA metabolic process
organophosphate biosynthetic process
purine nucleotide metabolic process
ribonucleotide metabolic process
mRNA processing
purine ribonucleotide metabolic process
establishment of protein localization to organelle
RNA splicing
generation of precursor metabolites and energy
ncRNA processing
nucleoside monophosphate metabolic process
ribonucleoside monophosphate metabolic process
purine ribonucleoside monophosphate metabolic process
purine nucleoside monophosphate metabolic process
nucleoside triphosphate metabolic process
ribonucleoside triphosphate metabolic process
purine nucleoside triphosphate metabolic process
purine ribonucleoside triphosphate metabolic process
ribosome biogenesis
nucleoside phosphate biosynthetic process
nucleotide biosynthetic process
positive regulation of cellular protein localization
nucleocytoplasmic transport
nuclear transport
ATP metabolic process
RNA splicing, via transesterification reactions
RNA splicing, via transesterification reactions with bulged adenosine as nucleophile
mRNA splicing, via spliceosome
ribonucleoprotein complex subunit organization
ribose phosphate biosynthetic process
purine-containing compound biosynthetic process
ribonucleoprotein complex assembly
ribonucleotide biosynthetic process
purine nucleotide biosynthetic process
purine ribonucleotide biosynthetic process
rRNA metabolic process
protein import
mitochondrial transport
rRNA processing
cellular respiration
nucleoside monophosphate biosynthetic process
ribonucleoside monophosphate biosynthetic process
tRNA metabolic process
purine nucleoside monophosphate biosynthetic process
purine ribonucleoside monophosphate biosynthetic process
mitochondrial respiratory chain complex assembly
0.01 0.02 0.03 0.04 0.05
GeneRatio
Count
50
100
150
200
p.adjust
5e-13
1e-12
B
Cluster 2
organelle fission
DNA repair
nuclear division
regulation of mitotic cell cycle
negative regulation of cell cycle
cell projection assembly
establishment of organelle localization
pattern specification process
plasma membrane bounded cell projection assembly
chromosome segregation
cellular process involved in reproduction in multicellular organism
regulation of DNA metabolic process
synapse organization
meiotic cell cycle
cell cycle phase transition
response to radiation
nuclear chromosome segregation
regionalization
positive regulation of cell cycle
mitotic cell cycle phase transition
dephosphorylation
cilium organization
DNA recombination
cilium assembly
regulation of cell cycle phase transition
double-strand break repair
mitotic nuclear division
meiotic cell cycle process
DNA replication
positive regulation of cell cycle process
negative regulation of mitotic cell cycle
meiotic nuclear division
synaptic vesicle cycle
negative regulation of cell cycle process
sister chromatid segregation
neurotransmitter secretion
regulation of synaptic vesicle cycle
double-strand break repair via homologous recombination
recombinational repair
microtubule organizing center organization
meiosis I cell cycle process
meiotic chromosome segregation
meiosis I
regulation of synaptic vesicle transport
chromosome organization involved in meiotic cell cycle
regulation of G2/M transition of mitotic cell cycle
homologous chromosome segregation
male meiotic nuclear division
sister chromatid cohesion
synapsis
0.01 0.02 0.03 0.04
GeneRatio
p.adjust
2.5e-07
5.0e-07
7.5e-07
Count
30
60
90
120
150
C
Cluster 3
negative regulation of immune system process
regulation of immune effector process
positive regulation of cytokine production
positive regulation of defense response
T cell activation
regulation of innate immune response
interspecies interaction between organisms
extracellular structure organization
symbiont process
regulation of cell-cell adhesion
regulation of multi-organism process
positive regulation of cell adhesion
regulation of inflammatory response
extracellular matrix organization
cell-substrate adhesion
cytokine-mediated signaling pathway
negative regulation of response to external stimulus
response to virus
leukocyte cell-cell adhesion
positive regulation of innate immune response
regulation of leukocyte cell-cell adhesion
connective tissue development
viral process
regulation of leukocyte mediated immunity
positive regulation of immune effector process
cytokine secretion
defense response to virus
negative regulation of defense response
myeloid leukocyte activation
regulation of cytokine secretion
activation of innate immune response
cell killing
response to interferon-gamma
negative regulation of immune response
leukocyte mediated cytotoxicity
pattern recognition receptor signaling pathway
innate immune response-activating signal transduction
cellular response to interferon-gamma
T cell mediated immunity
regulation of cell killing
response to interferon-beta
regulation of leukocyte mediated cytotoxicity
antigen processing and presentation
cellular response to interferon-beta
negative regulation of innate immune response
antigen processing and presentation of peptide antigen
collagen fibril organization
antigen processing and presentation of peptide antigen via MHC class I
antigen processing and presentation of endogenous peptide antigen
antigen processing and presentation of endogenous peptide antigen via MHC class I
0.01 0.02 0.03 0.04
GeneRatio
Count
20
40
60
80
p.adjust
2.5e-06
5.0e-06
7.5e-06
D
Cluster 4
ncRNA metabolic process
ribonucleoprotein complex biogenesis
ncRNA processing
proteasomal protein catabolic process
positive regulation of neuron differentiation
positive regulation of kinase activity
regulation of mitotic cell cycle
positive regulation of cell projection organization
covalent chromatin modification
negative regulation of phosphorylation
small GTPase mediated signal transduction
positive regulation of catabolic process
histone modification
regulation of cellular amide metabolic process
ribosome biogenesis
regulation of protein catabolic process
regulation of apoptotic signaling pathway
epithelial tube morphogenesis
negative regulation of protein phosphorylation
regulation of chromosome organization
Ras protein signal transduction
autophagy
process utilizing autophagic mechanism
regulation of binding
regulation of translation
peptidyl-lysine modification
positive regulation of neuron projection development
negative regulation of nervous system development
methylation
positive regulation of cellular catabolic process
cellular component disassembly
mitotic cell cycle phase transition
positive regulation of proteolysis
negative regulation of neurogenesis
rRNA metabolic process
macromolecule methylation
cellular response to peptide
connective tissue development
rRNA processing
protein localization to nucleus
morphogenesis of a branching structure
regulation of protein modification by small protein conjugation or removal
response to endoplasmic reticulum stress
regulation of proteolysis involved in cellular protein catabolic process
neural tube development
morphogenesis of embryonic epithelium
epithelial tube formation
embryonic epithelial tube formation
ribosomal large subunit biogenesis
ribosomal small subunit biogenesis
0.01 0.02 0.03 0.04
GeneRatio
p.adjust
5.0e-07
1.0e-06
1.5e-06
2.0e-06
Count
30
60
90
120
150

**fig. S5**

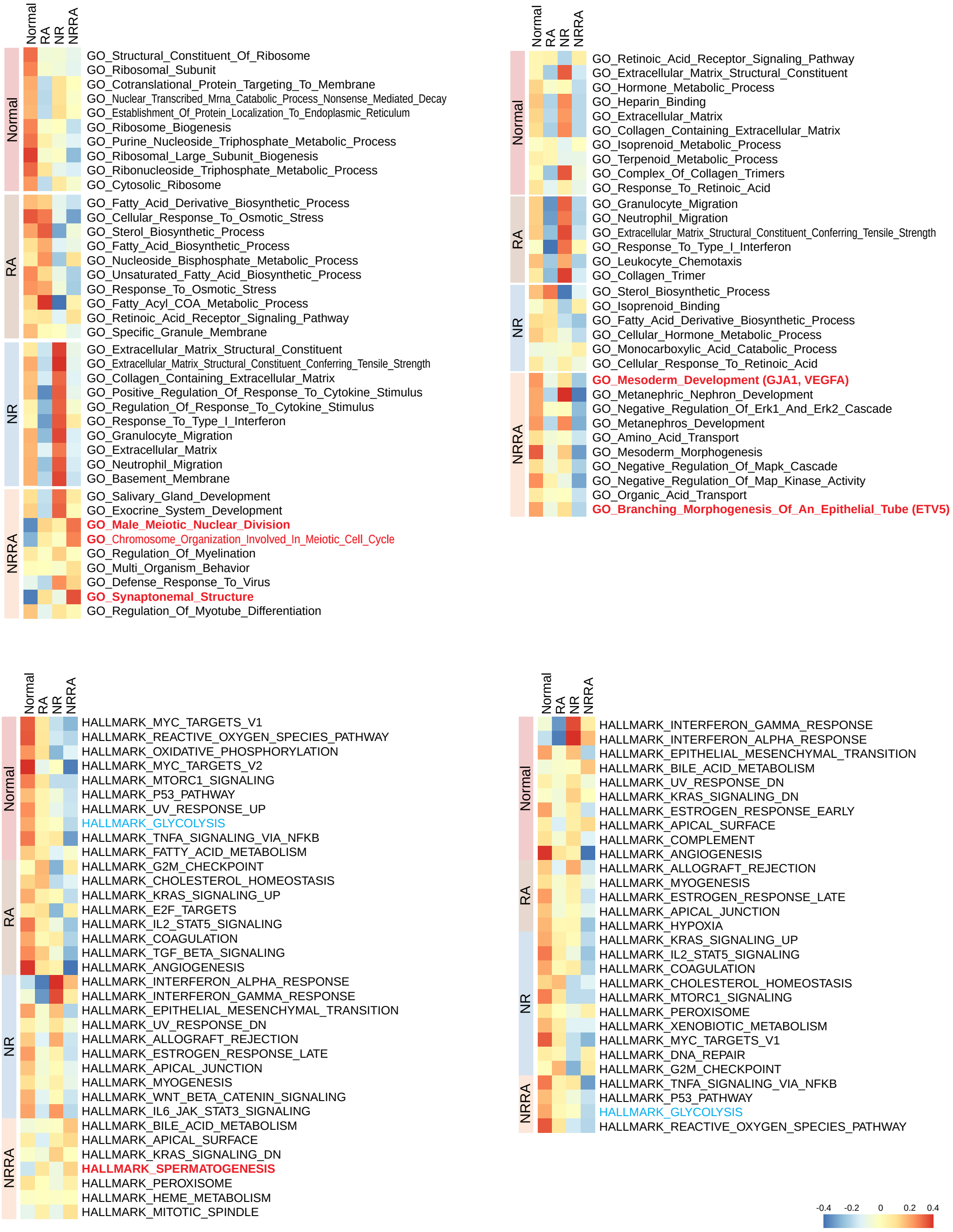
Normal
RA
NR
NRRA
GO_Structural_Constituent_Of_Ribosome
GO_Ribosomal_Subunit
GO_Cotranslational_Protein_Targeting_To_Membrane
GO_Nuclear_Transcribed_Mrna_Catabolic_Process_Nonsense_Mediated_Decay
GO_Establishment_Of_Protein_Localization_To_Endoplasmic_Reticulum
GO_Ribosome_Biogenesis
GO_Purine_Nucleoside_Triphosphate_Metabolic_Process
GO_Ribosomal_Large_Subunit_Biogenesis
GO_Ribonucleoside_Triphosphate_Metabolic_Process
GO_Cytosolic_Ribosome
GO_Fatty_Acid_Derivative_Biosynthetic_Process
GO_Cellular_Response_To_Osmotic_Stress
GO_Sterol_Biosynthetic_Process
GO_Fatty_Acid_Biosynthetic_Process
GO_Nucleoside_Bisphosphate_Metabolic_Process
GO_Unsaturated_Fatty_Acid_Biosynthetic_Process
GO_Response_To_Osmotic_Stress
GO_Fatty_Acyl_COA_Metabolic_Process
GO_Retinoic_Acid_Receptor_Signaling_Pathway
GO_Specific_Granule_Membrane
GO_Extracellular_Matrix_Structural_Constituent
GO_Extracellular_Matrix_Structural_Constituent_Conferring_Tensile_Strength
GO_Collagen_Containing_Extracellular_Matrix
GO_Positive_Regulation_Of_Response_To_Cytokine_Stimulus
GO_Regulation_Of_Response_To_Cytokine_Stimulus
GO_Response_To_Type_I_Interferon
GO_Granulocyte_Migration
GO_Extracellular_Matrix
GO_Neutrophil_Migration
GO_Basement_Membrane
GO_Salivary_Gland_Development
GO_Exocrine_System_Development
GO_Male_Meiotic_Nuclear_Division
GO_Chromosome_Organization_Involved_In_Meiotic_Cell_Cycle
GO_Regulation_Of_Myelination
GO_Multi_Organism_Behavior
GO_Defense_Response_To_Virus
GO_Synaptonemal_Structure
GO_Regulation_Of_Myotube_Differentiation
Normal
RA
NR
NRRA
GO_Retinoic_Acid_Receptor_Signaling_Pathway
GO_Extracellular_Matrix_Structural_Constituent
GO_Hormone_Metabolic_Process
GO_Heparin_Binding
GO_Extracellular_Matrix
GO_Collagen_Containing_Extracellular_Matrix
GO_Isoprenoid_Metabolic_Process
GO_Terpenoid_Metabolic_Process
GO_Complex_Of_Collagen_Trimers
GO_Response_To_Retinoic_Acid
GO_Granulocyte_Migration
GO_Neutrophil_Migration
GO_Extracellular_Matrix_Structural_Constituent_Conferring_Tensile_Strength
GO_Response_To_Type_I_Interferon
GO_Leukocyte_Chemotaxis
GO_Collagen_Trimer
GO_Sterol_Biosynthetic_Process
GO_Isoprenoid_Binding
GO_Fatty_Acid_Derivative_Biosynthetic_Process
GO_Cellular_Hormone_Metabolic_Process
GO_Monocarboxylic_Acid_Catabolic_Process
GO_Cellular_Response_To_Retinoic_Acid
GO_Mesoderm_Development (GJA1, VEGFA)
GO_Metanephric_Nephron_Development
GO_Negative_Regulation_Of_Erk1_And_Erk2_Cascade
GO_Metanephros_Development
GO_Amino_Acid_Transport
GO_Mesoderm_Morphogenesis
GO_Negative_Regulation_Of_Mapk_Cascade
GO_Negative_Regulation_Of_Map_Kinase_Activity
GO_Organic_Acid_Transport
GO_Branching_Morphogenesis_Of_An_Epithelial_Tube (ETV5)
Normal
RA
NR
NRRA
HALLMARK_MYC_TARGETS_V1
HALLMARK_REACTIVE_OXYGEN_SPECIES_PATHWAY
HALLMARK_OXIDATIVE_PHOSPHORYLATION
HALLMARK_MYC_TARGETS_V2
HALLMARK_MTORC1_SIGNALING
HALLMARK_P53_PATHWAY
HALLMARK_UV_RESPONSE_UP
HALLMARK_GLYCOLYSIS
HALLMARK_TNFA_SIGNALING_VIA_NFKB
HALLMARK_FATTY_ACID_METABOLISM
HALLMARK_G2M_CHECKPOINT
HALLMARK_CHOLESTEROL_HOMEOSTASIS
HALLMARK_KRAS_SIGNALING_UP
HALLMARK_E2F_TARGETS
HALLMARK_IL2_STAT5_SIGNALING
HALLMARK_COAGULATION
HALLMARK_TGF_BETA_SIGNALING
HALLMARK_ANGIOGENESIS
HALLMARK_INTERFERON_ALPHA_RESPONSE
HALLMARK_INTERFERON_GAMMA_RESPONSE
HALLMARK_EPITHELIAL_MESENCHYMAL_TRANSITION
HALLMARK_UV_RESPONSE_DN
HALLMARK_ALLOGRAFT_REJECTION
HALLMARK_ESTROGEN_RESPONSE_LATE
HALLMARK_APICAL_JUNCTION
HALLMARK_MYOGENESIS
HALLMARK_WNT_BETA_CATENIN_SIGNALING
HALLMARK_IL6_JAK_STAT3_SIGNALING
HALLMARK_BILE_ACID_METABOLISM
HALLMARK_APICAL_SURFACE
HALLMARK_KRAS_SIGNALING_DN
HALLMARK_SPERMATOGENESIS
HALLMARK_PEROXISOME
HALLMARK_HEME_METABOLISM
HALLMARK_MITOTIC_SPINDLE
Normal
RA
NR
NRRA
HALLMARK_INTERFERON_GAMMA_RESPONSE
HALLMARK_INTERFERON_ALPHA_RESPONSE
HALLMARK_EPITHELIAL_MESENCHYMAL_TRANSITION
HALLMARK_BILE_ACID_METABOLISM
HALLMARK_UV_RESPONSE_DN
HALLMARK_KRAS_SIGNALING_DN
HALLMARK_ESTROGEN_RESPONSE_EARLY
HALLMARK_APICAL_SURFACE
HALLMARK_COMPLEMENT
HALLMARK_ANGIOGENESIS
HALLMARK_ALLOGRAFT_REJECTION
HALLMARK_MYOGENESIS
HALLMARK_ESTROGEN_RESPONSE_LATE
HALLMARK_APICAL_JUNCTION
HALLMARK_HYPOXIA
HALLMARK_KRAS_SIGNALING_UP
HALLMARK_IL2_STAT5_SIGNALING
HALLMARK_COAGULATION
HALLMARK_CHOLESTEROL_HOMEOSTASIS
HALLMARK_MTORC1_SIGNALING
HALLMARK_PEROXISOME
HALLMARK_XENOBIOTIC_METABOLISM
HALLMARK_MYC_TARGETS_V1
HALLMARK_DNA_REPAIR
HALLMARK_G2M_CHECKPOINT
HALLMARK_TNFA_SIGNALING_VIA_NFKB
HALLMARK_P53_PATHWAY
HALLMARK_GLYCOLYSIS
HALLMARK_REACTIVE_OXYGEN_SPECIES_PATHWAY
-0.4
-0.2
0
0.2
0.4

### fig. S6

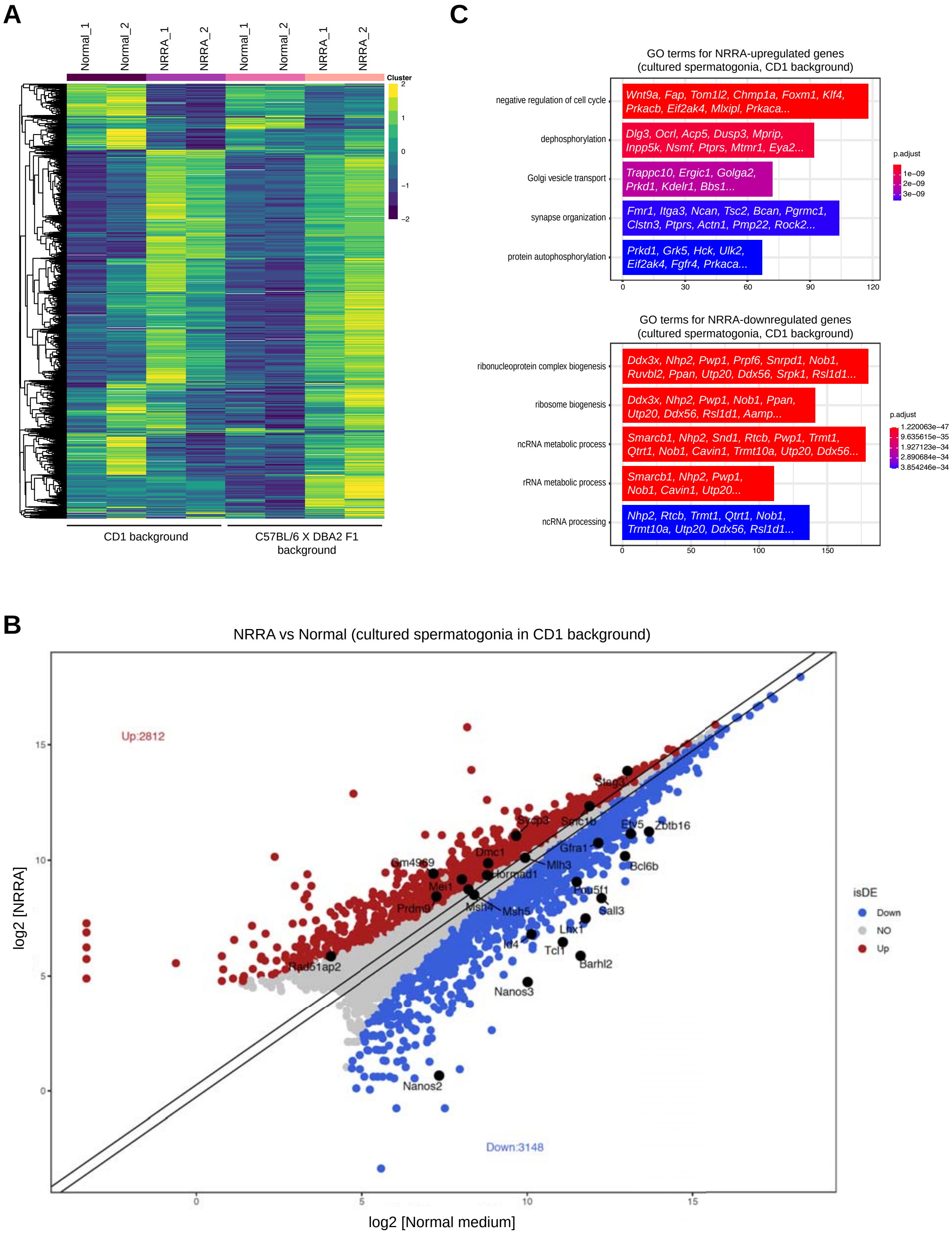
A
Normal_1
Normal_2
NRRA_1
NRRA_2
Normal_1
Normal_2
NRRA_1
NRRA_2
Cluster
2
1
0
−1
−2
CD1 background
C57BL/6 X DBA2 F1 background
C
GO terms for NRRA-upregulated genes (cultured spermatogonia, CD1 background)
negative regulation of cell cycle
Wnt9a, Fap, Tom1l2, Chmp1a, Foxm1, Klf4, Prkacb, Eif2ak4, Mlxipl, Prkaca...
dephosphorylation
Dlg3, Ocrl, Acp5, Dusp3, Mprip, Inpp5k, Nsmf, Ptprs, Mtmr1, Eya2...
Golgi vesicle transport
Trappc10, Ergic1, Golga2, Prkd1, Kdelr1, Bbs1...
synapse organization
Fmr1, Itga3, Ncan, Tsc2, Bcan, Pgrmc1, Clstn3, Ptprs, Actn1, Pmp22, Rock2...
protein autophosphorylation
Prkd1, Grk5, Hck, Ulk2, Eif2ak4, Fgfr4, Prkaca...
0
30
60
90
120
p.adjust
1e−09
2e−09
3e−09
GO terms for NRRA-downregulated genes (cultured spermatogonia, CD1 background)
ribonucleoprotein complex biogenesis
Ddx3x, Nhp2, Pwp1, Prpf6, Snrpd1, Nob1, Ruvbl2, Ppan, Utp20, Ddx56, Srpk1, Rsl1d1...
ribosome biogenesis
Ddx3x, Nhp2, Pwp1, Nob1, Ppan, Utp20, Ddx56, Rsl1d1, Aamp...
ncRNA metabolic process
Smarcb1, Nhp2, Snd1, Rtcb, Pwp1, Trmt1, Qtrt1, Nob1, Cavin1, Trmt10a, Utp20, Ddx56...
rRNA metabolic process
Smarcb1, Nhp2, Pwp1, Nob1, Cavin1, Utp20...
ncRNA processing
Nhp2, Rtcb, Trmt1, Qtrt1, Nob1, Trmt10a, Utp20, Ddx56, Rsl1d1...
0
50
100
150
p.adjust
1.220063e−47
9.635615e−35
1.927123e−34
2.890684e−34
3.854246e−34
B
NRRA vs Normal (cultured spermatogonia in CD1 background)
Up:2812
Down:3148
log2 [NRRA]
log2 [Normal medium]
0
5
10
15
Stag3
Sycp3
Smc1b
Etv5
Zbtb16
Gfra1
Bcl6b
Dmc1
Gm4969
Hormad1
Mlh3
Mei1
Pou5f1
Sall3
Prdm9
Msh4
Msh5
Lhx1
Id4
Tcl1
Rad51ap2
Barhl2
Nanos3
Nanos2
isDE
Down
NO
Up

### fig. S7

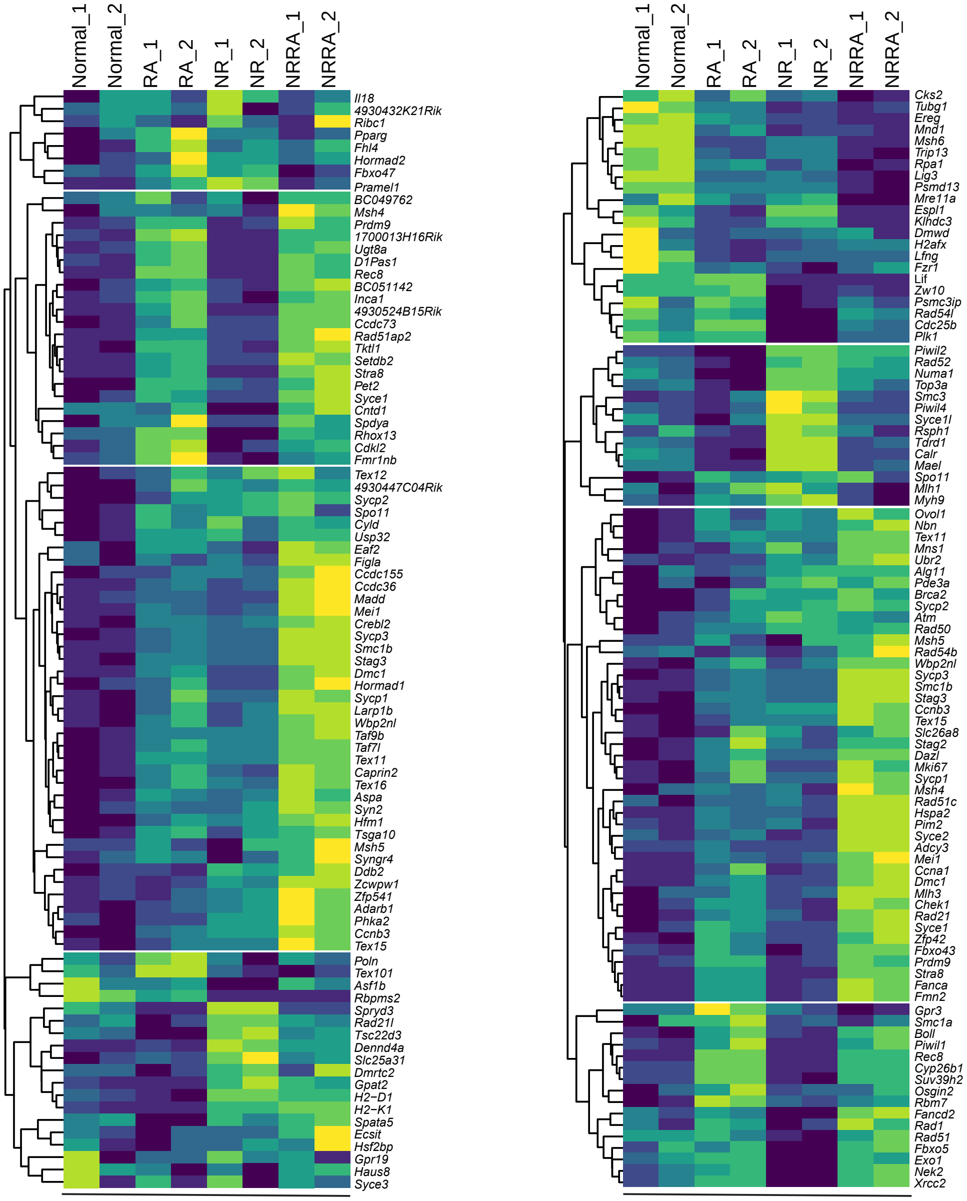


Early mouse meiosis (104 genes)
(Soh *et al*, 2015)

Early human meiosis (107 genes)
(Guo *et al*, 2017)

#### fig. S8

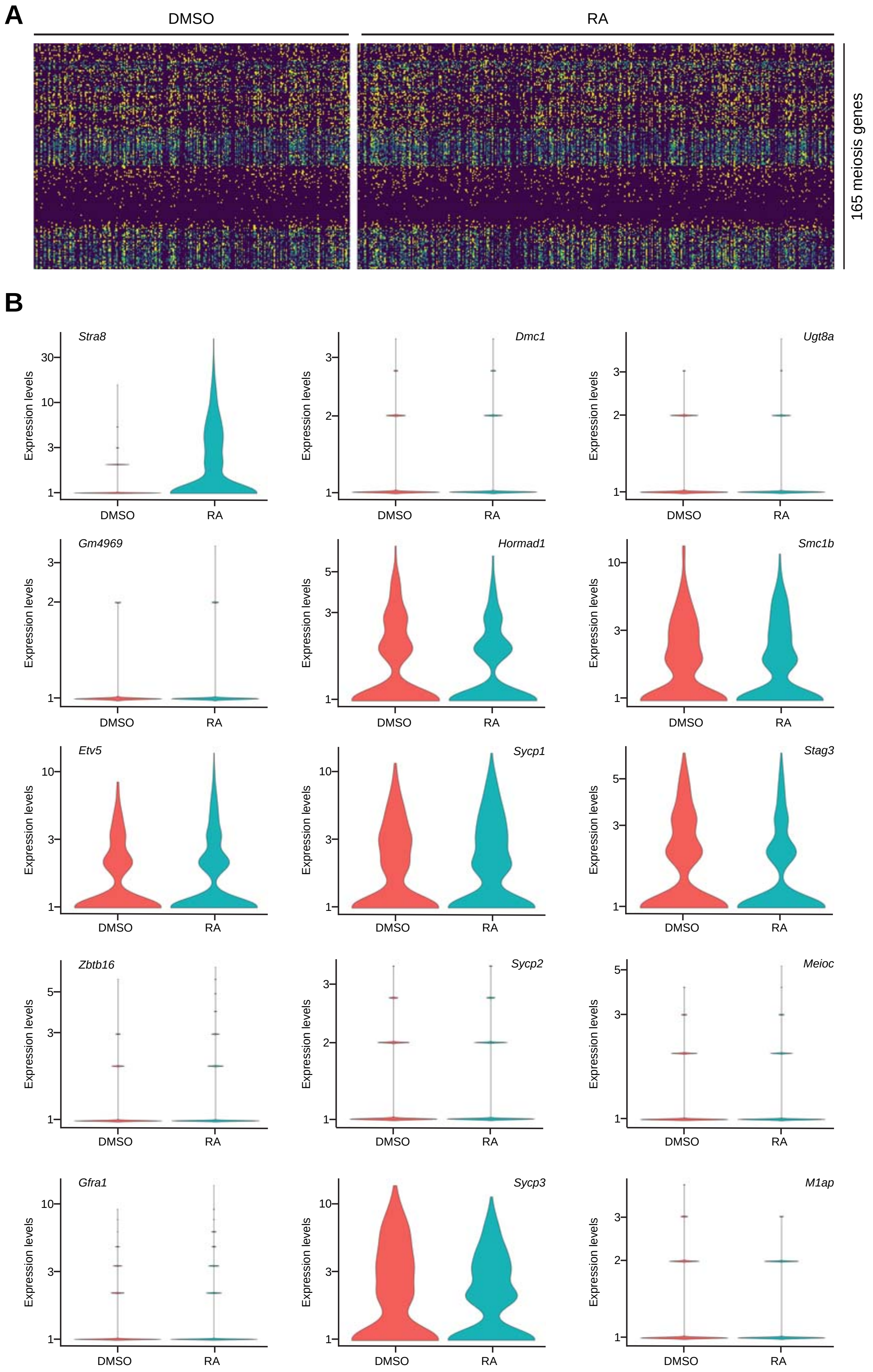
A
DMSO
RA
165 meiosis genes
B
Stra8
Dmc1
Ugt8a
Gm4969
Hormad1
Smc1b
Etv5
Sycp1
Stag3
Zbtb16
Sycp2
Meioc
Gfra1
Sycp3
M1ap
Expression levels
DMSO
RA

**fig. S9**

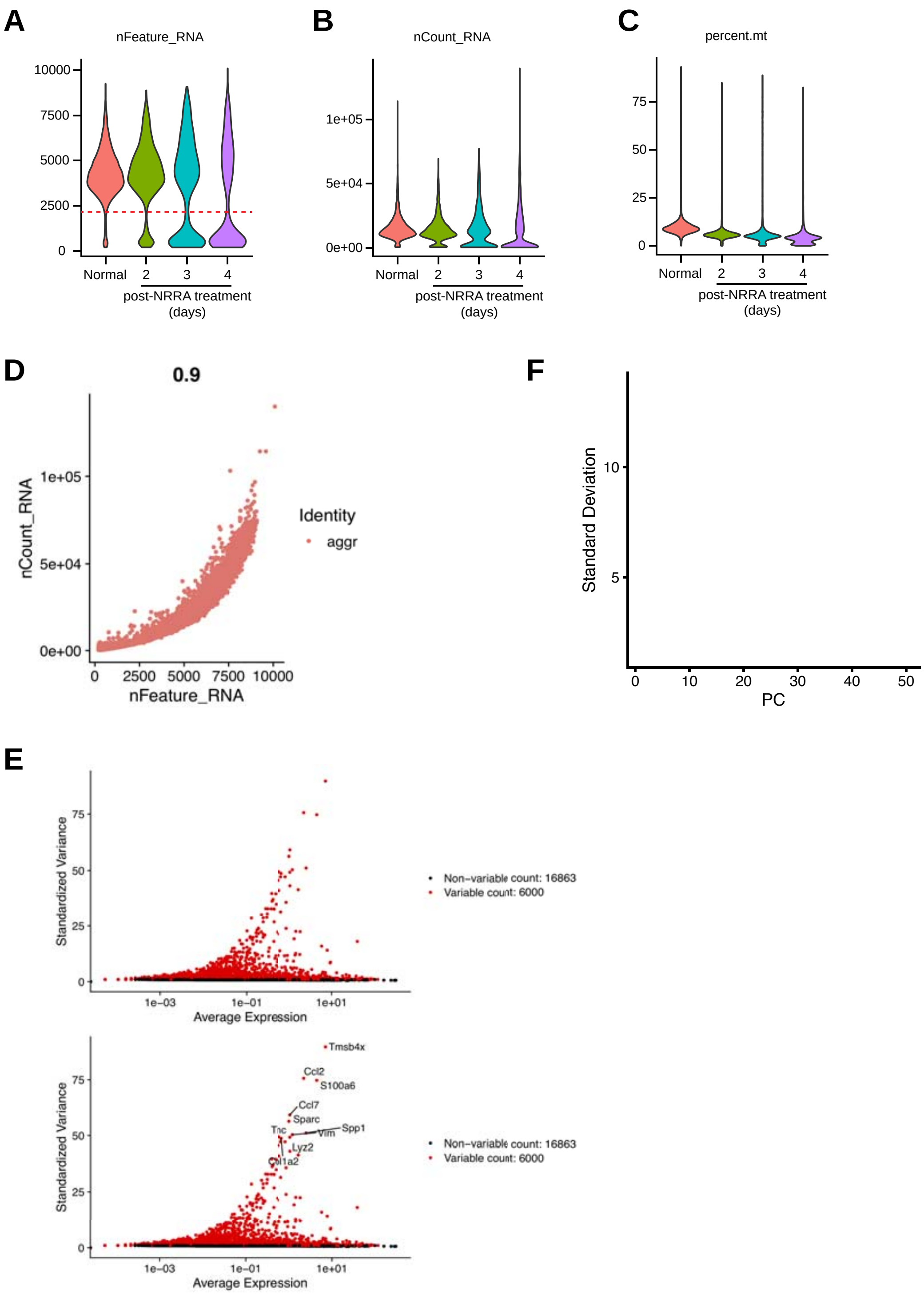
A
nFeature_RNA
10000
7500
5000
2500
0
Normal
2
3
4
post-NRRA treatment
(days)
B
nCount_RNA
1e+05
5e+04
0e+00
Normal
2
3
4
post-NRRA treatment
(days)
C
percent.mt
75
50
25
0
Normal
2
3
4
post-NRRA treatment
(days)
D
0.9
nCount_RNA
1e+05
5e+04
0e+00
0
2500
5000
7500
10000
nFeature_RNA
Identity
aggr
F
Standard Deviation
10
5
0
10
20
30
40
50
PC
E
Standardized Variance
75
50
25
0
1e−03
1e−01
1e+01
Average Expression
Non−variable count: 16863
Variable count: 6000
Tmsb4x
Ccl2
S100a6
Ccl7
Sparc
Tnc
Vim
Spp1
Lyz2
Col1a2
Standardized Variance
75
50
25
0
1e−03
1e−01
1e+01
Average Expression
Non−variable count: 16863
Variable count: 6000

fig. S10

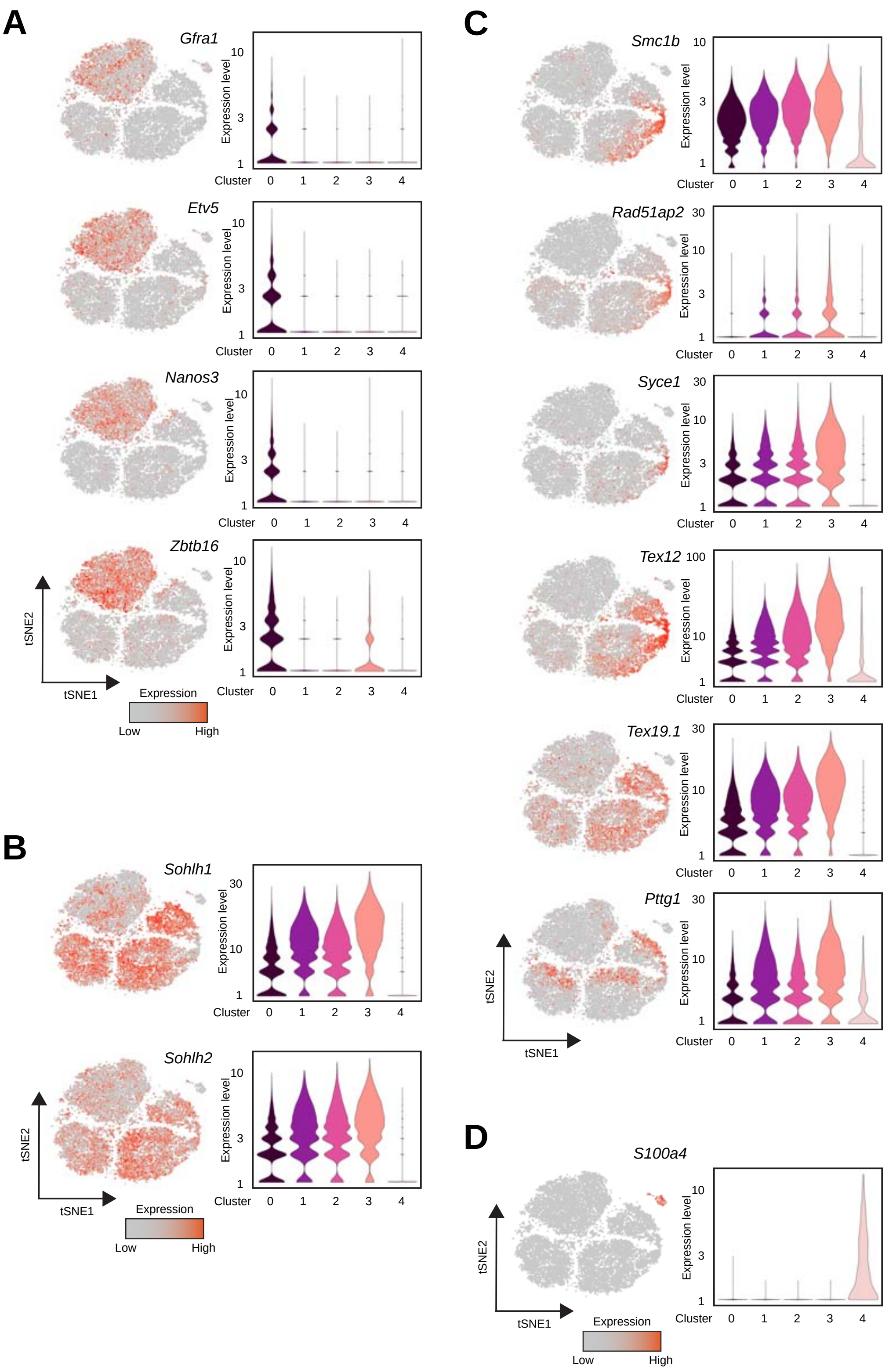
A
Gfra1
Etv5
Nanos3
Zbtb16
Expression level
Cluster 0 1 2 3 4
tSNE1
tSNE2
Expression
Low
High
B
Sohlh1
Sohlh2
C
Smc1b
Rad51ap2
Syce1
Tex12
Tex19.1
Pttg1
D
S100a4

### fig. S11

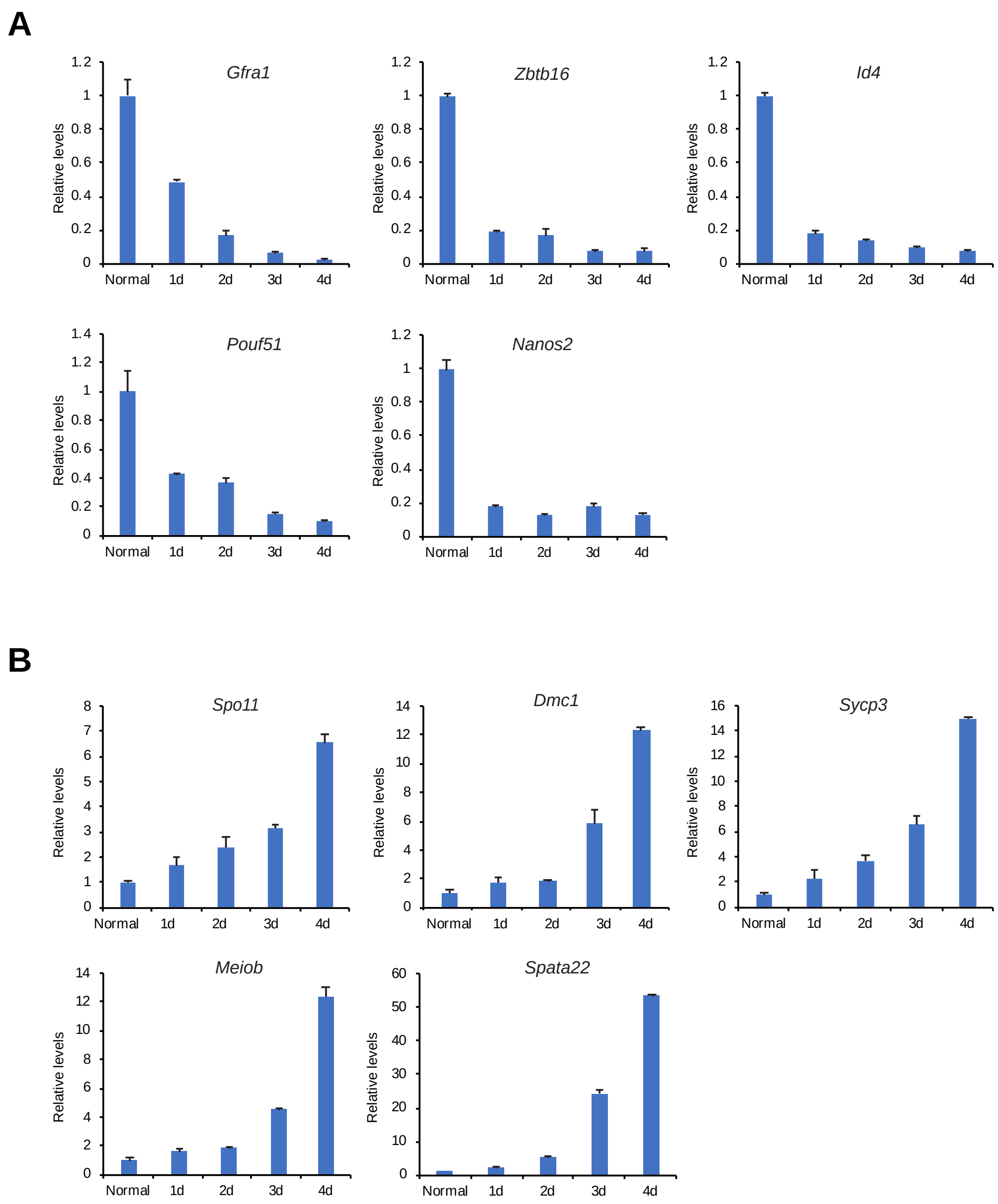

A
Gfra1
Zbtb16
Id4
Pouf51
Nanos2
Relative levels
Normal
1d
2d
3d
4d
B
Spo11
Dmc1
Sycp3
Meiob
Spata22

fig S12A

Cluster 0

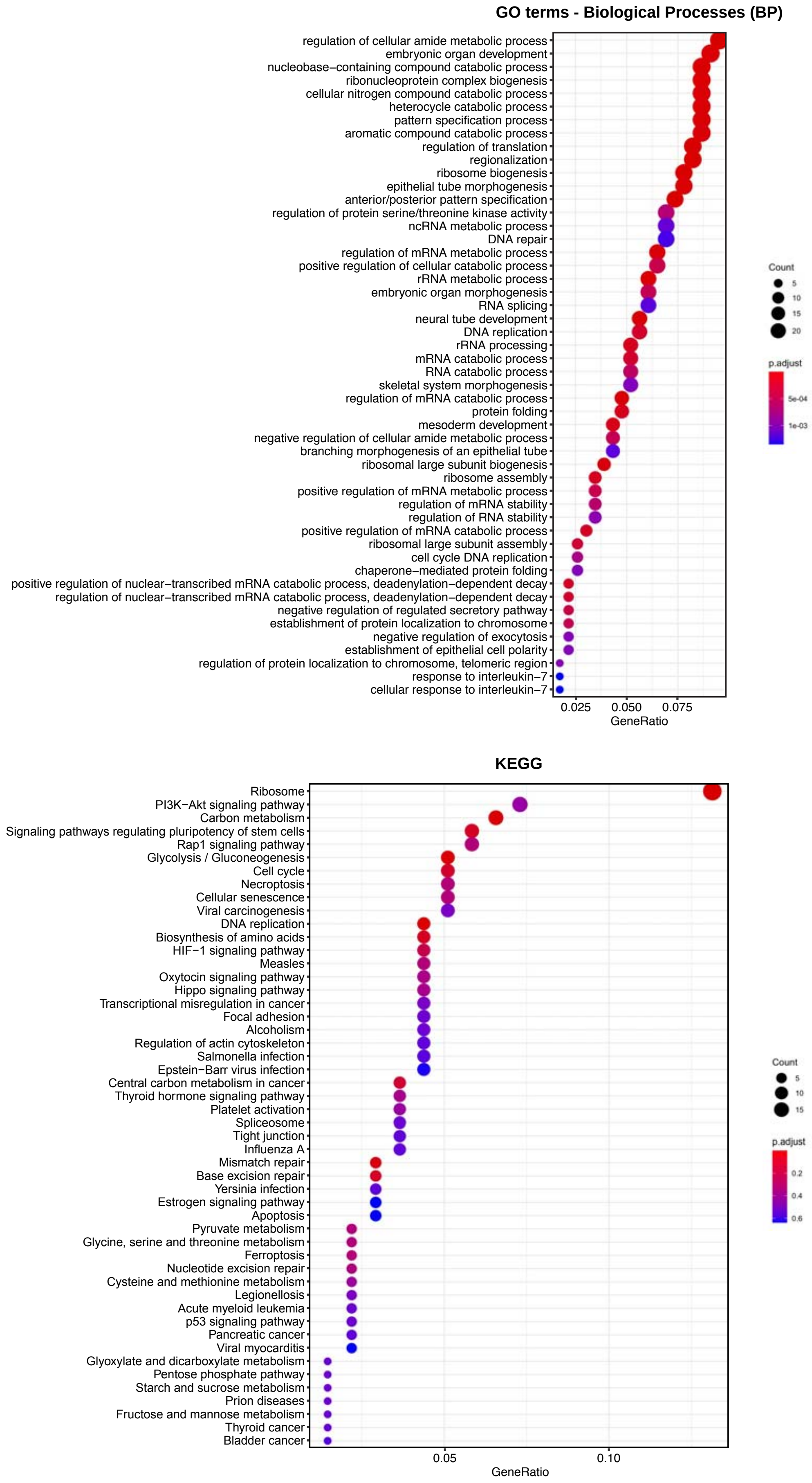

### fig S12B

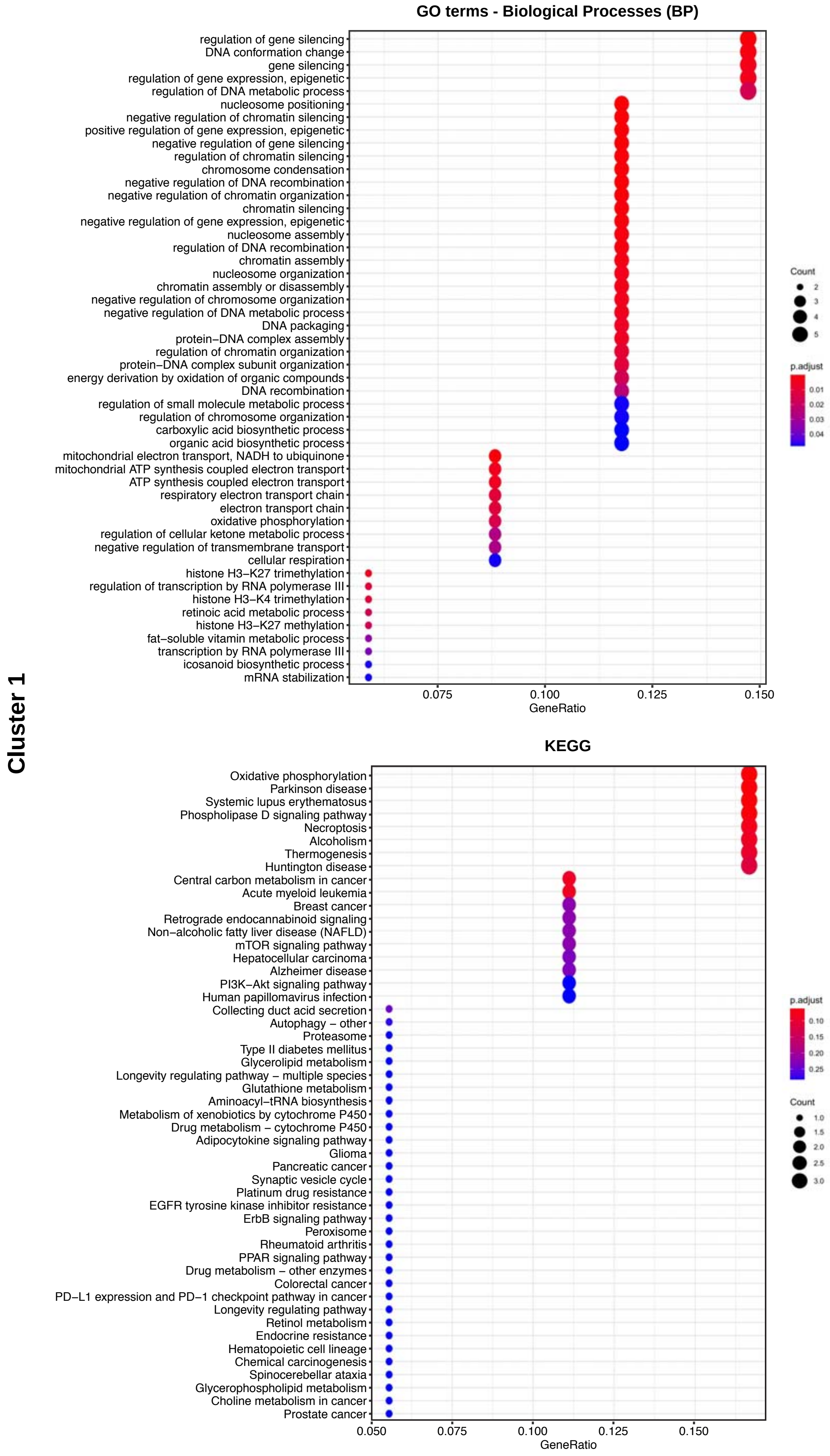

Cluster 1
GO terms - Biological Processes (BP)
regulation of gene silencing
DNA conformation change
gene silencing
regulation of gene expression, epigenetic
regulation of DNA metabolic process
nucleosome positioning
negative regulation of chromatin silencing
positive regulation of gene expression, epigenetic
negative regulation of gene silencing
regulation of chromatin silencing
chromosome condensation
negative regulation of DNA recombination
negative regulation of chromatin organization
chromatin silencing
negative regulation of gene expression, epigenetic
nucleosome assembly
regulation of DNA recombination
chromatin assembly
nucleosome organization
chromatin assembly or disassembly
negative regulation of chromosome organization
negative regulation of DNA metabolic process
DNA packaging
protein−DNA complex assembly
regulation of chromatin organization
protein−DNA complex subunit organization
energy derivation by oxidation of organic compounds
DNA recombination
regulation of small molecule metabolic process
regulation of chromosome organization
carboxylic acid biosynthetic process
organic acid biosynthetic process
mitochondrial electron transport, NADH to ubiquinone
mitochondrial ATP synthesis coupled electron transport
ATP synthesis coupled electron transport
respiratory electron transport chain
electron transport chain
oxidative phosphorylation
regulation of cellular ketone metabolic process
negative regulation of transmembrane transport
cellular respiration
histone H3−K27 trimethylation
regulation of transcription by RNA polymerase III
histone H3−K4 trimethylation
retinoic acid metabolic process
histone H3−K27 methylation
fat−soluble vitamin metabolic process
transcription by RNA polymerase III
icosanoid biosynthetic process
mRNA stabilization
0.075
0.100
0.125
0.150
GeneRatio
Count
2
3
4
5
p.adjust
0.01
0.02
0.03
0.04
KEGG
Oxidative phosphorylation
Parkinson disease
Systemic lupus erythematosus
Phospholipase D signaling pathway
Necroptosis
Alcoholism
Thermogenesis
Huntington disease
Central carbon metabolism in cancer
Acute myeloid leukemia
Breast cancer
Retrograde endocannabinoid signaling
Non−alcoholic fatty liver disease (NAFLD)
mTOR signaling pathway
Hepatocellular carcinoma
Alzheimer disease
PI3K−Akt signaling pathway
Human papillomavirus infection
Collecting duct acid secretion
Autophagy − other
Proteasome
Type II diabetes mellitus
Glycerolipid metabolism
Longevity regulating pathway − multiple species
Glutathione metabolism
Aminoacyl−tRNA biosynthesis
Metabolism of xenobiotics by cytochrome P450
Drug metabolism − cytochrome P450
Adipocytokine signaling pathway
Glioma
Pancreatic cancer
Synaptic vesicle cycle
Platinum drug resistance
EGFR tyrosine kinase inhibitor resistance
ErbB signaling pathway
Peroxisome
Rheumatoid arthritis
PPAR signaling pathway
Drug metabolism − other enzymes
Colorectal cancer
PD−L1 expression and PD−1 checkpoint pathway in cancer
Longevity regulating pathway
Retinol metabolism
Endocrine resistance
Hematopoietic cell lineage
Chemical carcinogenesis
Spinocerebellar ataxia
Glycerophospholipid metabolism
Choline metabolism in cancer
Prostate cancer
0.050
0.075
0.100
0.125
0.150
GeneRatio
p.adjust
0.10
0.15
0.20
0.25
Count
1.0
1.5
2.0
2.5
3.0

### fig. S12C

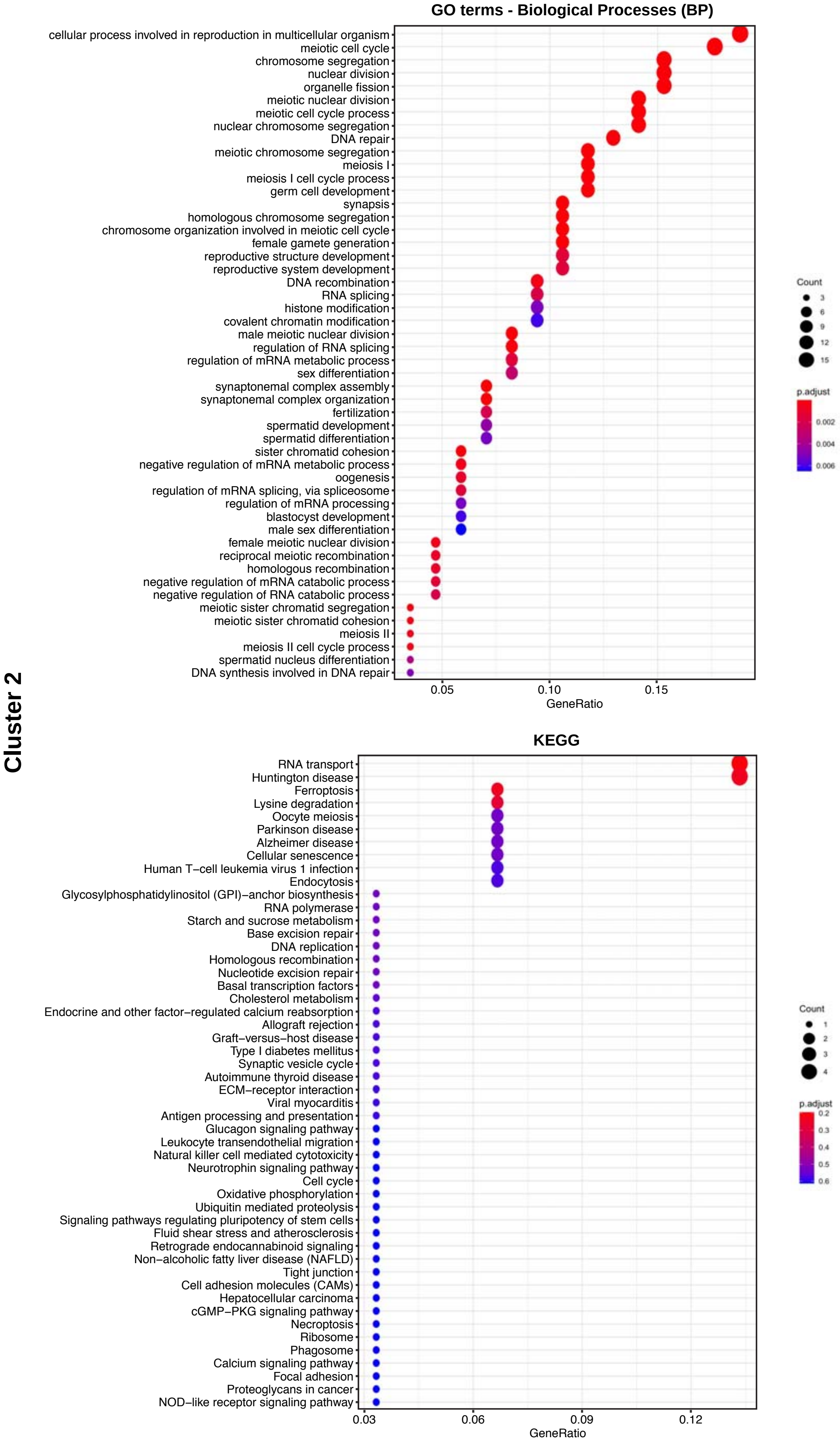

Cluster 2
GO terms - Biological Processes (BP)
cellular process involved in reproduction in multicellular organism
meiotic cell cycle
chromosome segregation
nuclear division
organelle fission
meiotic nuclear division
meiotic cell cycle process
nuclear chromosome segregation
DNA repair
meiotic chromosome segregation
meiosis I
meiosis I cell cycle process
germ cell development
synapsis
homologous chromosome segregation
chromosome organization involved in meiotic cell cycle
female gamete generation
reproductive structure development
reproductive system development
DNA recombination
RNA splicing
histone modification
covalent chromatin modification
male meiotic nuclear division
regulation of RNA splicing
regulation of mRNA metabolic process
sex differentiation
synaptonemal complex assembly
synaptonemal complex organization
fertilization
spermatid development
spermatid differentiation
sister chromatid cohesion
negative regulation of mRNA metabolic process
oogenesis
regulation of mRNA splicing, via spliceosome
regulation of mRNA processing
blastocyst development
male sex differentiation
female meiotic nuclear division
reciprocal meiotic recombination
homologous recombination
negative regulation of mRNA catabolic process
negative regulation of RNA catabolic process
meiotic sister chromatid segregation
meiotic sister chromatid cohesion
meiosis II
meiosis II cell cycle process
spermatid nucleus differentiation
DNA synthesis involved in DNA repair
0.05
0.10
0.15
GeneRatio
Count
3
6
9
12
15
p.adjust
0.002
0.004
0.006
KEGG
RNA transport
Huntington disease
Ferroptosis
Lysine degradation
Oocyte meiosis
Parkinson disease
Alzheimer disease
Cellular senescence
Human T−cell leukemia virus 1 infection
Endocytosis
Glycosylphosphatidylinositol (GPI)−anchor biosynthesis
RNA polymerase
Starch and sucrose metabolism
Base excision repair
DNA replication
Homologous recombination
Nucleotide excision repair
Basal transcription factors
Cholesterol metabolism
Endocrine and other factor−regulated calcium reabsorption
Allograft rejection
Graft−versus−host disease
Type I diabetes mellitus
Synaptic vesicle cycle
Autoimmune thyroid disease
ECM−receptor interaction
Viral myocarditis
Antigen processing and presentation
Glucagon signaling pathway
Leukocyte transendothelial migration
Natural killer cell mediated cytotoxicity
Neurotrophin signaling pathway
Cell cycle
Oxidative phosphorylation
Ubiquitin mediated proteolysis
Signaling pathways regulating pluripotency of stem cells
Fluid shear stress and atherosclerosis
Retrograde endocannabinoid signaling
Non−alcoholic fatty liver disease (NAFLD)
Tight junction
Cell adhesion molecules (CAMs)
Hepatocellular carcinoma
cGMP−PKG signaling pathway
Necroptosis
Ribosome
Phagosome
Calcium signaling pathway
Focal adhesion
Proteoglycans in cancer
NOD−like receptor signaling pathway
0.03
0.06
0.09
0.12
GeneRatio
Count
1
2
3
4
p.adjust
0.2
0.3
0.4
0.5
0.6

### fig. S12D

Cluster 3

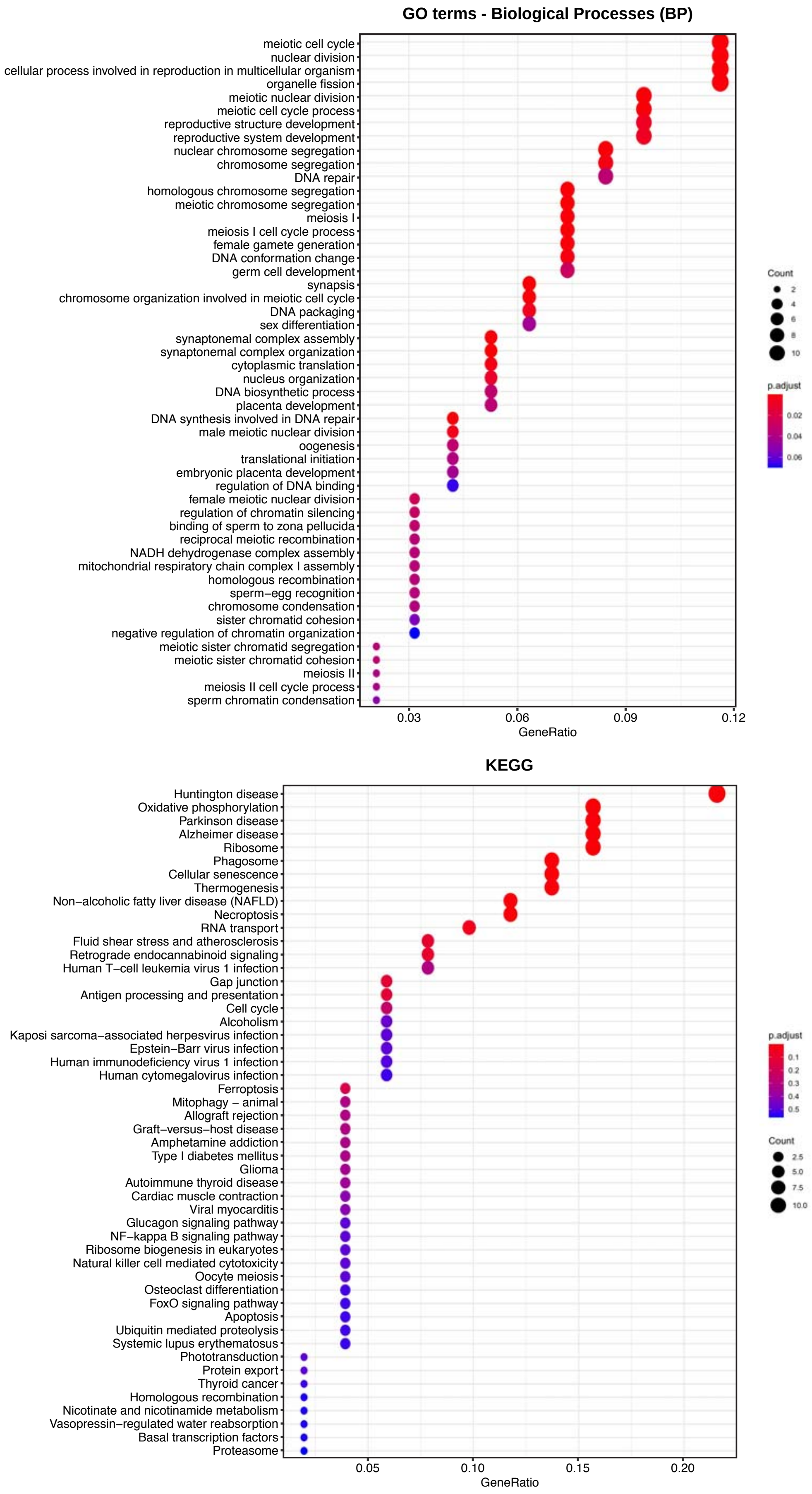

### Fig. S13

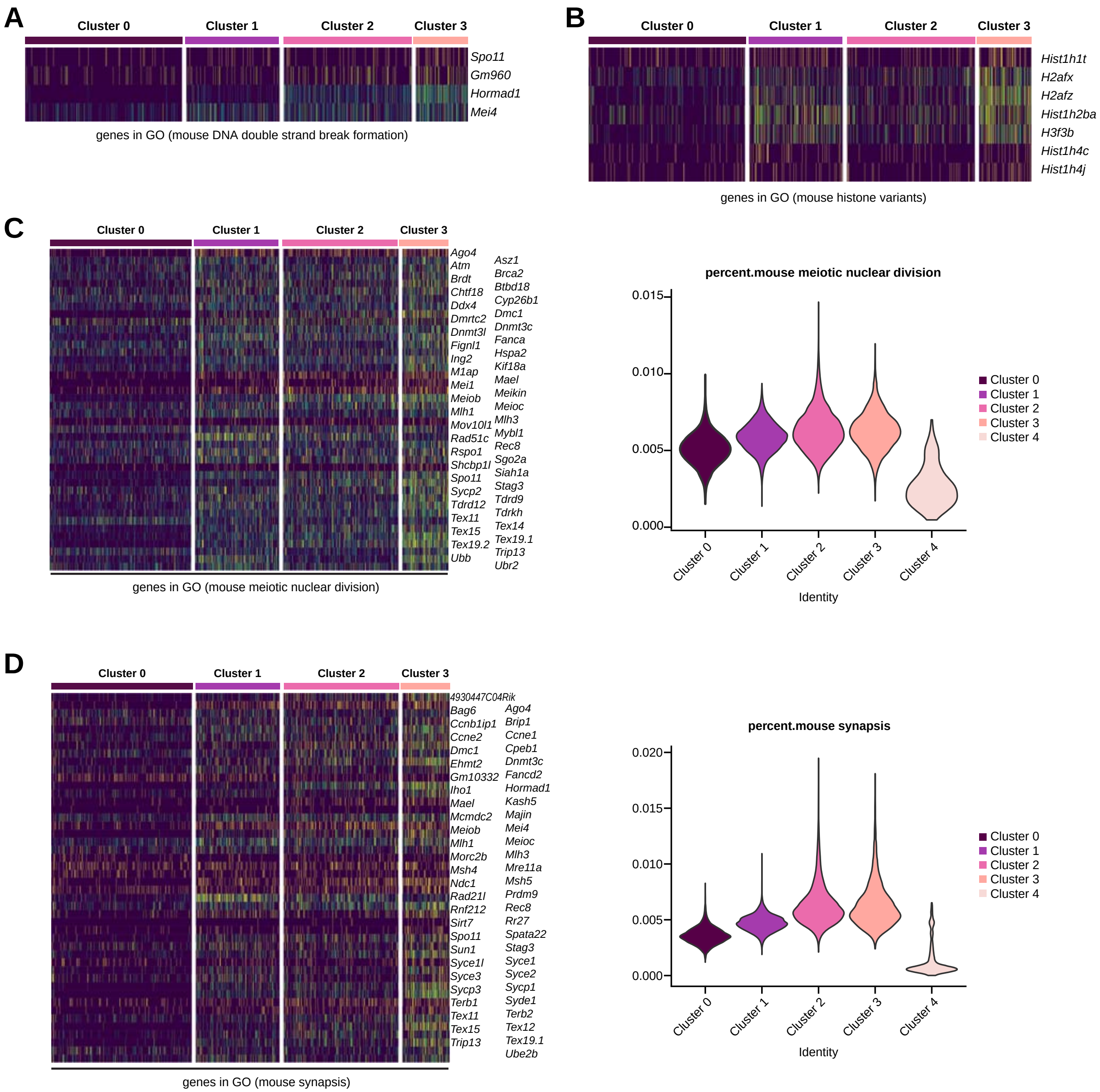
A
Cluster 0
Cluster 1
Cluster 2
Cluster 3
Spo11
Gm960
Hormad1
Mei4
genes in GO (mouse DNA double strand break formation)
B
Cluster 0
Cluster 1
Cluster 2
Cluster 3
Hist1h1t
H2afx
H2afz
Hist1h2ba
H3f3b
Hist1h4c
Hist1h4j
genes in GO (mouse histone variants)
C
Cluster 0
Cluster 1
Cluster 2
Cluster 3
Ago4
Atm
Brdt
Chtf18
Ddx4
Dmrtc2
Dnmt3l
Fignl1
Ing2
M1ap
Mei1
Meiob
Mlh1
Mov10l1
Rad51c
Rspo1
Shcbp1l
Spo11
Sycp2
Tdrd12
Tex11
Tex15
Tex19.2
Ubb
Asz1
Brca2
Btbd18
Cyp26b1
Dmc1
Dnmt3c
Fanca
Hspa2
Kif18a
Mael
Meikin
Meioc
Mlh3
Mybl1
Rec8
Sgo2a
Siah1a
Stag3
Tdrd9
Tdrkh
Tex14
Tex19.1
Trip13
Ubr2
genes in GO (mouse meiotic nuclear division)
percent.mouse meiotic nuclear division
0.015
0.010
0.005
0.000
Cluster 0
Cluster 1
Cluster 2
Cluster 3
Cluster 4
Identity
D
Cluster 0
Cluster 1
Cluster 2
Cluster 3
4930447C04Rik
Bag6
Ccnb1ip1
Ccne2
Dmc1
Ehmt2
Gm10332
Iho1
Mael
Mcmdc2
Meiob
Mlh1
Morc2b
Msh4
Ndc1
Rad21l
Rnf212
Sirt7
Spo11
Sun1
Syce1l
Syce3
Sycp3
Terb1
Tex11
Tex15
Trip13
Ago4
Brip1
Ccne1
Cpeb1
Dnmt3c
Fancd2
Hormad1
Kash5
Majin
Mei4
Meioc
Mlh3
Mre11a
Msh5
Prdm9
Rec8
Rr27
Spata22
Stag3
Syce1
Syce2
Sycp1
Syde1
Terb2
Tex12
Tex19.1
Ube2b
genes in GO (mouse synapsis)
percent.mouse synapsis
0.020
0.015
0.010
0.005
0.000
Cluster 0
Cluster 1
Cluster 2
Cluster 3
Cluster 4
Identity

### fig. S14

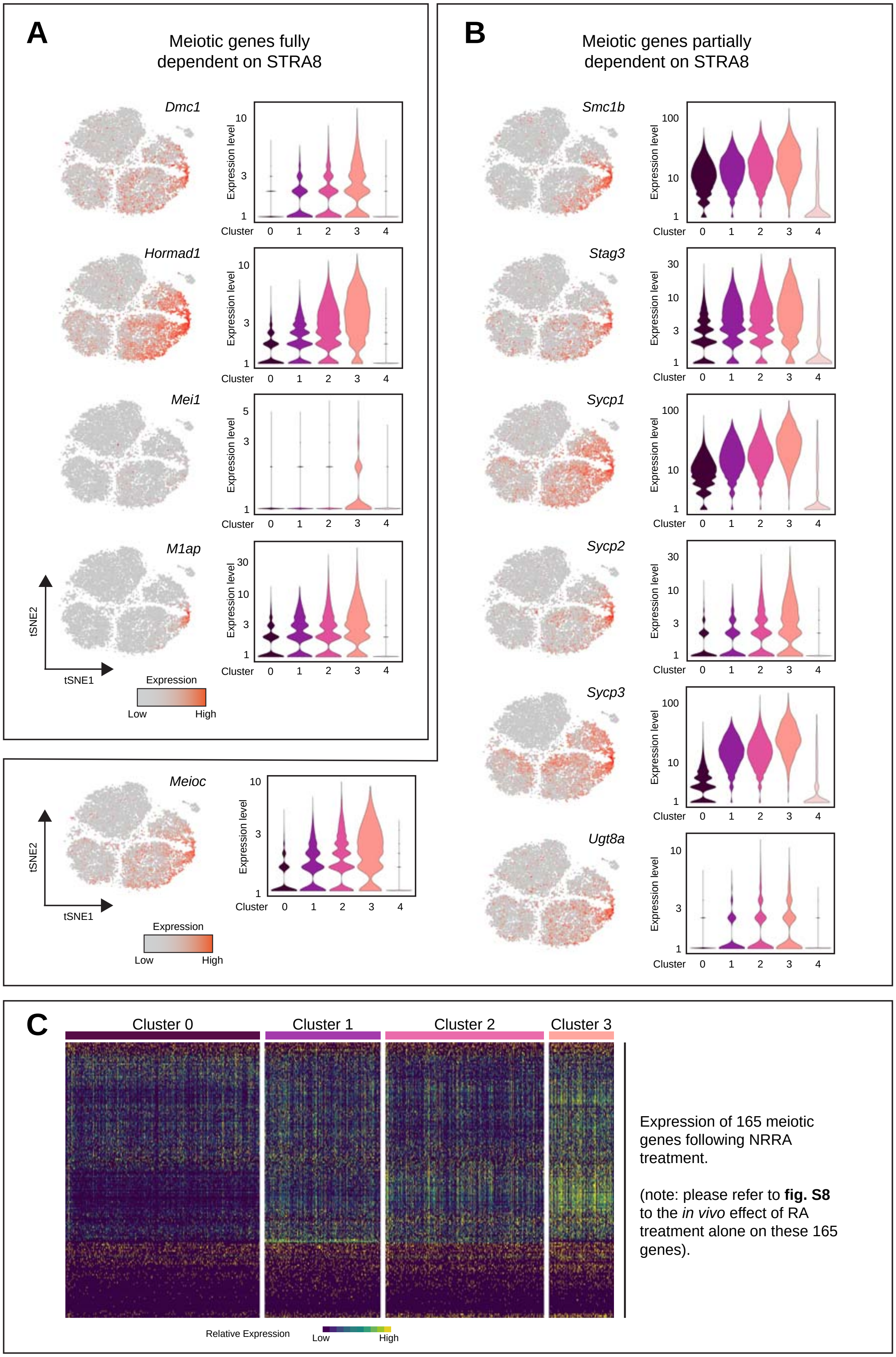
A
Meiotic genes fully dependent on STRA8
Dmc1
Hormad1
Mei1
M1ap
Expression level
Cluster
0
1
2
3
4
tSNE2
tSNE1
Expression
Low
High
Meioc
B
Meiotic genes partially dependent on STRA8
Smc1b
Stag3
Sycp1
Sycp2
Sycp3
Ugt8a
C
Cluster 0
Cluster 1
Cluster 2
Cluster 3
Relative Expression
Expression of 165 meiotic genes following NRRA treatment.
(note: please refer to fig. S8 to the in vivo effect of RA treatment alone on these 165 genes).

#### fig. S15

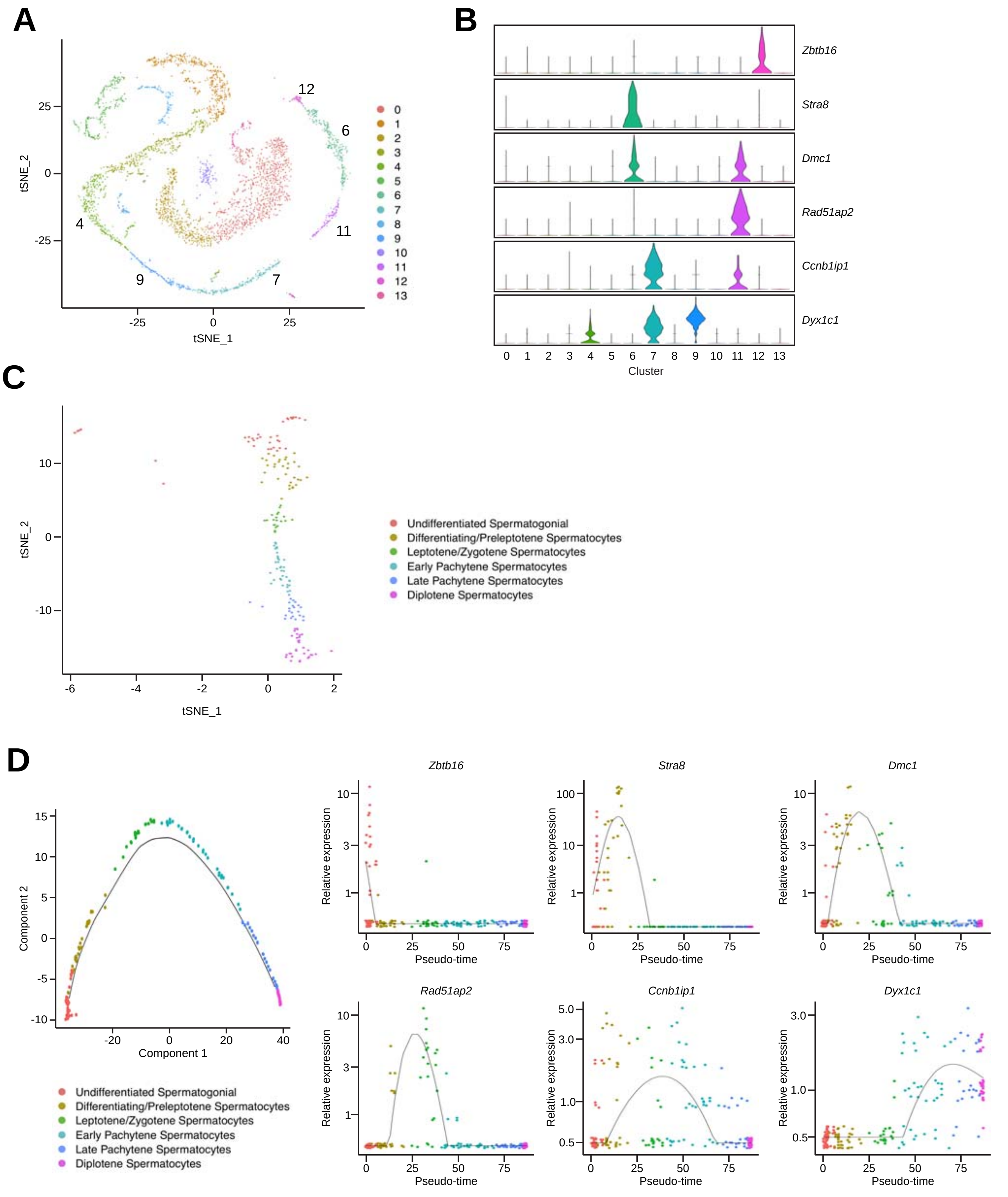

A
B
C
D
tSNE_1
tSNE_2
Zbtb16
Stra8
Dmc1
Rad51ap2
Ccnb1ip1
Dyx1c1
Cluster
Undifferentiated Spermatogonial
Differentiating/Preleptotene Spermatocytes
Leptotene/Zygotene Spermatocytes
Early Pachytene Spermatocytes
Late Pachytene Spermatocytes
Diplotene Spermatocytes
Component 1
Component 2
Pseudo-time
Relative expression

### fig. S16

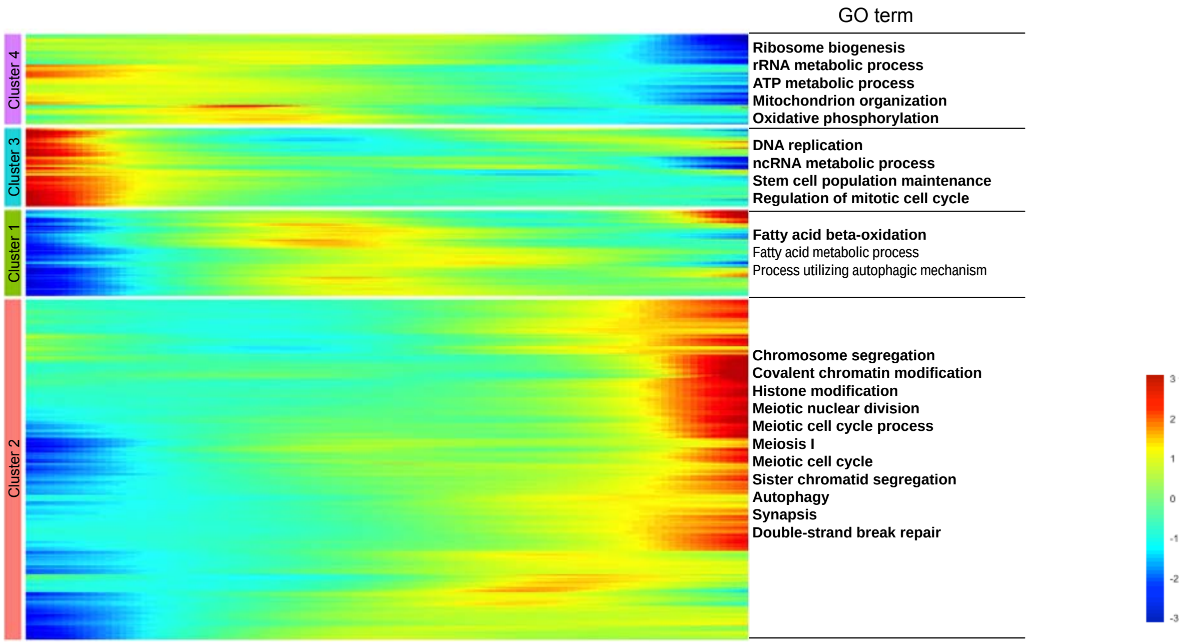
GO term
Cluster 4
Cluster 3
Cluster 1
Cluster 2
Ribosome biogenesis
rRNA metabolic process
ATP metabolic process
Mitochondrion organization
Oxidative phosphorylation
DNA replication
ncRNA metabolic process
Stem cell population maintenance
Regulation of mitotic cell cycle
Fatty acid beta-oxidation
Fatty acid metabolic process
Process utilizing autophagic mechanism
Chromosome segregation
Covalent chromatin modification
Histone modification
Meiotic nuclear division
Meiotic cell cycle process
Meiosis I
Meiotic cell cycle
Sister chromatid segregation
Autophagy
Synapsis
Double-strand break repair
3
2
1
0
-1
-2
-3

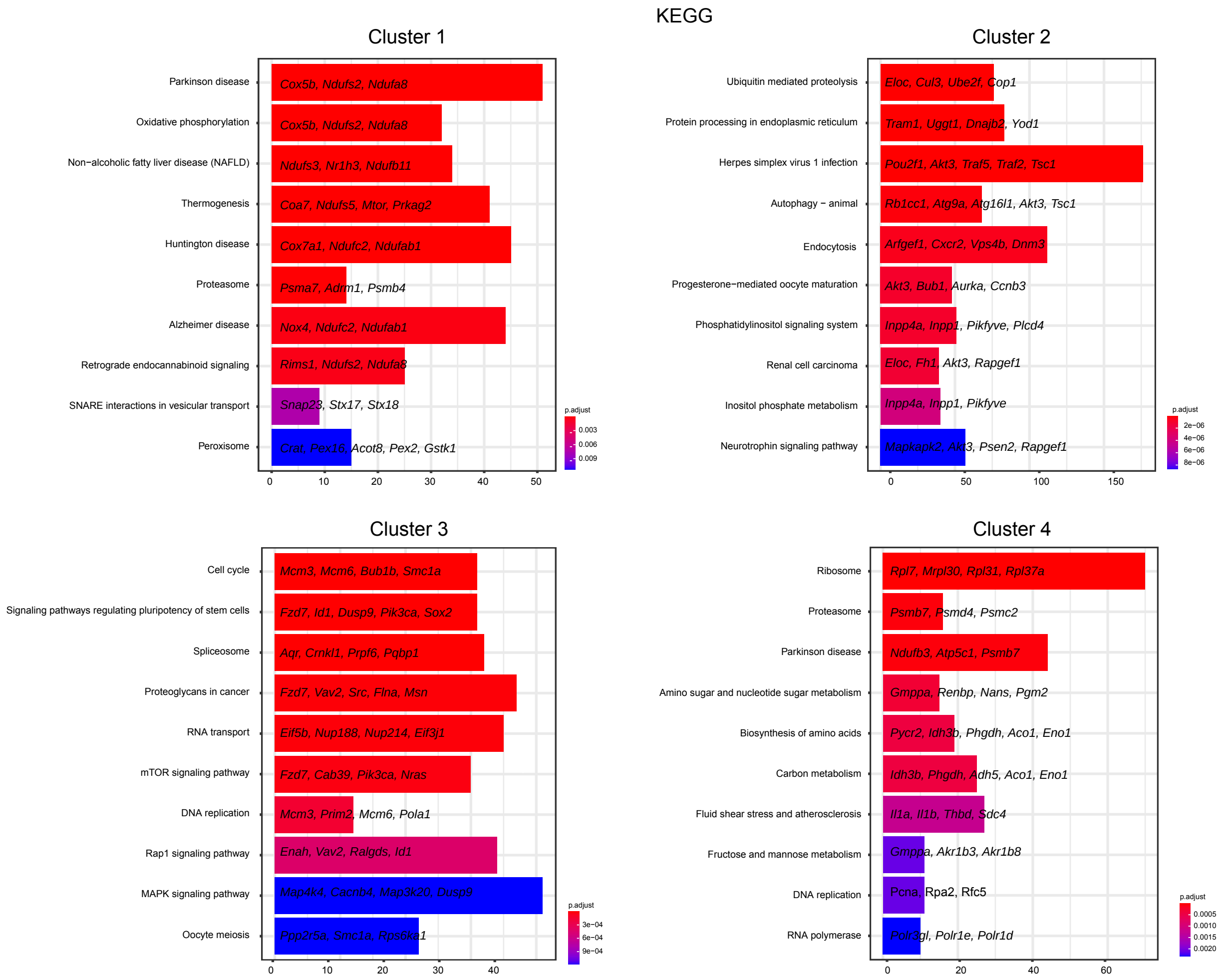
KEGG
Cluster 1
Parkinson disease
Cox5b, Ndufs2, Ndufa8
Oxidative phosphorylation
Cox5b, Ndufs2, Ndufa8
Non−alcoholic fatty liver disease (NAFLD)
Ndufs3, Nr1h3, Ndufb11
Thermogenesis
Coa7, Ndufs5, Mtor, Prkag2
Huntington disease
Cox7a1, Ndufc2, Ndufab1
Proteasome
Psma7, Adrm1, Psmb4
Alzheimer disease
Nox4, Ndufc2, Ndufab1
Retrograde endocannabinoid signaling
Rims1, Ndufs2, Ndufa8
SNARE interactions in vesicular transport
Snap23, Stx17, Stx18
Peroxisome
Crat, Pex16, Acot8, Pex2, Gstk1
0
10
20
30
40
50
p.adjust
0.003
0.006
0.009
Cluster 2
Ubiquitin mediated proteolysis
Eloc, Cul3, Ube2f, Cop1
Protein processing in endoplasmic reticulum
Tram1, Uggt1, Dnajb2, Yod1
Herpes simplex virus 1 infection
Pou2f1, Akt3, Traf5, Traf2, Tsc1
Autophagy − animal
Rb1cc1, Atg9a, Atg16l1, Akt3, Tsc1
Endocytosis
Arfgef1, Cxcr2, Vps4b, Dnm3
Progesterone−mediated oocyte maturation
Akt3, Bub1, Aurka, Ccnb3
Phosphatidylinositol signaling system
Inpp4a, Inpp1, Pikfyve, Plcd4
Renal cell carcinoma
Eloc, Fh1, Akt3, Rapgef1
Inositol phosphate metabolism
Inpp4a, Inpp1, Pikfyve
Neurotrophin signaling pathway
Mapkapk2, Akt3, Psen2, Rapgef1
0
50
100
150
p.adjust
2e−06
4e−06
6e−06
8e−06
Cluster 3
Cell cycle
Mcm3, Mcm6, Bub1b, Smc1a
Signaling pathways regulating pluripotency of stem cells
Fzd7, Id1, Dusp9, Pik3ca, Sox2
Spliceosome
Aqr, Crnkl1, Prpf6, Pqbp1
Proteoglycans in cancer
Fzd7, Vav2, Src, Flna, Msn
RNA transport
Eif5b, Nup188, Nup214, Eif3j1
mTOR signaling pathway
Fzd7, Cab39, Pik3ca, Nras
DNA replication
Mcm3, Prim2, Mcm6, Pola1
Rap1 signaling pathway
Enah, Vav2, Ralgds, Id1
MAPK signaling pathway
Map4k4, Cacnb4, Map3k20, Dusp9
Oocyte meiosis
Ppp2r5a, Smc1a, Rps6ka1
0
10
20
30
40
p.adjust
3e−04
6e−04
9e−04
Cluster 4
Ribosome
Rpl7, Mrpl30, Rpl31, Rpl37a
Proteasome
Psmb7, Psmd4, Psmc2
Parkinson disease
Ndufb3, Atp5c1, Psmb7
Amino sugar and nucleotide sugar metabolism
Gmppa, Renbp, Nans, Pgm2
Biosynthesis of amino acids
Pycr2, Idh3b, Phgdh, Aco1, Eno1
Carbon metabolism
Idh3b, Phgdh, Adh5, Aco1, Eno1
Fluid shear stress and atherosclerosis
Il1a, Il1b, Thbd, Sdc4
Fructose and mannose metabolism
Gmppa, Akr1b3, Akr1b8
DNA replication
Pcna, Rpa2, Rfc5
RNA polymerase
Polr3gl, Polr1e, Polr1d
0
20
40
60
p.adjust
0.0005
0.0010
0.0015
0.0020

### fig. S17

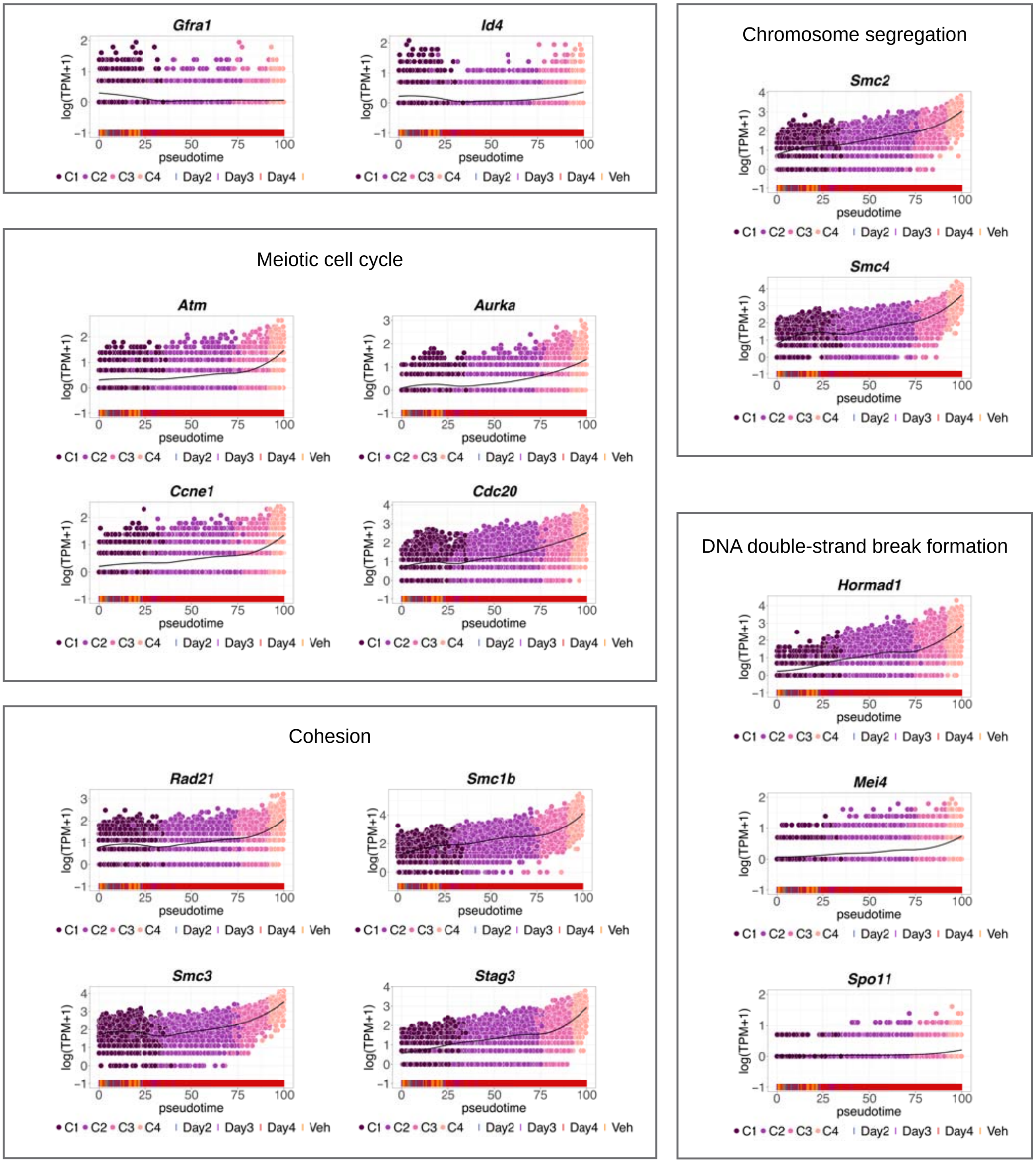

Gfra1
Id4
Chromosome segregation
Smc2
Smc4
Meiotic cell cycle
Atm
Aurka
Ccne1
Cdc20
DNA double-strand break formation
Hormad1
Mei4
Spo11
Cohesion
Rad21
Smc1b
Smc3
Stag3
log(TPM+1)
pseudotime
C1 C2 C3 C4 Day2 Day3 Day4 Veh

#### fig. S18

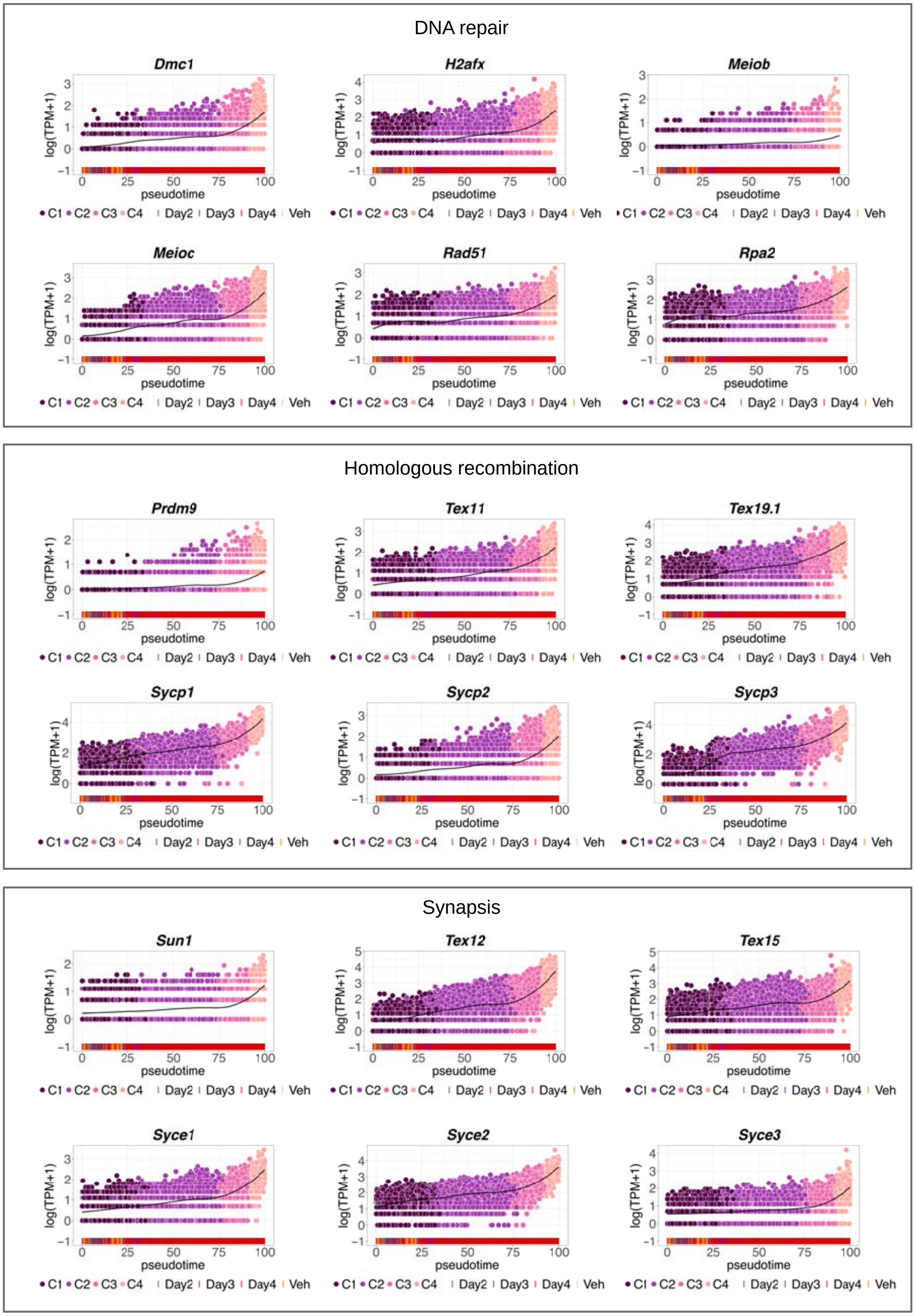
DNA repair
Dmc1
H2afx
Meiob
Meioc
Rad51
Rpa2
Homologous recombination
Prdm9
Tex11
Tex19.1
Sycp1
Sycp2
Sycp3
Synapsis
Sun1
Tex12
Tex15
Syce1
Syce2
Syce3
log(TPM+1)
pseudotime
C1 C2 C3 C4 Day2 Day3 Day4 Veh

### fig. S19

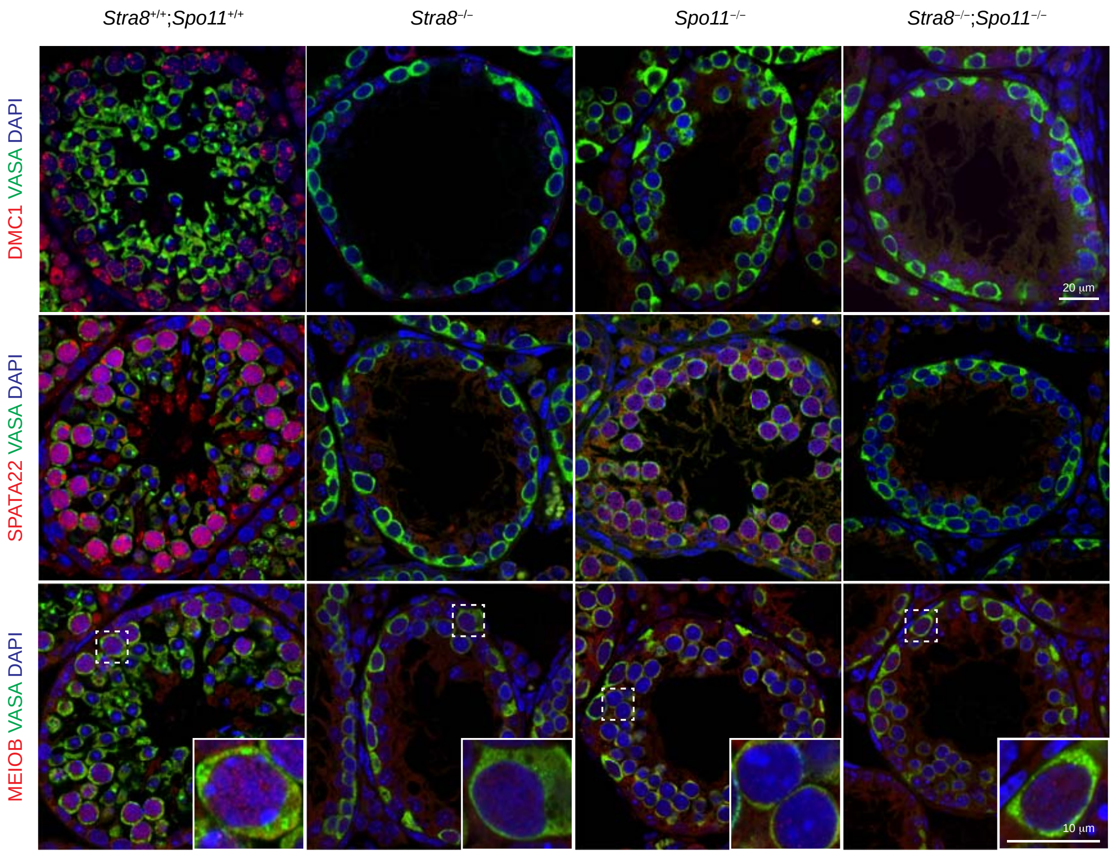

Stra8+/+;Spo11+/+
Stra8–/–
Spo11–/–
Stra8–/–;Spo11–/–
DMC1 VASA DAPI
SPATA22 VASA DAPI
MEIOB VASA DAPI
20 µm
10 µm

fig. S20

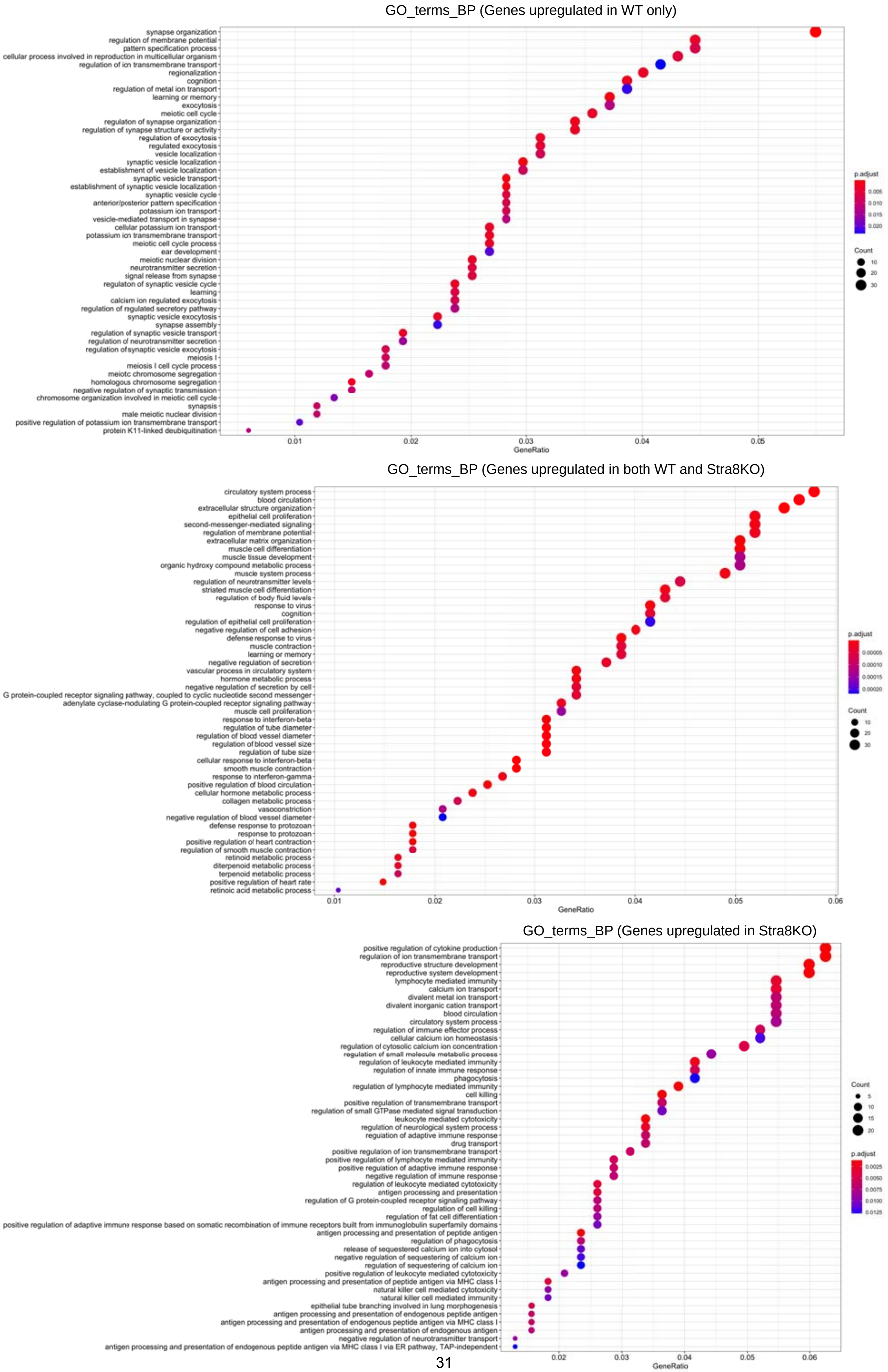

GO_terms_BP (Genes upregulated in WT only)
synapse organization
regulation of membrane potential
pattern specification process
cellular process involved in reproduction in multicellular organism
regulation of ion transmembrane transport
regionalization
cognition
regulation of metal ion transport
learning or memory
exocytosis
meiotic cell cycle
regulation of synapse organization
regulation of synapse structure or activity
regulation of exocytosis
regulated exocytosis
vesicle localization
synaptic vesicle localization
establishment of vesicle localization
synaptic vesicle transport
establishment of synaptic vesicle localization
synaptic vesicle cycle
anterior/posterior pattern specification
potassium ion transport
vesicle-mediated transport in synapse
cellular potassium ion transport
potassium ion transmembrane transport
meiotic cell cycle process
ear development
meiotic nuclear division
neurotransmitter secretion
signal release from synapse
regulation of synaptic vesicle cycle
learning
calcium ion regulated exocytosis
regulation of regulated secretory pathway
synaptic vesicle exocytosis
synapse assembly
regulation of synaptic vesicle transport
regulation of neurotransmitter secretion
regulation of synaptic vesicle exocytosis
meiosis I
meiosis I cell cycle process
meiotic chromosome segregation
homologous chromosome segregation
negative regulation of synaptic transmission
chromosome organization involved in meiotic cell cycle
synapsis
male meiotic nuclear division
positive regulation of potassium ion transmembrane transport
protein K11-linked deubiquitination
0.01
0.02
0.03
0.04
0.05
GeneRatio
p.adjust
0.005
0.010
0.015
0.020
Count
10
20
30
GO_terms_BP (Genes upregulated in both WT and Stra8KO)
circulatory system process
blood circulation
extracellular structure organization
epithelial cell proliferation
second-messenger-mediated signaling
regulation of membrane potential
extracellular matrix organization
muscle cell differentiation
muscle tissue development
organic hydroxy compound metabolic process
muscle system process
regulation of neurotransmitter levels
striated muscle cell differentiation
regulation of body fluid levels
response to virus
cognition
regulation of epithelial cell proliferation
negative regulation of cell adhesion
defense response to virus
muscle contraction
learning or memory
negative regulation of secretion
vascular process in circulatory system
hormone metabolic process
negative regulation of secretion by cell
G protein-coupled receptor signaling pathway, coupled to cyclic nucleotide second messenger
adenylate cyclase-modulating G protein-coupled receptor signaling pathway
muscle cell proliferation
response to interferon-beta
regulation of tube diameter
regulation of blood vessel diameter
regulation of blood vessel size
regulation of tube size
cellular response to interferon-beta
smooth muscle contraction
response to interferon-gamma
positive regulation of blood circulation
cellular hormone metabolic process
collagen metabolic process
vasoconstriction
negative regulation of blood vessel diameter
defense response to protozoan
response to protozoan
positive regulation of heart contraction
regulation of smooth muscle contraction
retinoid metabolic process
diterpenoid metabolic process
terpenoid metabolic process
positive regulation of heart rate
retinoic acid metabolic process
0.01
0.02
0.03
0.04
0.05
0.06
GeneRatio
p.adjust
0.00005
0.00010
0.00015
0.00020
Count
10
20
30
GO_terms_BP (Genes upregulated in Stra8KO)
positive regulation of cytokine production
regulation of ion transmembrane transport
reproductive structure development
reproductive system development
lymphocyte mediated immunity
calcium ion transport
divalent metal ion transport
divalent inorganic cation transport
blood circulation
circulatory system process
regulation of immune effector process
cellular calcium ion homeostasis
regulation of cytosolic calcium ion concentration
regulation of small molecule metabolic process
regulation of leukocyte mediated immunity
regulation of innate immune response
phagocytosis
regulation of lymphocyte mediated immunity
cell killing
positive regulation of transmembrane transport
regulation of small GTPase mediated signal transduction
leukocyte mediated cytotoxicity
regulation of neurological system process
regulation of adaptive immune response
drug transport
positive regulation of ion transmembrane transport
positive regulation of lymphocyte mediated immunity
positive regulation of adaptive immune response
negative regulation of immune response
regulation of leukocyte mediated cytotoxicity
antigen processing and presentation
regulation of G protein-coupled receptor signaling pathway
regulation of cell killing
regulation of fat cell differentiation
positive regulation of adaptive immune response based on somatic recombination of immune receptors built from immunoglobulin superfamily domains
antigen processing and presentation of peptide antigen
regulation of phagocytosis
release of sequestered calcium ion into cytosol
negative regulation of sequestering of calcium ion
regulation of sequestering of calcium ion
positive regulation of leukocyte mediated cytotoxicity
antigen processing and presentation of peptide antigen via MHC class I
natural killer cell mediated cytotoxicity
natural killer cell mediated immunity
epithelial tube branching involved in lung morphogenesis
antigen processing and presentation of endogenous peptide antigen
antigen processing and presentation of endogenous peptide antigen via MHC class I
antigen processing and presentation of endogenous antigen
negative regulation of neurotransmitter transport
antigen processing and presentation of endogenous peptide antigen via MHC class I via ER pathway, TAP-independent
0.02
0.03
0.04
0.05
0.06
GeneRatio
Count
5
10
15
20
p.adjust
0.0025
0.0050
0.0075
0.0100
0.0125

fig. S21

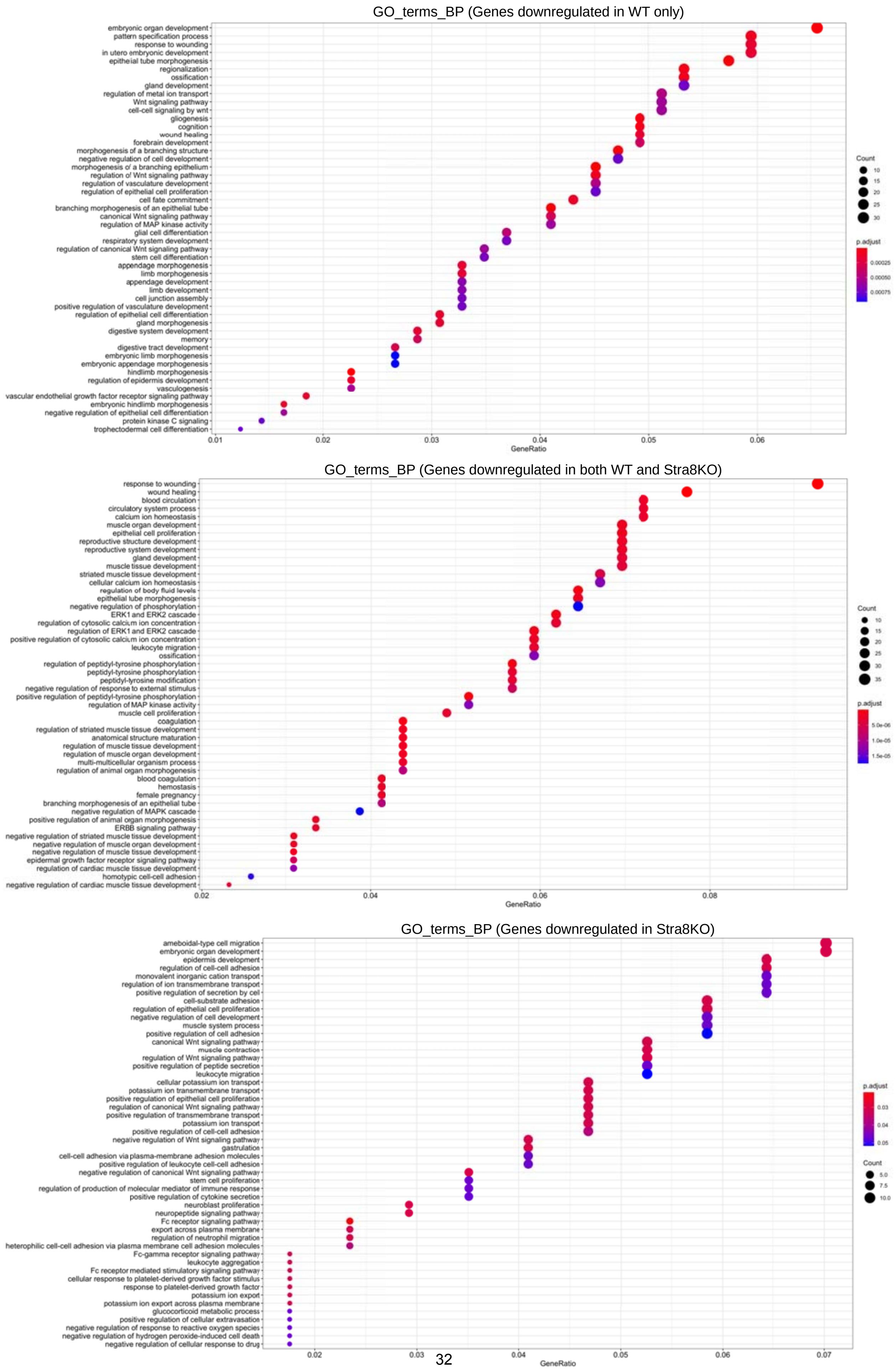

GO_terms_BP (Genes downregulated in WT only)
embryonic organ development
pattern specification process
response to wounding
in utero embryonic development
epithelial tube morphogenesis
regionalization
ossification
gland development
regulation of metal ion transport
Wnt signaling pathway
cell-cell signaling by wnt
gliogenesis
cognition
wound healing
forebrain development
morphogenesis of a branching structure
negative regulation of cell development
morphogenesis of a branching epithelium
regulation of Wnt signaling pathway
regulation of vasculature development
regulation of epithelial cell proliferation
cell fate commitment
branching morphogenesis of an epithelial tube
canonical Wnt signaling pathway
regulation of MAP kinase activity
glial cell differentiation
respiratory system development
regulation of canonical Wnt signaling pathway
stem cell differentiation
appendage morphogenesis
limb morphogenesis
appendage development
limb development
cell junction assembly
positive regulation of vasculature development
regulation of epithelial cell differentiation
gland morphogenesis
digestive system development
memory
digestive tract development
embryonic limb morphogenesis
embryonic appendage morphogenesis
hindlimb morphogenesis
regulation of epidermis development
vasculogenesis
vascular endothelial growth factor receptor signaling pathway
embryonic hindlimb morphogenesis
negative regulation of epithelial cell differentiation
protein kinase C signaling
trophectodermal cell differentiation
0.01
0.02
0.03
0.04
0.05
0.06
GeneRatio
Count
10
15
20
25
30
p.adjust
0.00025
0.00050
0.00075
GO_terms_BP (Genes downregulated in both WT and Stra8KO)
response to wounding
wound healing
blood circulation
circulatory system process
calcium ion homeostasis
muscle organ development
epithelial cell proliferation
reproductive structure development
reproductive system development
gland development
muscle tissue development
striated muscle tissue development
cellular calcium ion homeostasis
regulation of body fluid levels
epithelial tube morphogenesis
negative regulation of phosphorylation
ERK1 and ERK2 cascade
regulation of cytosolic calcium ion concentration
regulation of ERK1 and ERK2 cascade
positive regulation of cytosolic calcium ion concentration
leukocyte migration
ossification
regulation of peptidyl-tyrosine phosphorylation
peptidyl-tyrosine phosphorylation
peptidyl-tyrosine modification
negative regulation of response to external stimulus
positive regulation of peptidyl-tyrosine phosphorylation
regulation of MAP kinase activity
muscle cell proliferation
coagulation
regulation of striated muscle tissue development
anatomical structure maturation
regulation of muscle tissue development
regulation of muscle organ development
multi-multicellular organism process
regulation of animal organ morphogenesis
blood coagulation
hemostasis
female pregnancy
branching morphogenesis of an epithelial tube
negative regulation of MAPK cascade
positive regulation of animal organ morphogenesis
ERBB signaling pathway
negative regulation of striated muscle tissue development
negative regulation of muscle organ development
negative regulation of muscle tissue development
epidermal growth factor receptor signaling pathway
regulation of cardiac muscle tissue development
homotypic cell-cell adhesion
negative regulation of cardiac muscle tissue development
0.02
0.04
0.06
0.08
GeneRatio
Count
10
15
20
25
30
35
p.adjust
5.0e-06
1.0e-05
1.5e-05
GO_terms_BP (Genes downregulated in Stra8KO)
ameboidal-type cell migration
embryonic organ development
epidermis development
regulation of cell-cell adhesion
monovalent inorganic cation transport
regulation of ion transmembrane transport
positive regulation of secretion by cell
cell-substrate adhesion
regulation of epithelial cell proliferation
negative regulation of cell development
muscle system process
positive regulation of cell adhesion
canonical Wnt signaling pathway
muscle contraction
regulation of Wnt signaling pathway
positive regulation of peptide secretion
leukocyte migration
cellular potassium ion transport
potassium ion transmembrane transport
positive regulation of epithelial cell proliferation
regulation of canonical Wnt signaling pathway
positive regulation of transmembrane transport
potassium ion transport
positive regulation of cell-cell adhesion
negative regulation of Wnt signaling pathway
gastrulation
cell-cell adhesion via plasma-membrane adhesion molecules
positive regulation of leukocyte cell-cell adhesion
negative regulation of canonical Wnt signaling pathway
stem cell proliferation
regulation of production of molecular mediator of immune response
positive regulation of cytokine secretion
neuroblast proliferation
neuropeptide signaling pathway
Fc receptor signaling pathway
export across plasma membrane
regulation of neutrophil migration
heterophilic cell-cell adhesion via plasma membrane cell adhesion molecules
Fc-gamma receptor signaling pathway
leukocyte aggregation
Fc receptor mediated stimulatory signaling pathway
cellular response to platelet-derived growth factor stimulus
response to platelet-derived growth factor
potassium ion export
potassium ion export across plasma membrane
glucocorticoid metabolic process
positive regulation of cellular extravasation
negative regulation of response to reactive oxygen species
negative regulation of hydrogen peroxide-induced cell death
negative regulation of cellular response to drug
0.02
0.03
0.04
0.05
0.06
0.07
GeneRatio
p.adjust
0.03
0.04
0.05
Count
5.0
7.5
10.0

### fig. S22

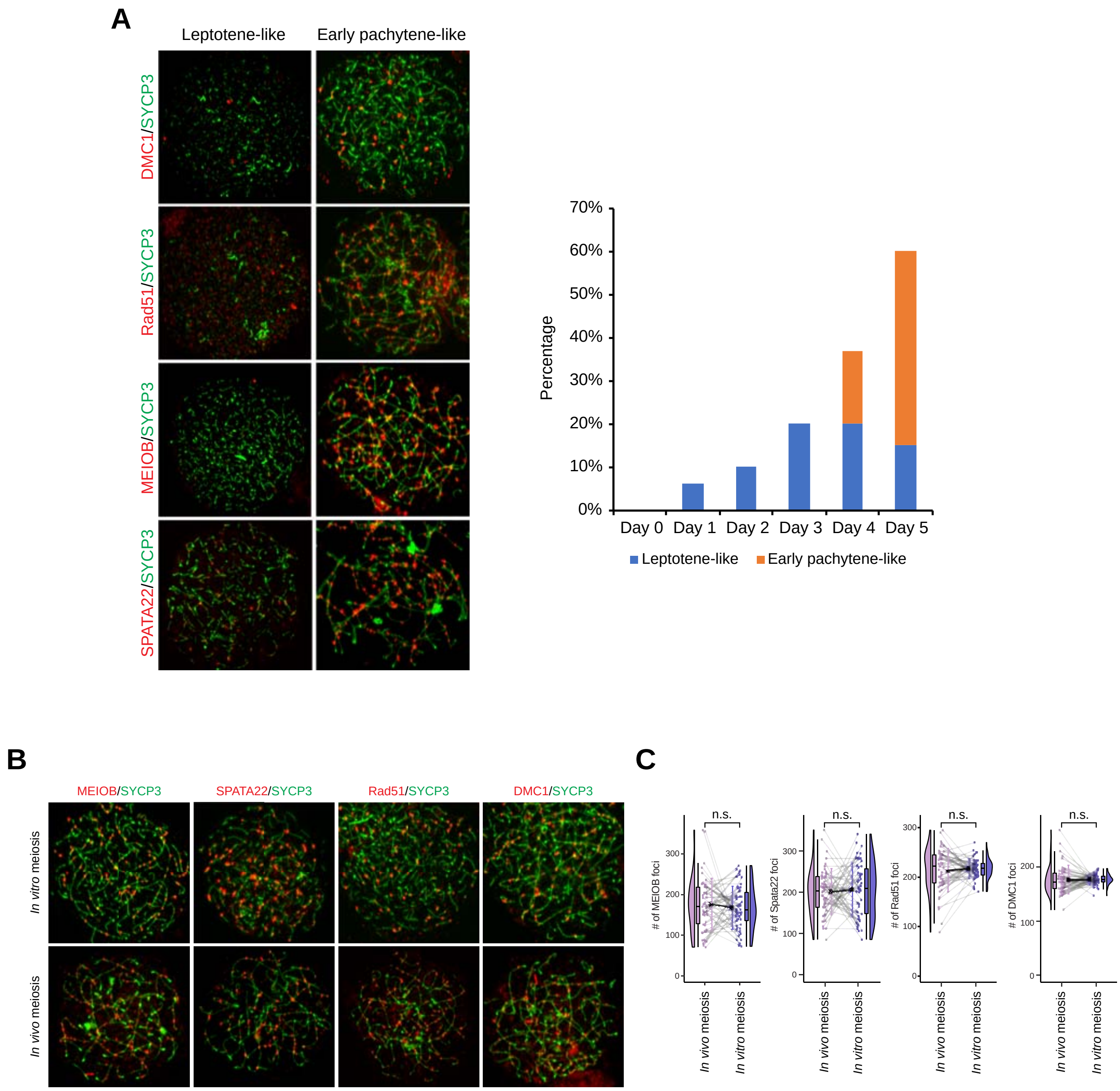
A
Leptotene-like
Early pachytene-like
DMC1/SYCP3
Rad51/SYCP3
MEIOB/SYCP3
SPATA22/SYCP3
70%
60%
50%
40%
30%
20%
10%
0%
Percentage
Day 0
Day 1
Day 2
Day 3
Day 4
Day 5
Leptotene-like
Early pachytene-like
B
MEIOB/SYCP3
SPATA22/SYCP3
Rad51/SYCP3
DMC1/SYCP3
In vitro meiosis
In vivo meiosis
C
n.s.
### of MEIOB foci
### of Spata22 foci
### of Rad51 foci
### of DMC1 foci
300
200
100
0
In vivo meiosis
In vitro meiosis

### fig. S23

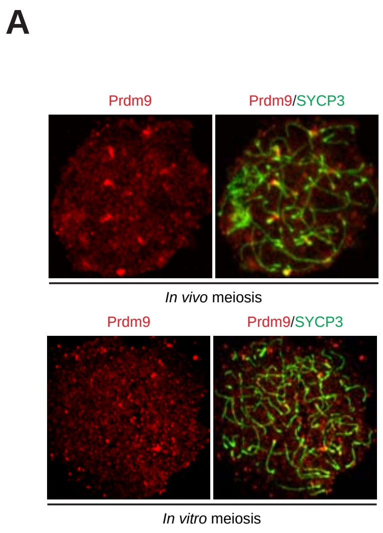
A
Prdm9
Prdm9/SYCP3
In vivo meiosis
Prdm9
Prdm9/SYCP3
In vitro meiosis

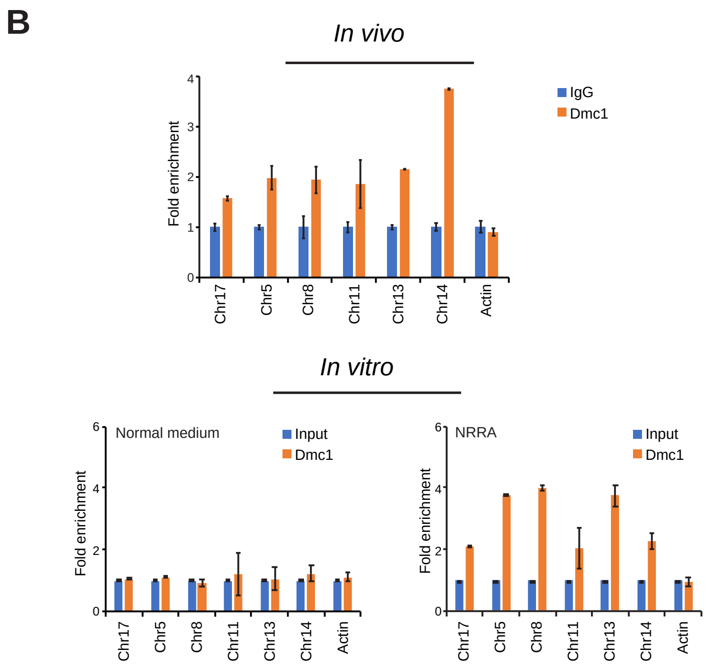
B
In vivo
Fold enrichment
0
1
2
3
4
IgG
Dmc1
Chr17
Chr5
Chr8
Chr11
Chr13
Chr14
Actin
In vitro
Normal medium
Input
Dmc1
Fold enrichment
0
2
4
6
Chr17
Chr5
Chr8
Chr11
Chr13
Chr14
Actin
NRRA
Input
Dmc1
Fold enrichment
0
2
4
6
Chr17
Chr5
Chr8
Chr11
Chr13
Chr14
Actin

### fig. S24

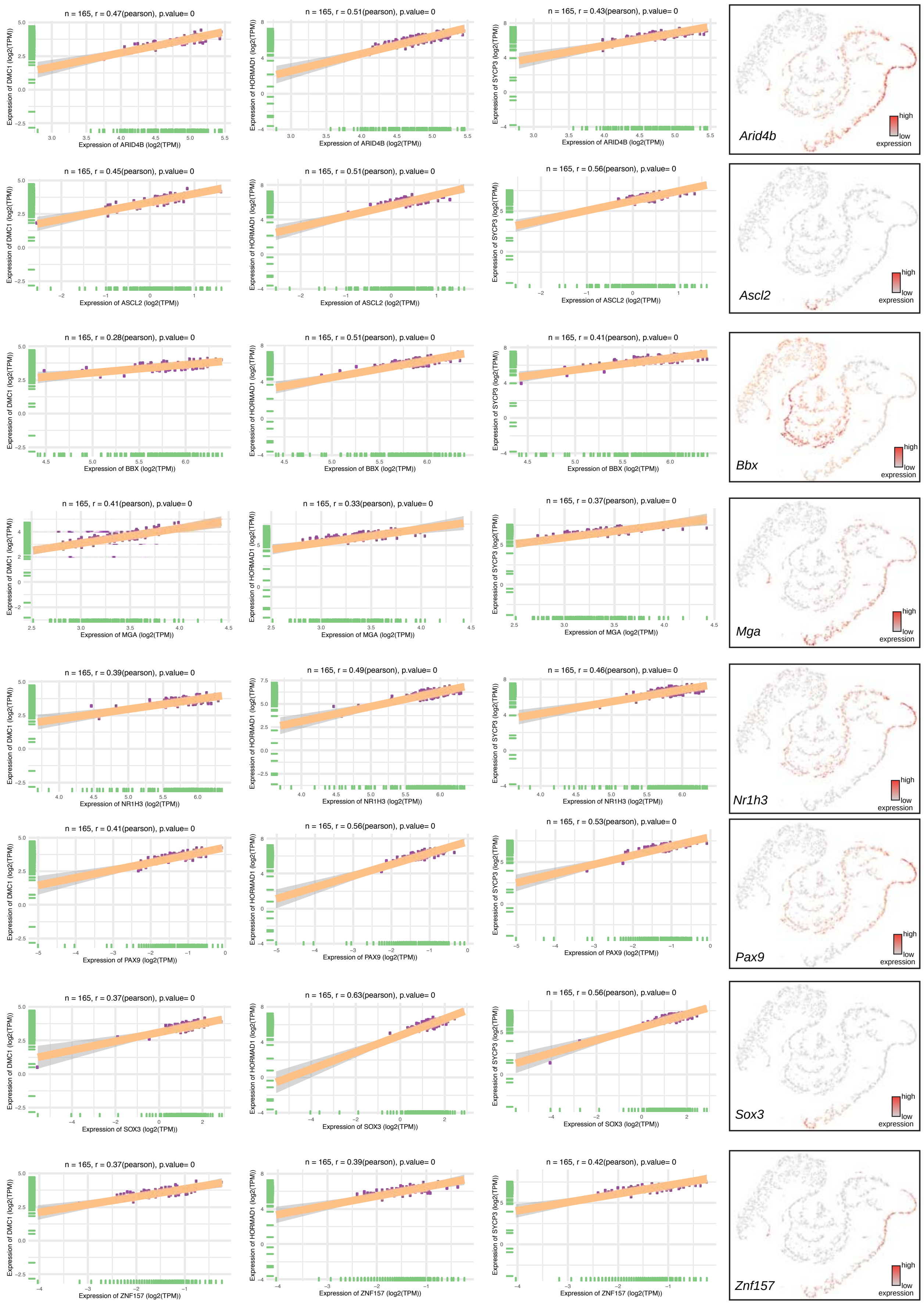

n = 165, r = 0.47(pearson), p.value= 0
n = 165, r = 0.51(pearson), p.value= 0
n = 165, r = 0.43(pearson), p.value= 0
Expression of ARID4B (log2(TPM))
n = 165, r = 0.45(pearson), p.value= 0
n = 165, r = 0.51(pearson), p.value= 0
n = 165, r = 0.56(pearson), p.value= 0
Expression of ASCL2 (log2(TPM))
n = 165, r = 0.28(pearson), p.value= 0
n = 165, r = 0.51(pearson), p.value= 0
n = 165, r = 0.41(pearson), p.value= 0
Expression of BBX (log2(TPM))
n = 165, r = 0.41(pearson), p.value= 0
n = 165, r = 0.33(pearson), p.value= 0
n = 165, r = 0.37(pearson), p.value= 0
Expression of MGA (log2(TPM))
n = 165, r = 0.39(pearson), p.value= 0
n = 165, r = 0.49(pearson), p.value= 0
n = 165, r = 0.46(pearson), p.value= 0
Expression of NR1H3 (log2(TPM))
n = 165, r = 0.41(pearson), p.value= 0
n = 165, r = 0.56(pearson), p.value= 0
n = 165, r = 0.53(pearson), p.value= 0
Expression of PAX9 (log2(TPM))
n = 165, r = 0.37(pearson), p.value= 0
n = 165, r = 0.63(pearson), p.value= 0
n = 165, r = 0.56(pearson), p.value= 0
Expression of SOX3 (log2(TPM))
n = 165, r = 0.37(pearson), p.value= 0
n = 165, r = 0.39(pearson), p.value= 0
n = 165, r = 0.42(pearson), p.value= 0
Expression of ZNF157 (log2(TPM))
Expression of DMC1 (log2(TPM))
Expression of HORMAD1 (log2(TPM))
Expression of SYCP3 (log2(TPM))
Arid4b
Ascl2
Bbx
Mga
Nr1h3
Pax9
Sox3
Znf157
high
low
expression

#### fig. S25

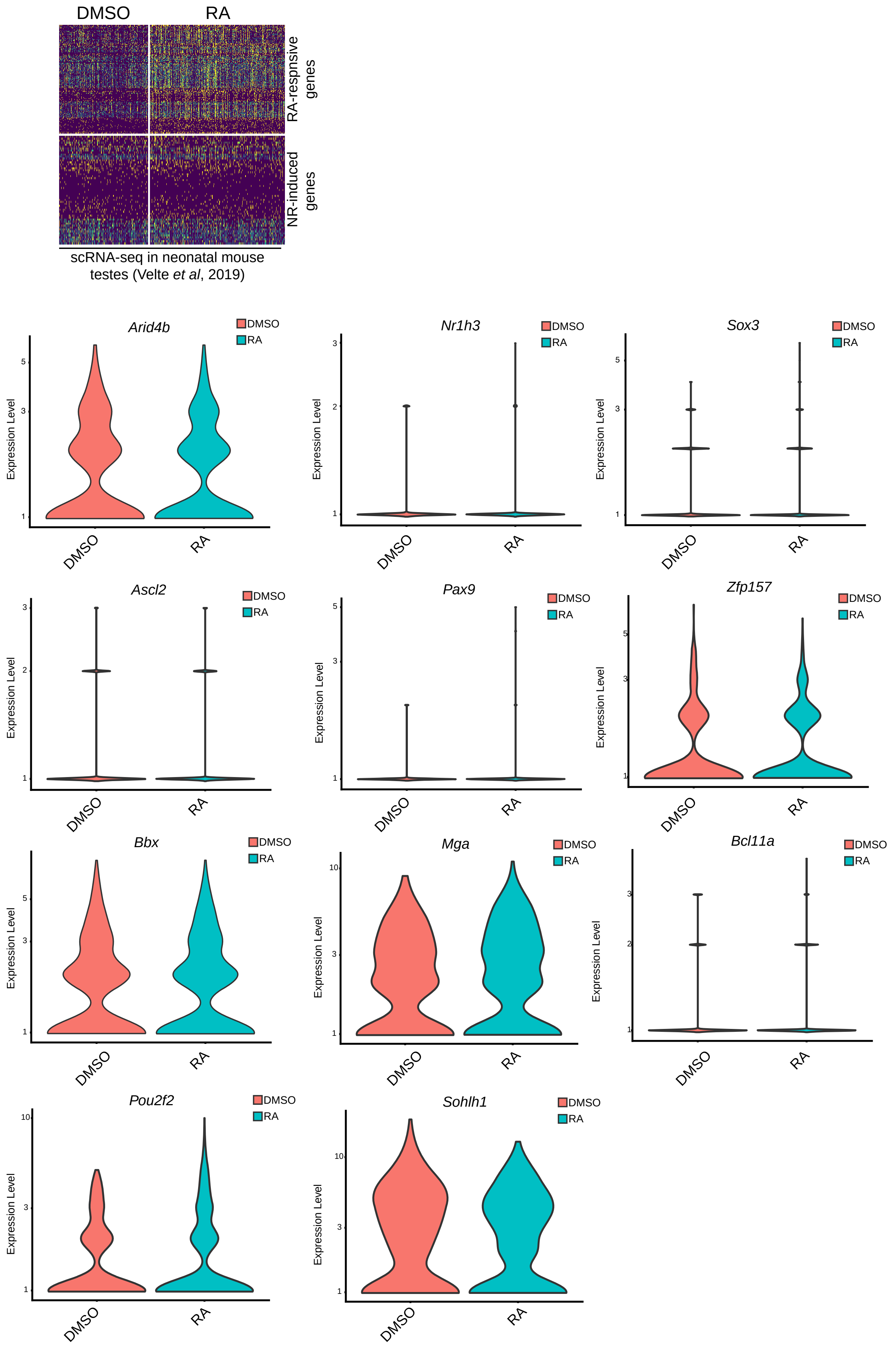
DMSO
RA
RA-respnsive genes
NR-induced genes
scRNA-seq in neonatal mouse testes (Velte *et al*, 2019)
Arid4b
Nr1h3
Sox3
Ascl2
Pax9
Zfp157
Bbx
Mga
Bcl11a
Pou2f2
Sohlh1
Expression Level
DMSO
RA

**fig. S26**

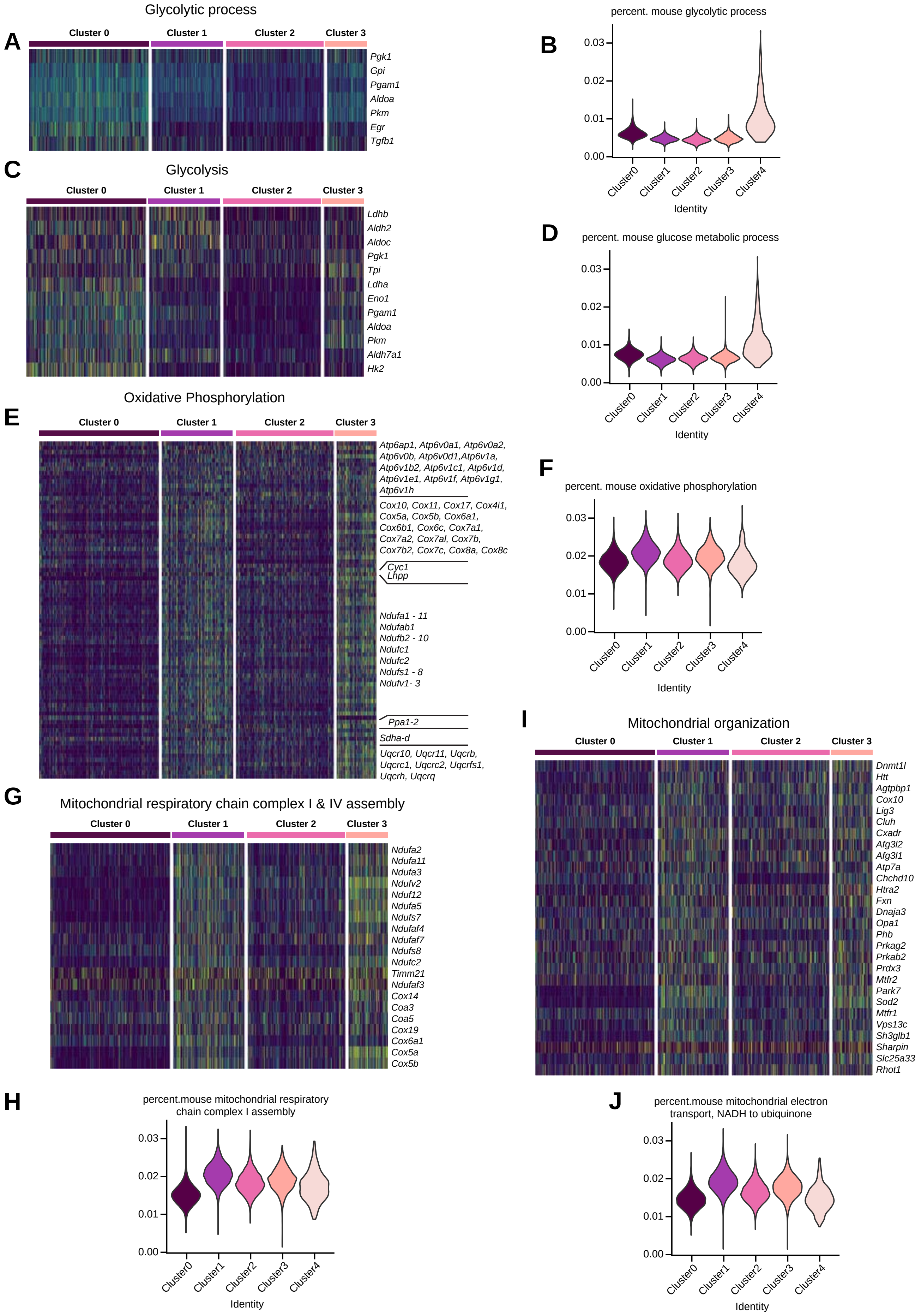

A
Glycolytic process
Cluster 0
Cluster 1
Cluster 2
Cluster 3
Pgk1
Gpi
Pgam1
Aldoa
Pkm
Egr
Tgfb1
B
percent. mouse glycolytic process
0.03
0.02
0.01
0.00
Cluster0
Cluster1
Cluster2
Cluster3
Cluster4
Identity
C
Glycolysis
Ldhb
Aldh2
Aldoc
Pgk1
Tpi
Ldha
Eno1
Pgam1
Aldoa
Pkm
Aldh7a1
Hk2
D
percent. mouse glucose metabolic process
E
Oxidative Phosphorylation
Atp6ap1, Atp6v0a1, Atp6v0a2, Atp6v0b, Atp6v0d1,Atp6v1a, Atp6v1b2, Atp6v1c1, Atp6v1d, Atp6v1e1, Atp6v1f, Atp6v1g1, Atp6v1h
Cox10, Cox11, Cox17, Cox4i1, Cox5a, Cox5b, Cox6a1, Cox6b1, Cox6c, Cox7a1, Cox7a2, Cox7al, Cox7b, Cox7b2, Cox7c, Cox8a, Cox8c
Cyc1
Lhpp
Ndufa1 - 11
Ndufab1
Ndufb2 - 10
Ndufc1
Ndufc2
Ndufs1 - 8
Ndufv1- 3
Ppa1-2
Sdha-d
Uqcr10, Uqcr11, Uqcrb, Uqcrc1, Uqcrc2, Uqcrfs1, Uqcrh, Uqcrq
F
percent. mouse oxidative phosphorylation
G
Mitochondrial respiratory chain complex I & IV assembly
Ndufa2
Ndufa11
Ndufa3
Ndufv2
Nduf12
Ndufa5
Ndufs7
Ndufaf4
Ndufaf7
Ndufs8
Ndufc2
Timm21
Ndufaf3
Cox14
Coa3
Coa5
Cox19
Cox6a1
Cox5a
Cox5b
H
percent.mouse mitochondrial respiratory chain complex I assembly
I
Mitochondrial organization
Dnmt1l
Htt
Agtpbp1
Cox10
Lig3
Cluh
Cxadr
Afg3l2
Afg3l1
Atp7a
Chchd10
Htra2
Fxn
Dnaja3
Opa1
Phb
Prkag2
Prkab2
Prdx3
Mtfr2
Park7
Sod2
Mtfr1
Vps13c
Sh3glb1
Sharpin
Slc25a33
Rhot1
J
percent.mouse mitochondrial electron transport, NADH to ubiquinone
